# A Biodiversity Composition Map of California Derived from Environmental DNA Metabarcoding and Earth Observation

**DOI:** 10.1101/2020.06.19.160374

**Authors:** Meixi Lin, Ariel Levi Simons, Emily E. Curd, Ryan J. Harrigan, Fabian D. Schneider, Dannise V. Ruiz-Ramos, Zack Gold, Melisa G. Osborne, Sabrina Shirazi, Teia M. Schweizer, Tiara N. Moore, Emma A. Fox, Rachel Turba, Ana E. Garcia-Vedrenne, Sarah K. Helman, Kelsi Rutledge, Maura Palacios Mejia, Miroslava N. Munguia Ramos, Regina Wetzer, Dean Pentcheff, Emily Jane McTavish, Michael N. Dawson, Beth Shapiro, Robert K. Wayne, Rachel S. Meyer

## Abstract

Unique ecosystems globally are under threat from ongoing anthropogenic environmental change. Effective conservation management requires more thorough biodiversity surveys that can reveal system-level patterns and that can be applied rapidly across space and time. We offer a way to use environmental DNA, community science and remote sensing together as methods to reduce the discrepancy between the magnitude of change and historical approaches to measure it. Taking advantages of modern ecological models, we integrate environmental DNA and Earth observations to evaluate regional biodiversity patterns for a snapshot of time, and provide critical community-level characterization. We collected 278 samples in Spring 2017 from coastal, shrub and lowland forest sites in California, a large-scale biodiversity hotspot. We applied gradient forest to model 915 family occurrences and community composition together with environmental variables and multi-scalar habitat classifications to produce a statewide biodiversity-based map. 16,118 taxonomic entries recovered were associated with environmental variables to test their predictive strength on alpha, beta, and zeta diversity. Local habitat classification was diagnostic of community composition, illuminating a characteristic of biodiversity hotspots. Using gradient forest models, environmental variables predicted 35% of the variance in eDNA patterns at the family level, with elevation, sand percentage, and greenness (NDVI32) as the top predictors. This predictive power was higher than we found in published literature at global scale. In addition to this indication of substantial environmental filtering, we also found a positive relationship between environmentally predicted families and their numbers of biotic interactions. In aggregate, these analyses showed that strong eDNA community-environment correlation is a general characteristic of temperate ecosystems, and may explain why communities easily destabilize under disturbances. Our study provides the first example of integrating citizen science based eDNA with biodiversity mapping across the tree of life, with promises to produce large scale, high resolution assessments that promote a more comprehensive and predictive understanding of the factors that influence biodiversity and enhance its maintenance.

## Introduction

Global biodiversity is currently undergoing rapid loss (Pimm et al. 2014, Ceballos et al. 2015, Díaz et al. 2019) with many species and geographic areas (Myers et al. 2000) threatened by unique environmental challenges such as climate change and habitat degradation. The scientific community needs better tools to provide critical baseline biodiversity data that can be applied rapidly with minimal cost and effort (Bush et al. 2017). A recently discussed goal of biodiversity conservation is to estimate the Essential Biodiversity Variables (Pereira et al. 2013) that are a minimal set of fundamental observations needed to support multi-purpose, long-term planning at various scales, from genetic variation, to species diversity and ecosystem structure. However, scaling up from *in situ* biological measures remains challenging (Pereira et al. 2013). Changes in the richness and turnover of species are usually tracked in taxonomically or spatiotemporally restricted studies because technical feasibility limits large scale monitoring (Cristescu 2014). Very few studies attempt to assess the complex composition of entire biological communities (Karimi et al. 2018, George et al. 2019).

As one new approach to achieving this need for tracking biodiversity over large scales, the past decade has showcased the growing power of biodiversity inventory by non-experts: the use of technology assisted citizen and community science (CCS). CCS for traditional biomonitoring through observation has already dramatically scaled up biomonitoring efforts (Theobald et al. 2015, Kobori et al. 2016, Waller 2019). However, these approaches favor diurnal macroscopic taxa and often omit cryptic and microbial taxa (Theobald et al. 2015). Recently, programs have launched that arm community scientists with the capacity to sample environmental DNA (eDNA) from their surroundings (Biggs et al. 2015, Miralles et al. 2016, Deiner et al. 2017, Meyer et al. 2019) which can be probed for nearly any taxonomic group using metabarcoding methods (Bohmann et al. 2014, Deiner et al. 2016, Thompson et al. 2017, Taberlet et al. 2018, Franklin et al. 2019). These methods are improving in accuracy as public DNA sequence databases grow, while simultaneously decreasing in cost, suggesting community-powered eDNA surveys may provide the more complete biodiversity data needed for mapping and predicting taxon presence as well as determining community assemblage patterns and the Essential Biodiversity Variables.

Multi-locus metabarcoding of eDNA from surface soil and sediment retains a record of taxa recently present in the local area. The temporal span of the surface DNA record from these substrates may be weeks to months, even several years (Andersen et al. 2012, Parducci et al. 2017). Surface DNA can be easily collected with minimal environmental impact (Meyer et al. 2019) and the taxonomic signatures preserved in eDNA perform similarly to direct observation surveys (Tedersoo et al. 2014, Lejzerowicz et al. 2015) including vertebrate and vascular plant communities that are of conservation concern, but also yield a species catalog that is much broader, including bacteria and archaea, often-overlooked meiofauna, protozoans, non-vascular plants, algae, and fungi.

Efforts to integrate surface eDNA with remote measures of ecosystem properties present an opportunity to improve models and ecological theories (Yamasaki et al. 2017). In parallel with the developments in eDNA metabarcoding on biodiversity monitoring, on the ground and space-based technologies yield increasingly copious and accessible abiotic data (Pettorelli et al. 2014, Schimel et al. 2019) on land cover, topography, soil property (e.g. US Geological Survey), bioclimate (Fick and Hijmans 2017), human impact (WCS and CIESIN 2005) and vegetation (e.g. Sentinel-2; European Space Agency) which can be used to characterize eDNA biodiversity changes across a landscape (van den Hoogen et al. 2019, Crowther et al. 2019). Biotic-abiotic interactions among soil properties (e.g. pH and nutrient availabilities), climate, plant coverage, and habitat type affect soil alpha and beta diversity (Fierer and Jackson 2006, Ranjard et al. 2013, Montagna et al. 2018, George et al. 2019). Different taxonomic groups have differential responses to environmental cues in diverse ecosystems from tropical mountains to temperate ecosystems (Thompson et al. 2017, Karimi et al. 2018, Montagna et al. 2018, Peters et al. 2019). For example, a national-scale soil eDNA survey in England showed that animal and microbial richness responded to different environment factors but beta-diversity trends were shared across taxonomic groups (George et al. 2019). However, biodiversity rich regions and biodiversity hotspots have scarcely been studied in taxonomic biodiversity–ecological response models. Such regions are discontinuous in environmental clines and have high endemism (Myers et al. 2000, Thompson et al. 2017). Surface eDNA signatures could aid predictions of regional biodiversity patterns to guide conservation planning (Bush et al. 2017, Jetz et al. 2019, Breed et al. 2019) and theory development (Konopka 2009, Kurtz et al. 2015, Prosser 2015, Koskella et al. 2017).

To integrate the regional environmental variables for predictions on eDNA biodiversity turnover, joint-species distribution models are needed that can utilize big-data. The gradient forest model is an offshoot of the random forest algorithm (Breiman 2001) that attempts to explain turnover in community assemblages and can be used for regional community turnover predictions (Ferrier and Guisan 2006, Ellis et al. 2012). Briefly, the gradient forest first establishes a composite relationship of species’ response (in this case, eDNA profiles) to environmental predictors, and then projects the turnover of this response across unsampled regions using this relationship (Ellis et al. 2012). The gradient forest is robust, performs well with large number of correlated explanatory variables and captures non-linear responses of community turnover (Fitzpatrick and Keller 2015). It also handles the input of biotic matrix without the need for ordination, thus utilizing all information available in modeling (Ellis et al. 2012, Pitcher et al. 2012). However, such integration of remotely sensed variables with eDNA for biodiversity mapping remains unexplored.

Here, we present an analysis of eDNA-based biodiversity from the California Environmental DNA (CALeDNA; Meyer et al. 2019) program and propose a framework to perform citizen and community science enabled biodiversity mapping using surface eDNA profiles and environmental variables. It is the first test case of an eDNA survey to describe, analyze and predict the biotic composition of a large-scale biodiversity hotspot, the state of California (CA). Launched in 2017, CALeDNA volunteers have contributed over 5000 georeferenced surface samples and have participated in the evaluation of presence patterns through online tools (www.ucedna.com). Results from 278 samples collected in one seasonal window largely from natural reserves are presented here and integrated multiple Earth observations in diversity analyses and models to explain habitat diversity and community turnover. Our research demonstrates how large-scale surface collections can be used for routine mapping of biodiversity, which may help track the resilience of species and community responses to environmental fluctuations and anthropogenic threats.

## Methods

### Sampling design

We aimed to sample biodiversity from a wide variety of habitats across the state of California using directed volunteers and eDNA metabarcoding. CALeDNA scientists recruited volunteers to perform sampling, and were guided through virtual and in-person training to select soil and sediment sites from natural areas where permission was granted (Supplemental Methods). Sampling steps were guided by a smartphone webform (Meyer et al. 2019) made in Kobo Toolbox (kobotoolbox.org). Surface samples were collected by filling 2 mL tubes with substrate from < 2 cm depth in three biological replicates, each 30 cm apart. Volunteers changed gloves between sampling sites. Samples were frozen at −80 °C immediately upon their return to CALeDNA headquarters at UC Los Angeles.

To minimize the potential effect of seasonal variations in eDNA profiles, we selected samples from March 2017 to July 2017. We classified the predominant biome using photographs and a variety of geolocation data (Supplemental Methods; Table 1; Table S1). We selected 100 samples from each of three *transect* types which were coast, shrub/scrub (abbreviated as “shrub”), and forest that covered the broadest latitudinal range possible. Samples with ambiguous metadata were removed, resulting in a total of 278 samples (98 coast, 89 shrub and 91 forest) used in subsequent analyses.

**Table 1.**
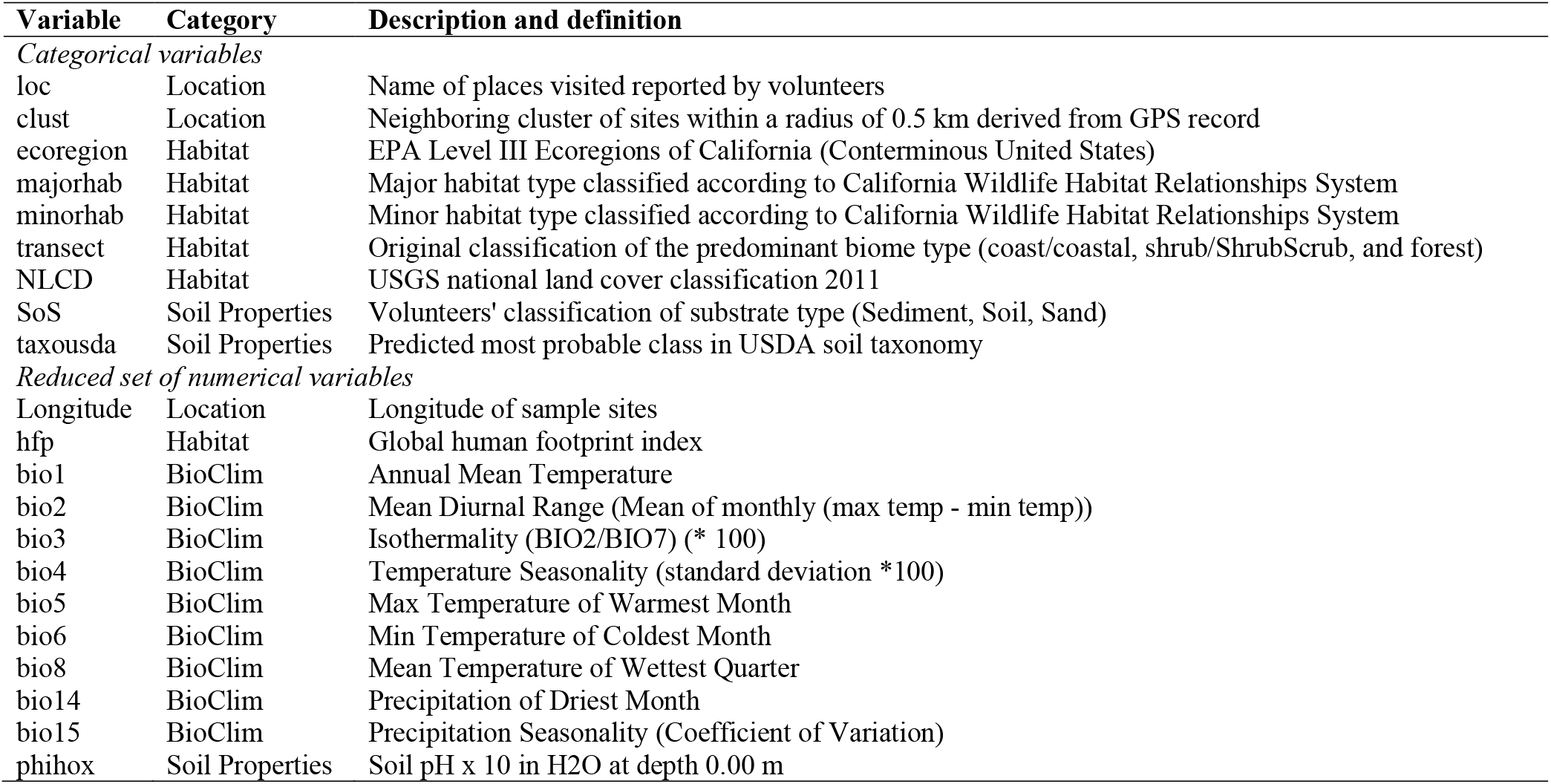
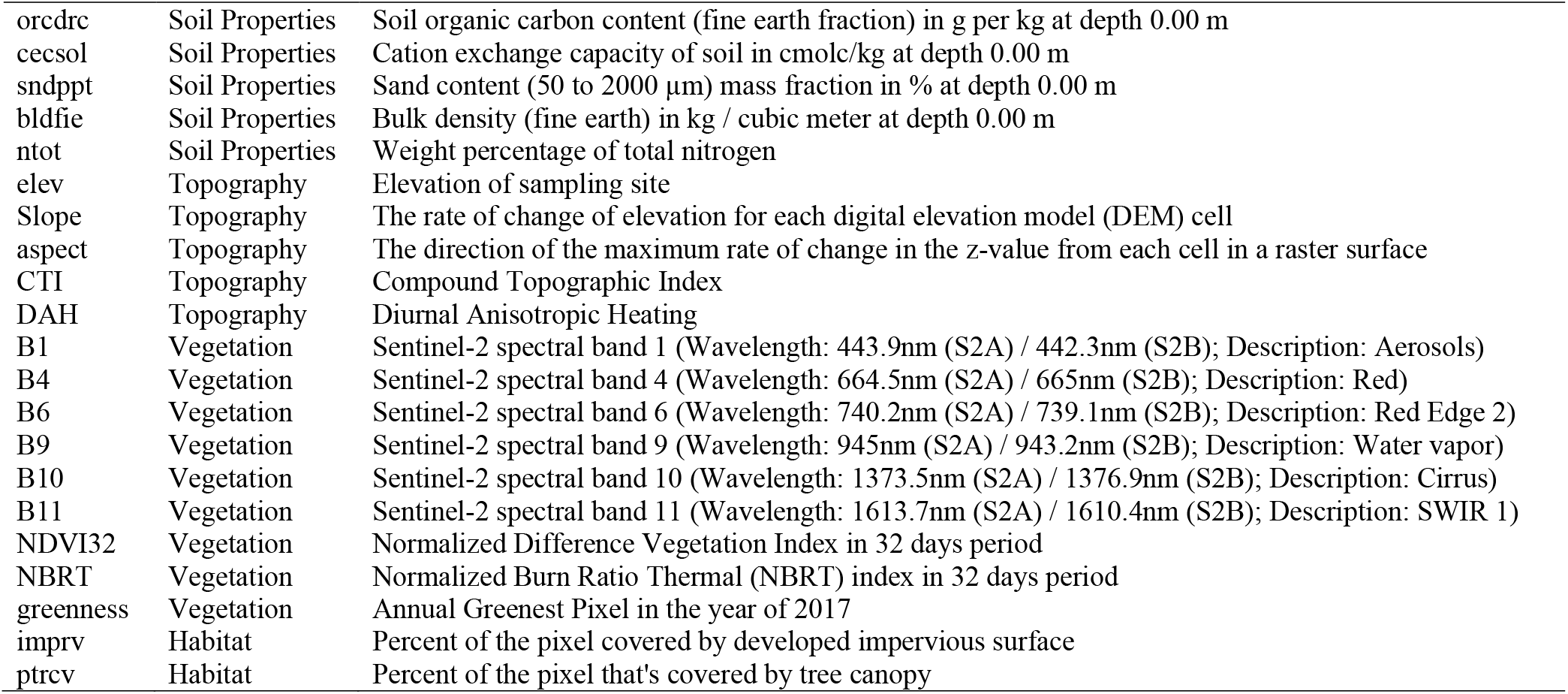
List of the categorical and a reduced set of numerical variables used in the diversity analysis and gradient forest modeling. For a complete list of variables, detailed description and data URL, refer to Table S1.

| Variable | Category | Description and definition |
| --- | --- | --- |
| *Categorical variables* |  |  |
| loc | Location | Name of places visited reported by volunteers |
| clust | Location | Neighboring cluster of sites within a radius of 0.5 km derived from GPS record |
| ecoregion | Habitat | EPA Level III Ecoregions of California (Conterminous United States) |
| majorhab | Habitat | Major habitat type classified according to California Wildlife Habitat Relationships System |
| minorhab | Habitat | Minor habitat type classified according to California Wildlife Habitat Relationships System |
| transect | Habitat | Original classification of the predominant biome type (coast/coastal, shrub/ShrubScrub, and forest) |
| NLCD | Habitat | USGS national land cover classification 2011 |
| SoS | Soil Properties | Volunteers' classification of substrate type (Sediment, Soil, Sand) |
| taxousda | Soil Properties | Predicted most probable class in USDA soil taxonomy |
| *Reduced set of numerical variables* |  |  |
| Longitude | Location | Longitude of sample sites |
| hfp | Habitat | Global human footprint index |
| bio1 | BioClim | Annual Mean Temperature |
| bio2 | BioClim | Mean Diurnal Range (Mean of monthly (max temp - min temp)) |
| bio3 | BioClim | Isothermality (BIO2/BIO7) (* 100) |
| bio4 | BioClim | Temperature Seasonality (standard deviation *100) |
| bio5 | BioClim | Max Temperature of Warmest Month |
| bio6 | BioClim | Min Temperature of Coldest Month |
| bio8 | BioClim | Mean Temperature of Wettest Quarter |
| bio14 | BioClim | Precipitation of Driest Month |
| bio15 | BioClim | Precipitation Seasonality (Coefficient of Variation) |
| phihox | Soil Properties | Soil pH x 10 in H2O at depth 0.00 m |
| orcdrc | Soil Properties | Soil organic carbon content (fine earth fraction) in g per kg at depth 0.00 m |
| cecsol | Soil Properties | Cation exchange capacity of soil in cmolc/kg at depth 0.00 m |
| sndppt | Soil Properties | Sand content (50 to 2000 µm) mass fraction in % at depth 0.00 m |
| bldfie | Soil Properties | Bulk density (fine earth) in kg / cubic meter at depth 0.00 m |
| ntot | Soil Properties | Weight percentage of total nitrogen |
| elev | Topography | Elevation of sampling site |
| Slope | Topography | The rate of change of elevation for each digital elevation model (DEM) cell |
| aspect | Topography | The direction of the maximum rate of change in the z-value from each cell in a raster surface |
| CTI | Topography | Compound Topographic Index |
| DAH | Topography | Diurnal Anisotropic Heating |
| B1 | Vegetation | Sentinel-2 spectral band 1 (Wavelength: 443.9nm (S2A) / 442.3nm (S2B); Description: Aerosols) |
| B4 | Vegetation | Sentinel-2 spectral band 4 (Wavelength: 664.5nm (S2A) / 665nm (S2B); Description: Red) |
| B6 | Vegetation | Sentinel-2 spectral band 6 (Wavelength: 740.2nm (S2A) / 739.1nm (S2B); Description: Red Edge 2) |
| B9 | Vegetation | Sentinel-2 spectral band 9 (Wavelength: 945nm (S2A) / 943.2nm (S2B); Description: Water vapor) |
| B10 | Vegetation | Sentinel-2 spectral band 10 (Wavelength: 1373.5nm (S2A) / 1376.9nm (S2B); Description: Cirrus) |
| B11 | Vegetation | Sentinel-2 spectral band 11 (Wavelength: 1613.7nm (S2A) / 1610.4nm (S2B); Description: SWIR 1) |
| NDVI32 | Vegetation | Normalized Difference Vegetation Index in 32 days period |
| NBRT | Vegetation | Normalized Burn Ratio Thermal (NBRT) index in 32 days period |
| greenness | Vegetation | Annual Greenest Pixel in the year of 2017 |
| imprv | Habitat | Percent of the pixel covered by developed impervious surface |
| ptrcv | Habitat | Percent of the pixel that's covered by tree canopy |

### Compilation of environmental variables

We assembled environmental variables across six main categories: location, habitat, bioclimate, topography, human impact and vegetation (Supplemental Methods; Table S1; Text S1). All raster layers were aligned and projected to a unified 100 x 100 m grid from Google Earth Engine (Coordinate Reference System for this project: ESPG 4326, WGS84). Layers with a higher original resolution were down-sampled using a mean aggregation method. Layers with a lower or same original resolution were projected using bilinear method for continuous values and nearest neighbor method for categorical values. Layers were stacked and clipped to California’s extent, and used for point extraction. For coastal sites outside of the raster’s geographical coverage, values were extracted by the closest point available in 0.5 km radius. If the closest point fell outside of the radius, that site was assigned “NA” value. All computation and analyses were performed in R version 3.5.3 (R Core Team 2019). Raster operations were performed using R package *raster* (Hijmans 2017).

Considering that many environmental variables are correlated, we evaluated the Pearson’s correlation coefficient of the 56 numerical environmental variables and hierarchically clustered the variables according to the coefficients into variable groups using R functions *cor, hclust* and *cutree.* To reduce collinearity and improve interpretability in community modeling, we created a reduced set of numerical environmental variables that had an R^2^ < 0.8 (Table 1) for downstream analysis.

**Table 2.** Envfit result on PCoA ordination for each metabarcode. Here we present the three significant (P < 0.001) environmental variables with the highest correlation coefficient. The significance of the correlation was tested by 1999 permutations. For a complete result of all variables, please refer to Table S12. The direction of changes is included in Figure S20.

| Metabarcode | 1st variable | R<sup>2</sup> | 2nd variable | R<sup>2</sup> | 3rd variable | R<sup>2</sup> |
| --- | --- | --- | --- | --- | --- | --- |
| 16S | NDVI32 | 0.49 | greenness | 0.47 | B1 | 0.42 |
| 18S | NDVI32 | 0.51 | greenness | 0.49 | B1 | 0.43 |
| CO1 | orcdrc | 0.41 | ptrcv | 0.36 | NBRT | 0.33 |
| FITS | greenness | 0.52 | B1 | 0.5 | orcdrc | 0.46 |
| PITS | bio3 | 0.21 | sndppt | 0.2 | B11 | 0.13 |

### DNA extractions, amplification and sequencing

DNA extraction, amplification and sequencing followed Curd et al. 2019. Briefly, three 250 mg biological replicate soil samples from each site were fully homogenized and pooled per site. DNA was extracted using QIAGEN DNeasy PowerSoil Kit (Qiagen, Valencia, CA, USA) according to the manufacturer’s instructions. Negative controls were included in every batch of 12-18 extractions. DNA was amplified by polymerase chain reaction (PCR), using primers for five barcode regions: *16S* (515F and 806R; Caporaso et al. 2012), *18S* (Euk_1391f and EukBr; Amaral-Zettler et al. 2009), *CO1* (mlCOIintF and Fol-degen-rev; Yu et al. 2012, Leray et al. 2013), fungal *ITS1 (“FITS”;* ITS5 and 5.8S; White et al. 1990, Epp et al. 2012), and plant *ITS2 (“PITS”* ITS-S2F and ITS-S3R; Gu et al. 2013). For samples belonging to the coast transect, they were additionally amplified using *12S* barcode targeting fish (MiFish; Miya et al. 2015) for another study. Although the result is not described here, the raw sequence file contains those sequences. Primer sequences and thermocycling profiles can be found in Table S2.1-2. All PCR amplifications were performed in triplicate and with additional PCR negative controls. Triplicate positive amplifications confirmed by gel electrophoresis, were pooled by sample and barcode to equimolar levels (Supplemental Methods), indexed and sequenced on an Illumina MiSeq v6 platform for 2×300 bp reads (QB3-Berkeley FGL; University of California, Berkeley, CA, USA) with a target sequencing depth of 50,000 reads/sample/metabarcode. Five of the 278 sites were processed as biological replicates by different technicians to inspect taxonomic variation in independent DNA extraction and technical processing.

### Bioinformatics and data processing

We used default settings in the *Anacapa* Toolkit (Curd et al. 2019) for multi-locus sequence data processing and taxonomy assignment. In brief, quality control of raw sequences was performed using *Cutadapt* (Martin 2011) and *FastX-Toolkit* (Gordon et al. 2010), and inference of Amplicon Sequence Variants (ASVs) was made with *DADA2* (Callahan et al. 2016). Taxonomy assignment was made on each ASV using *Bowtie2* (Langmead and Salzberg 2012) and the Bayesian Lowest Common Ancestor algorithm *(BLCA;* Gao et al. 2017) on custom metabarcode-specific reference databases, created using *Creating Reference libraries Using eXisting tools (CRUX;* Curd et al. 2019). *Bowtie2* first aligns ASVs against the corresponding metabarcode-specific reference database and returns up to 100 alignments for each ASV. *BLCA* then determines the lowest common ancestor (LCA) from *Bowtie2* hits for each ASV and assigns a bootstrap confidence for each level of the taxonomic path. Taxonomy assignments with a bootstrap confidence cutoff score over 0.6 were kept for each ASV. ASVs with the exact same inferred LCA passing confidence filter were summed into one taxonomic entry as the species/phylotype/MOTU equivalent in this study and will be referred as “taxonomic entry” in the following text.

To informatically control for contamination, we further removed all singleton or doubleton ASVs, and ASVs that occurred more than or equal to three times in all blank samples from subsequent analyses (Table S3). Resulting ASVs from each primer were converted to *phyloseq* objects using the R package *ranacapa* (Kandlikar et al. 2018). For subsequent alpha and beta diversity analyses requiring rarefaction, we performed rarefaction in 10 replicates and took the mean using the *custom_rarefaction* function in the R package *ranacapa* (Table S2.3; Text S2). Reads with no assignment were not removed before rarefaction.

To evaluate how material aliquoted for DNA extraction and independent lab processing influenced taxon profiles, we estimated concordance in the decontaminated and further rarefied datasets between biological replicates (Text S3).

### Comparisons with Traditional Surveys

To compare the eDNA taxonomic results to traditional surveys, we compared eDNA results to the curated species inventory of the University of California Natural Reserve System (UCNRS), which records Chordata, Arthropoda, and Streptophyta (see Supplemental Methods). We counted how many taxon records were shared or unique to eDNA results or traditional records (UCNRS) at classification levels of order, family and genus combining all reserves and within each reserve.

We then developed a metric of traditional observation score (TOS) in eDNA taxonomic assignment. TOS uses all species observation and collection records in the Global Biodiversity Information Facility (GBIF) database from a broad region centered on California to score whether the taxon assignment of an eDNA ASV has been observed. A TOS > 0 suggests there is support for the assignment of an ASV based on its presence in the TOS region (see Supplemental Methods).

### Alpha Diversity

Alpha diversity was calculated using Observed and Shannon’s Diversity Index in R package *vegan* (Oksanen et al. 2019) to compare categorical variable groups (details in Supplemental Methods). The significance level was set at 0.05. The difference between each category was tested using the Kruskal-Wallis Test with Bonferroni correction for multiple testing. *Post hoc* analysis was performed using the Dunn Test (function *dunnTest* in R package *FSA;* Ogle et al. 2019).

We evaluated the relationships of alpha diversity measures and the reduced set of 33 continuous environmental variables (Table 1) as well using individual linear regressions (`alpha_diversity ~ variable`) of alpha diversity measures and environmental variables using function *lm* in R. To provide a more complete evaluation and account for the collinearity in the 33 variables, we also used partial least square (PLS) models with methods adapted from Lallias et al. (Lallias et al. 2015, George et al. 2019; see Supplemental Methods). PLS models are designed to handle regressions that have many, possibly correlated, explanatory variables and relatively few observations. We fit PLS models for alpha diversity in each metabarcode using function *plsr* in R package *pls* (Mevik et al. 2019) with three components included in the model (ncomp = 3), and the variable importance in projection measure was calculated using function *VIP.R* in R package *pls* extension (https://mevik.net/work/software/VIP.R).

### Beta Diversity

Unless otherwise specified, all functions mentioned in this section are from R package *vegan.* Community composition was visualized by plotting sample relative abundance of the top ten phyla for metabarcodes *16S, 18S,* and *CO1,* and top ten classes for *PITS* and *FITS*, partitioned by the major habitat. This was implemented by function *plot_taxa* in R package *microbiomeSeq* (Ssekagiri et al. 2017).

Composition was analyzed using unconstrained ordination. We calculated the binary Jaccard dissimilarity distance from the rarefied dataset for each metabarcode dataset and performed principal coordinate analysis (PCoA), permutational multivariate ANOVA (PERMANOVA) analysis by function *adonis* with 2999 permutations, and tests for the assumption of homogeneity of dispersion using the *betadisp* function (Supplemental Methods). *Permutest* function was used to test for the significance, with 2999 permutations. P values were adjusted using the Bonferroni method for both PERMANOVA and *betadisp* tests. To control for the effect of spatial correlation, partial constrained analysis of proximities (CAP) was done for the habitat and soil property variables while removing the effect of geographic coordinates. Partition of variance was evaluated using the *varpart* function. To control for the large variation in aquatic/coastal sites, PERMANOVA was repeated by excluding the samples from the coastal sites. We also partitioned the data by the four categories in the *majorhab* variable (aquatic; herbaceous; shrub and tree dominated habitats) and performed PCoA and PERMANOVA analyses within each major habitat. *Post hoc* explanation of the ordination axes was performed by fitting the reduced set of numerical variables (Table 1) onto the PCoA result using the *envfit* and *ordisurf* functions (Supplemental Methods).

### Zeta Diversity

To measure the fraction of unique categories of organisms held in common among nearby communities, we set cluster size to 4 nearby sites (Supplemental Methods) and calculated zeta four diversity (ζ_4_). The value of ζ_4_ was scaled for each geographic cluster as a fraction of its average taxonomic richness per sample (ζ_1_) using the function *Zeta.decline.ex* in the R package *ZETADIV* (Latombe et al. 2018). We tested the likelihood of two models of community assembly, using the Akaike Information Criterion (AIC) score within *ZETADIV,* through calculations of how zeta diversity decays with sampling order. Based on prior analyses (Hui et al. 2014) decays which follow a power-law of the form ζ_*N*_ = *ζ*_1_*N*^-*b*^, or an exponential of the form *ζ*_*N*_ = *ζ*_1_*e*^*b*·(*N*-1)^, were associated with a niche differentiation or stochastic process of community assembly, respectively. Scaled ζ_4_ diversity values were then plotted on a map of California using the R package *Leaflet* (Cheng et al. 2019).

Environmental factor groups were made by binning environmental variables according to their categories (Table 1). To determine the variation in ζ_4_ diversity attributed to either geographic distance or an environmental factor group, we used the function *Zeta.varpart* to analyze generalized linear models (GLM) generated using the function *Zeta.msgdm* within the R package *ZETADIV*. For each GLM we only used samples where both the geographic cluster ID and particular environmental factors were known.

### Gradient Forest Modeling

We used the gradient forest classification model in R package *gradientForest* (Ellis et al. 2012) to test which environmental variables best explained eDNA-detected taxon presence patterns across California using all 272 sites collected from three transects. Due to large variation in the coastal sites, we also performed additional analyses excluding all coastal sites using the same methods described below (details in Supplemental Methods). The gradient forest model was built with the reduced model containing 33 numerical environmental variables (Table 1; Figures S1,S2) that had a correlation coefficient < 0.8 with all other variables. We fit a classification-tree based gradient forest model using default settings to the biological matrix derived, but increased the number of trees to 2000 per family to increase the stability of the model (Breiman 2001). To assess model robustness and stability, we repeated the gradient forest model 20 times with the same settings and recorded the families with positive R^2^ values, the overall corrected R^2^, the importance of each explanatory variable. To assess model power and reliability, we used a permutation approach akin to Bay et al. 2018: we randomized the predictor matrix 100 times and ran the model (Supplemental Methods).

To visualize the community turnover gradient forest model over space, we used the input of all 33 environmental variables from 100 m x 100 m grids in the extent of California without extrapolation according to the gradient forest manual (Pitcher et al. 2011; Supplemental Methods). We used the top three principal components from the transformed environmental variables and visualized them by red, green and blue (RGB) bands (Ellis et al. 2012). Each environmental variable loading was visualized by function *biplot* in R. To differentiate model performance from the innate high-dimensional nature of the environmental variable matrix, we scaled the environmental variables and performed the same PCA and visualization procedure without using the model (“uninformed map”) and performed a mantel test and a monotonic regression between the biological matrix and either the *uninformed* map or *gradient forest informed* map.

### Network analysis

Results for each metabarcode were summarized by family, filtered on read depth and frequency and used in SPIEC-EASI ecological co-occurrence network analysis using R package *SpiecEasi* (Kurtz et al. 2015) for cross domain analysis that incorporates all five metabarcodes into one complex network (see Tipton et al. 2018). Topological parameters were determined in *Cytoscape* (v. 3.6.1; Shannon et al. 2003) using the *NetworkAnalyzer* tool. To observe the relationship between network degrees and the prediction R^2^ of each family from gradient forest, an ordinary least squares (OLS) linear regression model was made using the *lm* function in R and interactions were with R package *Interactions* (v. 1.1.1; Long 2019).

To evaluate the co-occurrence and gradient forest predictor patterns in a phylogenetic framework, the 915 families used in the gradient forest modeling were mapped onto the Open Tree of Life (tree.opentreeoflife.org) and a synthetic tree was generated using synthesis release v12.3. Datasets were mapped next to the phylogeny tips using the Interactive Tree of Life (https://itol.embl.de/).

## Results

### Metabarcoding result summary

The 278 selected samples that citizen scientists collected between March 2017 to July 2017 represented coast, shrub, and forest transects (Figure 1A; Table S1) and were sequenced to capture diversity across kingdoms. Each metabarcode recovers their target groups as expected (Table S2.1; Figure 1C), with *16S* amplifying Bacteria and Archaea, *18S* and *CO1* broadly amplifying eukaryotes including Animalia, Chromista, Fungi, Protozoa and some Plantae, *ITS1* targeting fungi (‘*FITS*’) selecting for Ascomycota and Basidiomycota, and the *ITS2* region targeting plants (‘*PITS*’) predominantly selecting for Chlorophyta and Streptophyta.

**Figure 1.**
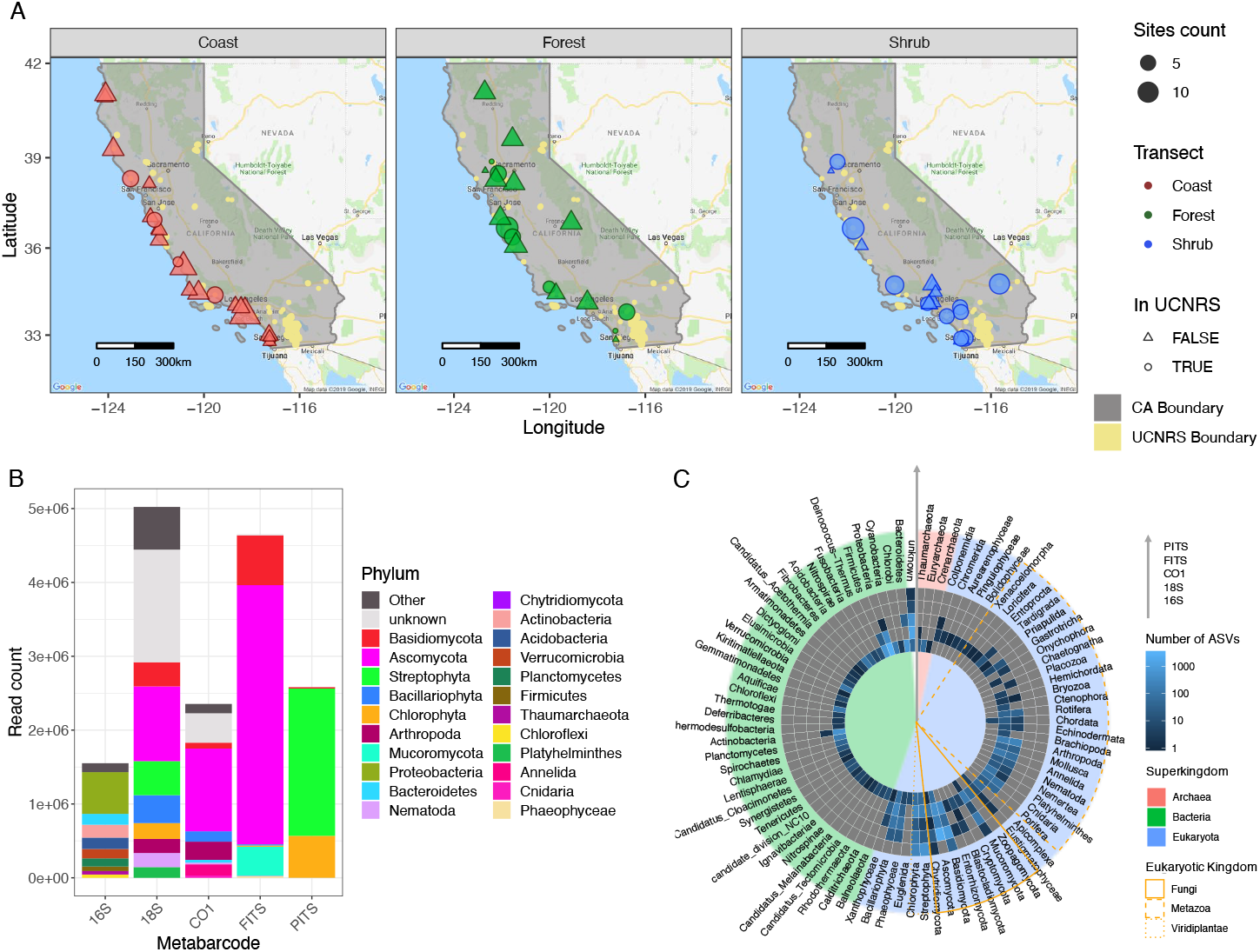
Map of 278 sites included in this study and illustration of taxonomic entries recovered with five metabarcodes. (A) Study area (gray shade) is defined within the State of California, United States. Sample sites are colored by three transect designations: coast (red), forest (blue) and shrub (green). Size of the points corresponds to the number of samples taken in the same area. Shape of the points (circle or triangle) represents areas within and outside of the University of California’s Natural Reserve System (UCNRS, yellow shade, area size not to scale for visibility). Circles are within UCNRS, triangles are not. (B) Read abundance is grouped by the phylum they belong to after taxonomy assignment and decontamination for five metabarcodes targeted Bacteria and Archaea *(16S),* Eukaryota *(18S),* Metazoa *(CO1),* Fungi *(FITS)* and Viridiplantae *(PITS).* Only the most abundant 10 phyla are plotted for each metabarcode. All other phyla are summarized in the “Others” category. (C) Heatmap shows each metabarcode’s taxonomic specificity. The results from each metabarcode *(16S, 18S, CO1, FITS, PITS)* are represented from inner to outer rings. Lighter color in one cell represents more taxonomic entries were recovered by that metabarcode for that phylum, gray color represents no entries. Phyla are indicated on the periphery. Background color of each pie wedge denotes the superkingdom (Red: Archaea, Blue: Eukaryota, Green: Bacteria, No background: Unknown) to which the phyla belonged at the time of taxonomy assignment (taxonomy file downloaded from NCBI at January 19, 2018). For eukaryotic phyla, Fungi (solid), Metazoa (dashed) and Viridiplantae (dotted) kingdoms are marked by different line types in an orange outline.

The 278 samples, five ‘biological replicate’ samples processed in duplicate by different technicians to assess the stability of results, and 21 negative controls as PCR blanks or extraction blanks, were sequenced, amounting to 75,830,796 reads for the five metabarcoding loci (Table S3). The average read depth per sample per locus was 54,554 reads. After several steps of quality control implemented in the *Anacapa Toolkit* (Curd et al. 2019), taxonomic assignment to NCBI-based reference databases, and subsequent sequence decontamination based on the negative controls, a total of 16,157,425 reads were assigned to 16,118 taxa from the five metabarcodes included in all transects. The median assigned read depth was 7,717 (Table S3; Figure S3) and mean taxa identified was 778 per sample.

A small test on variation in the replicate extraction and processing showed our methods were similar to what others have reported. An average of 97% taxonomic entries overlapped when negative results were counted and 44% entries overlapped when only presence was counted (Text S3; Table S4.2; Figure S4). Our eDNA metabarcoding results are sensitive to stochastic amplification even in triplicate PCR (Shirazi et al. *in prep*) and DNA extraction replicates may truly differ by small scale spatial heterogeneity (Lanzén et al. 2017). We interpreted this to expect zero-enriched data and less power to predict taxon presence with eDNA.

To prepare results for alpha and beta diversity analyses, sequence rarefaction was used to remove variation in sequencing depth. For the *16S* metabarcode, 210 samples were kept with 2,000 reads/sample; for *18S*, 221 samples were kept with 4,000 reads/sample; for *CO1,* 220 samples were kept with 1,000 reads/sample; for *FITS*, 225 samples were kept with 4,000 reads/sample; for *PITS,* 221 samples were kept with 1,000 reads/sample (Text S2; Table S2.3; Figures S5,S6). Despite fairly deep sequencing, stringent sample filtration was necessary to meet sufficiency metrics practiced by the metabarcoding community (Goldberg et al. 2016, Taberlet et al. 2018).

In summary, assignments spanned 81 phyla with most reads and taxonomic entries being assigned to Proteobacteria, Ascomycota and Basidiomycota (Figure 1B,C). The 16S metabarcode targeting prokaryotes generated most of the unique taxonomic entries (Table S3; Figure 1C).

### Comparison with traditional surveys: eDNA results partially overlap with traditional observations

Our comparison between metabarcoding results from within the UCNRS and a curated species list of UCNRS Streptophyta, Arthropoda and Chordata made by traditional surveys (Table S5) showed partial overlap across multiple classification levels. Forty-four Streptophyta families were only found in eDNA, 77 were only in traditional observations, 65 were recovered from both methods. We found 110 Arthropoda families were only recovered from eDNA, 139 were only in traditional observations, and 16 were recovered from both methods. No Chordata families were jointly recovered from both methods (Table S5.1). These results suggest many families are more conducive to being detected by either eDNA or observation, and the overlap is substantial in plants but not in arthropods. We acknowledge the seasonality of the UCNRS observations was not known, and could include many species that would not be expected to occur in the seasonal window eDNA was sampled (Bolger et al. 2000).

Our second comparison was to generate a Traditional Observation Score (TOS) for taxonomic lineages identified by eDNA using the GBIF records from Western North America and the Eastern Pacific. Only lineages resolved below the level of order were assigned a TOS, hence 974 lineages were omitted. Of the remaining lineages, only 5.6% of lineages had an adjusted TOS of 0, and 50% of lineages had an adjusted TOS of 1 (Table S6). Partial concordance was found in remaining lineages. No relationship was found between TOS and the frequency at which a taxon was found in eDNA samples (Pearson’s R^2^=0.004; P< .00001), suggesting the TOS is not heavily biased toward common or ubiquitous taxa. Because these two traditional observation tests showed high concordance at the family level, we selected family level classification for downstream gradient forest and network analyses.

### Alpha diversity varies at the local scale and across the terrestrial-marine interface

Three patterns emerged from alpha diversity Kruskal-Wallis tests of observed and Shannon Index alpha diversity of rarefied metabarcode datasets. First, we found spatial stratification for alpha diversity measures in the *loc* variable (name of places visited reported by volunteers) and *minorhab* (minor habitat) variable for all metabarcodes besides *CO1*, and stratification for the *clust* variable (neighboring cluster of sites within a radius of 0.5 km derived from GPS record) for *16S* and *FITS* (Figure 2A; Figure S7; Table S7), indicating bacterial and fungal alpha diversity is locally constrained in California. Second, transect was not significant for *18S* and *PITS* (Figure 2B; Figure S8) and major habitat *(majorhab)* was not significant in any tests (Figure 2C; Figure S9). Land use classification *(NLCD)* and *ecoregion,* yielded mixed results where Shannon and Observed tests conflicted in significance (Table S7). *Post-hoc* Dunn tests of *NLCD* in *16S* and *FITS* metabarcodes showed that samples belonging to open water, wetlands, and developed space were often part of the pairs that were different (Figure S10; Table S7.3). Third, we found substrate type (*SoS*) differed for all metabarcode tests except for *18S*, and *post hoc* tests showed sand samples were consistently lower in alpha diversity across metabarcodes (Figure 2D; Figure S11) compared to soil and sediment. Soil class (*taxousda*) tests showed no significance.

**Figure 2.**
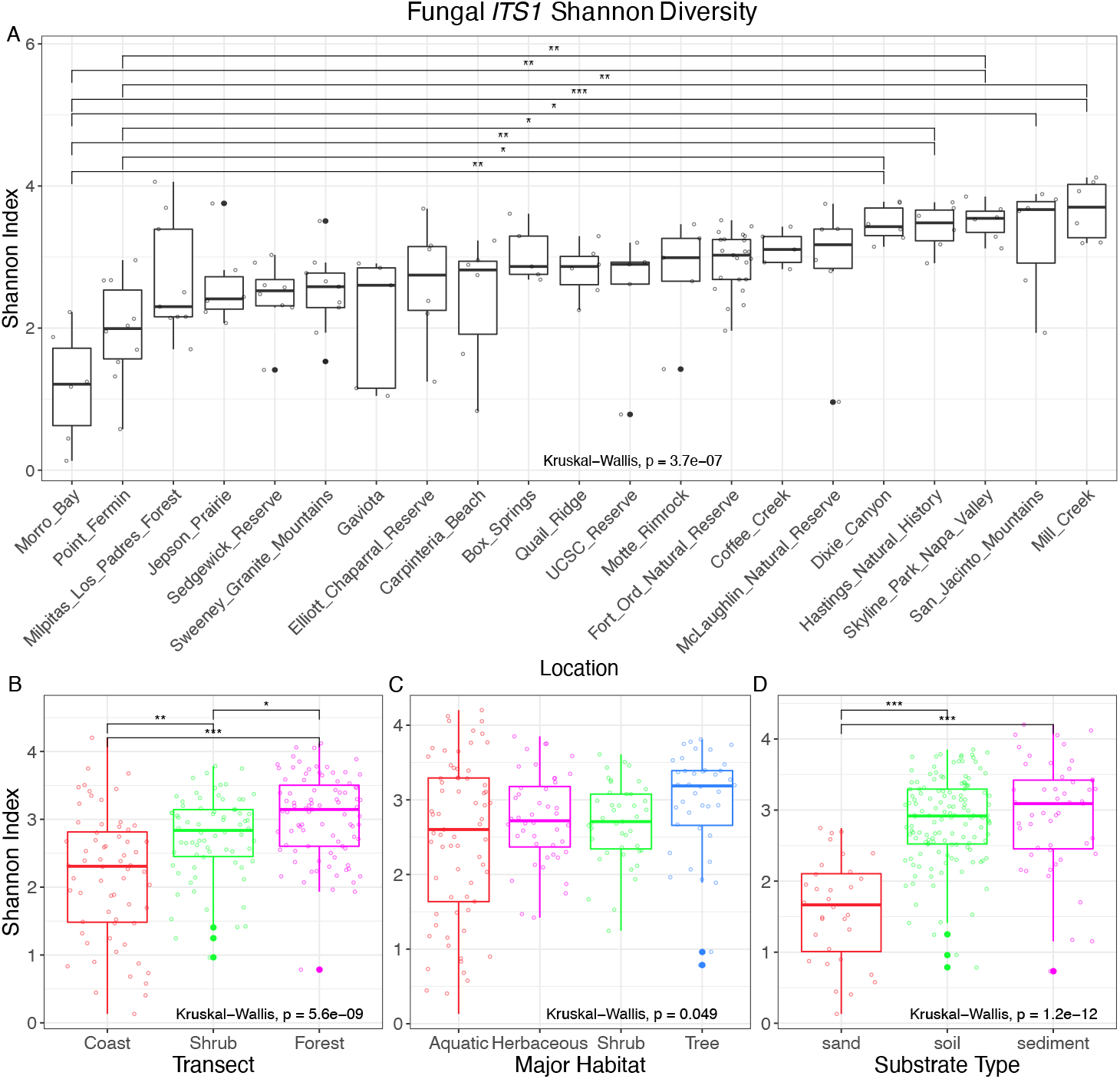
Boxplots of Shannon Index alpha diversity binned by different categories for the fungal ITS1 *‘FITS’* metabarcode. The Shannon diversity Index for rarified *FITS* metabarcode is grouped by (A) location, (B) transect designation, (C) major habitat designation and (D) substrate type designation. Horizontal notch stands for the medians. Lower and upper box ranges (hinge) represent the 25th and 75th percentiles. The whiskers extend to data points no more than 1.5 * IQR (inter-quartile range) from the hinges. All data points are plotted additionally as circles using *geom_jitter.* P-values from Kruskal-Wallis tests on mean alpha diversity metrics across categories are denoted at the bottom of each panel. Post-hoc pairwise Dunn Test significance is denoted as *P <0.05, ** P< 0.01, *** P< 0.001. Bonferroni Adjusted P values are in Table S7.

Individual linear regressions of alpha diversity statistics with continuous environmental observations showed fungal organisms *(FITS)* were best predicted compared to other metabarcodes, and five *FITS* models had an R^2^ > 0.2. Observed *FITS* alpha diversity with *greenness* had the highest correlation (R^2^ = 0.29; Table S8; Figure S12). In contrast, while a second analysis to evaluate the combined effect of the 33 environmental variables using PLS models also found the highest regression coefficients for *FITS* alpha diversity (R^2^ = 0.17 and 0.15), soil carbon content variable *orcdrc,* and *aspect* were the variables with the highest importance (Table S9).

### Beta diversity: Community composition varied by habitat characteristics

Bar plots showing relative abundance displayed similar patterns of variation across major habitats in *16S* and *FITS* results (Figure S13A,D). Bar plots of all other metabarcodes showed differences in composition and relative abundance profiles both within major habitats and between aquatic and terrestrial habitats. Differences were especially prominent in aquatic samples (Figure S13), where Ascomycota were largely absent and Chlorophyceae and Ulvophyceae were frequently dominant (Figure S13C,E). Interestingly, some *CO1* samples were dominated by Bacteroidetes, one of the most dominant bacterial phyla in the ocean (Figure S13C). We followed these observations of compositional diversity with several beta diversity analyses.

In community dissimilarity analysis, beta dispersion testing showed significant heterogeneity of multivariate dispersion (variance) within groups for all metabarcode and category combinations except *loc, majorhab, transect,* and *clust* for the *PITS* metabarcode (Table S10.1). PERMANOVA results showed significant differences among all different environmental variable categories (Table S10.2; Figures S14-S17). Beta diversity was significantly different across major habitat groups despite many overlapping sites in the ordination plots (Figure 3; Figures S14-S17). Groups exhibited differences both in within-group variation (dispersion) and in mean values of group centroids. In particular, samples from aquatic environments were more dispersed in the ordination (Figure 3A). To examine if this dispersion was driven by coastal samples that contain some community members restricted to marine systems, we removed coastal sites and re-examined community clusters (Figure 3C,D). Samples still exhibited clustering by major habitat in PERMANOVA and beta dispersion tests (Table S10.3-4; Figure 3C; Figure S18), suggesting observed aquatic beta diversity was also driven by freshwater sites.

**Figure 3.**
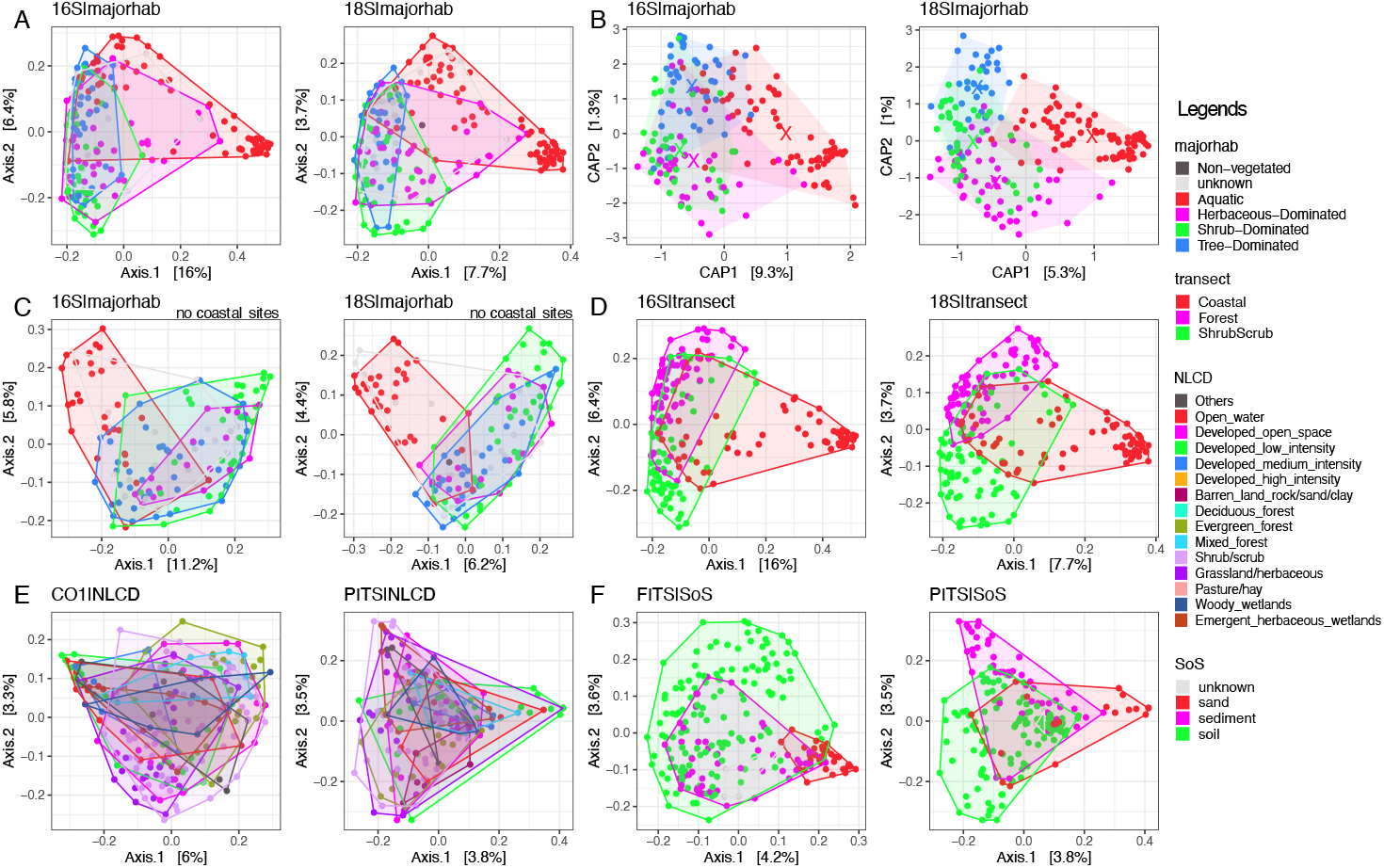
Beta diversity plots based on Jaccard dissimilarity. The first two principal coordinates are plotted with percentage of variance explained included in axis label. We show selected Principal Coordinate Analysis (PCoA) plots from (A) *16S* and *18S* for major habitat. For comparison, (B) shows a partial constrained analysis of proximities (CAP) of community composition by major habitat while excluding the location effect. The first two axes are plotted for each metabarcode. Each point stands for a sampling site. The “X” marks the position of each group centroid. Major habitats still substantially overlap in (B) compared to (A) for *16S.* Tree-dominated and herbaceous plant-dominated habitats show little overlap compared to (A) for *18S.* (C) Re-analysis of (A) but with coastal sites removed shows the aquatic and herbaceous-dominated space is greatly reduced. (D) Selected PCoA plots grouped by transect designations, showing samples belonging to coastal sites account for the greater amount of variation in PCoA Axis. 1. (E) Selected PCoA plots grouped by national land cover classification, showing developed space frequently overlap with other natural areas. Examples are for *CO1* and *PITS.* (F) Selected PCoA plots for *FITS* and *PITS* showing fungal beta diversity in sand is low compared to other substrates and compared to beta diversity for non-fungal taxa.

Minor habitat *(minorhab)* composition within each of the four major habitats was found to contribute strongly to dissimilarity (PERMANOVA, adjusted P < 0.01 in 2,999 permutations; Table S10.5-6). Jaccard dissimilarity PCoA revealed interesting features of beta diversity. In aquatic habitats, lacustrine and riverine samples overlapped and were largely separate from coastal minor habitats in all metabarcodes. The ‘marine intertidal zone III’ cluster separated from other marine and estuarine groups in all metabarcodes except for *16S*. In contrast, the *16S* community of ‘estuarine bay tidal flat’ was distinct from others when it was not for other metabarcodes (Figure 4). In herbaceous-dominated habitats, grassland samples were less dispersed in *16S* results than in other metabarcodes. In *PITS* results, minor habitat groups largely overlapped, but in all other metabarcodes, grasslands were distinct in composition. Scrub habitats were similar in *16S* and *CO1* but distinct in *FITS* communities. Mixed chaparral and coastal scrub overlapped in composition. In tree-dominated habitats, the coastal oak woodland, which was the most heavily sampled (Table S1.3), was dispersed (Figure 4). Other minor habitats formed smaller clusters, which were most separated from each other in the *16S* metabarcode results.

**Figure 4.**
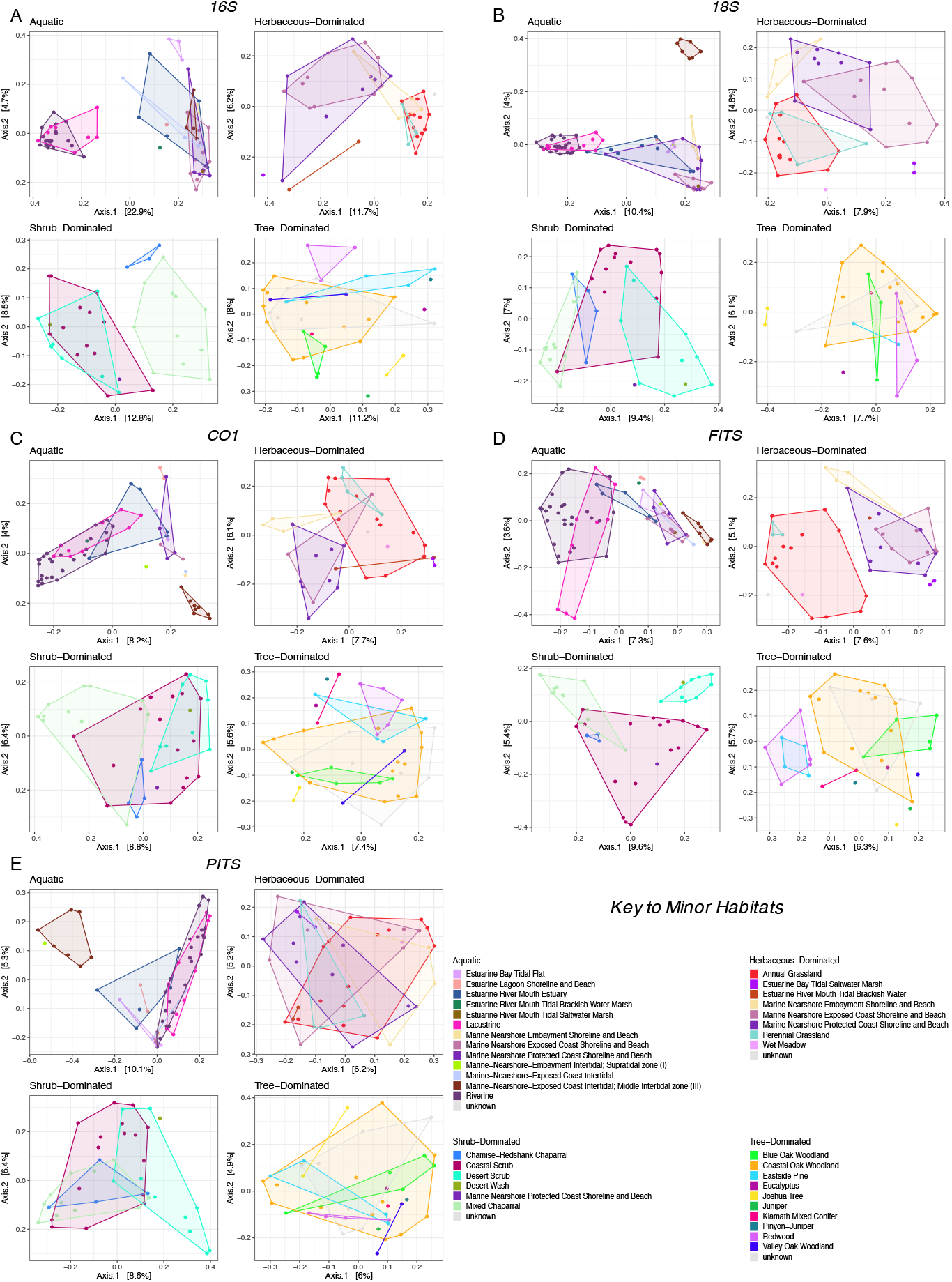
PCoA based on Jaccard dissimilarity with samples grouped by minor habitat and plotted within four major habitat categories. Results of PERMANOVA and beta dispersion show significant results (P < 0.001, number of permutations = 2999; Table S10.5-6) in minor habitat classification. Some minor habitat groups separate while others overlap, and patterns of compositional similarity (overlap) are different for different metabarcodes: (A) *16S*, (B) *18S*, (C) *CO1*, (D) *FITS,* (E) *PITS.*

CAP analysis removing the effect of spatial autocorrelation found that samples clustered tighter according to major habitat (Figure 3A,B; Figure S19). The unique contribution of major habitat (adjusted R^2^ ranging from 2.14% *(PITS)* to 9.65% (*16S*)) is larger than the contribution of local geographical categories (adjusted R^2^ < 2.00%, Table S11).

*Envfit* results showed that photosynthetic activities *(NDVI32* and *greenness)* were most highly correlated with *16S, 18S* and *FITS* (Table 2; Table S12; Figure S20). Soil organic carbon content *(orcdrc)* was most highly correlated with *CO1,* and Isothermality *(bio3)* was most highly correlated with *PITS.* Only 24 of 165 tests did not display significant relationships between variables and community composition. *Bio14* and *bio3* were the least frequently significant variables (Table S12).

### Zeta diversity species retention rate over space and composition predictability

Zeta diversity describes the degree of overlap in the number of unique categories of organisms held in common between *N* sites or communities (ζ_N_) (Figure S21A), which as *N* grows larger captures more variation due to turnover. This framework allows for an assessment in trends in regional scale turnover of relatively common organisms which are less biased towards the presence of rare, or spuriously detected taxa than the more established measures of α and β diversity (Hui et al. 2018). Environmental factor groups explained 1 to 32% of the observed variation in ζ diversity (Table 3). Vegetation variables were among the top predictors for *18S, CO1* and *FITS* datasets, with the highest variance explained at 32% for the *FITS* dataset. *16S* and *PITS* datasets had low ζ_4_ predictabilities. Variables related to small-scale location describe minimal variation (< 1%) in ζ_4_ diversity for communities (Table 3). To better understand spatial community stability, two models of zeta diversity decay were tested: the power law model and the exponential model. The power law model was found to be a better fit for communities described in all but the *PITS* metabarcode results, which followed the exponential decay model, suggesting lower spatial autocorrelation in plant and algal communities (Table S13; Figure S21).

**Table 3.** Variation in ζ4 diversity, for communities defined at the family level, attributed to geographic separation distance between samples versus variation in an environmental factor group for those same samples. Within each metabarcode, factor groups were ordered from lowest to highest contributions to variations in zeta diversity.

| Metabarcocode | FactorGroup | NumSamples | VarFactor | VarDistance | VarUnknown |
| --- | --- | --- | --- | --- | --- |
| 16S | Location | 184 | 0.00% | 0.29% | 99.70% |
| 16S | Topography | 184 | 0.94% | 0.17% | 98.90% |
| 16S | Habitat | 156 | 1.33% | 0.00% | 98.70% |
| 16S | Vegetation | 169 | 5.92% | 0.00% | 94.10% |
| 16S | BioClim | 184 | 7.17% | 0.00% | 92.20% |
| 16S | Soil Properties | 180 | 9.21% | 0.00% | 90.70% |
| 18S | Location | 184 | 0.14% | 0.00% | 99.90% |
| 18S | Habitat | 156 | 5.49% | 0.00% | 94.50% |
| 18S | Topography | 184 | 7.15% | 0.00% | 92.80% |
| 18S | BioClim | 184 | 7.30% | 0.00% | 92.70% |
| 18S | Soil Properties | 180 | 15.30% | 0.00% | 84.70% |
| 18S | Vegetation | 169 | 18.50% | 0.00% | 81.50% |
| CO1 | Location | 184 | 0.11% | 0.22% | 99.60% |
| CO1 | Habitat | 156 | 1.86% | 0.00% | 98.10% |
| CO1 | Topography | 184 | 3.30% | 0.46% | 96.20% |
| CO1 | BioClim | 184 | 12.00% | 0.00% | 88.00% |
| CO1 | Vegetation | 169 | 18.20% | 0.31% | 81.10% |
| CO1 | Soil Properties | 180 | 18.60% | 0.00% | 81.30% |
| FITS | Topography | 184 | 0.69% | 0.55% | 98.70% |
| FITS | Location | 184 | 0.93% | 0.38% | 98.20% |
| FITS | Habitat | 156 | 2.24% | 0.37% | 97.10% |
| FITS | BioClim | 184 | 18.50% | 0.00% | 80.40% |
| FITS | Soil Properties | 180 | 22.40% | 0.00% | 77.50% |
| FITS | Vegetation | 169 | 32.40% | 1.05% | 66.40% |
| PITS | Location | 184 | 0.03% | 0.00% | 100.00% |
| PITS | BioClim | 184 | 1.30% | 0.00% | 98.70% |
| PITS | Habitat | 156 | 2.16% | 0.00% | 97.80% |
| PITS | Topography | 184 | 2.98% | 0.03% | 96.90% |
| PITS | Soil Properties | 180 | 4.23% | 0.00% | 95.70% |
| PITS | Vegetation | 169 | 9.00% | 0.00% | 91.00% |

### Gradient Forest: Mapping biodiversity turnover in California

Our community mapping approach used a gradient forest model that inputs 272 sites x 915 families as a response variable matrix and 272 sites x 33 “reduced” set of environmental variables as a predictor matrix (Table S14). The gradient forest model explained 35% of variation in the biotic matrix, and all 915 families were effectively modeled (i.e. had an R^2^ > 0) and had high stability across 20 replicated runs (Average R^2^ = 0.349 ± 0.0004; Average families effectively modeled = 915 ± 0; Table S15). Using a permutation approach, we confirmed the mean overall R^2^ and number of families with positive R^2^ for true observations were higher than all the permuted runs (Figure S22). Many of the most responsive families were from marine aquatic sites, and some of these were low in observation frequency (Figure 5B; Figure S23). However, we did not observe a relationship between the observation frequency of a family and its gradient forest R^2^ (Figure S24), in contrast to what others have reported (Stephenson et al. 2018). This full gradient forest model showed elevation (*elev*), sand percentage (*sndppt*), photosynthetic activities based on vegetation greenness *(NDVI32)* and temperature of the wettest quarter *(bio8)* were the four most important predictors (Figure 5A) with a maximum of 0.018 in importance (Figure 5A).

**Figure 5.**
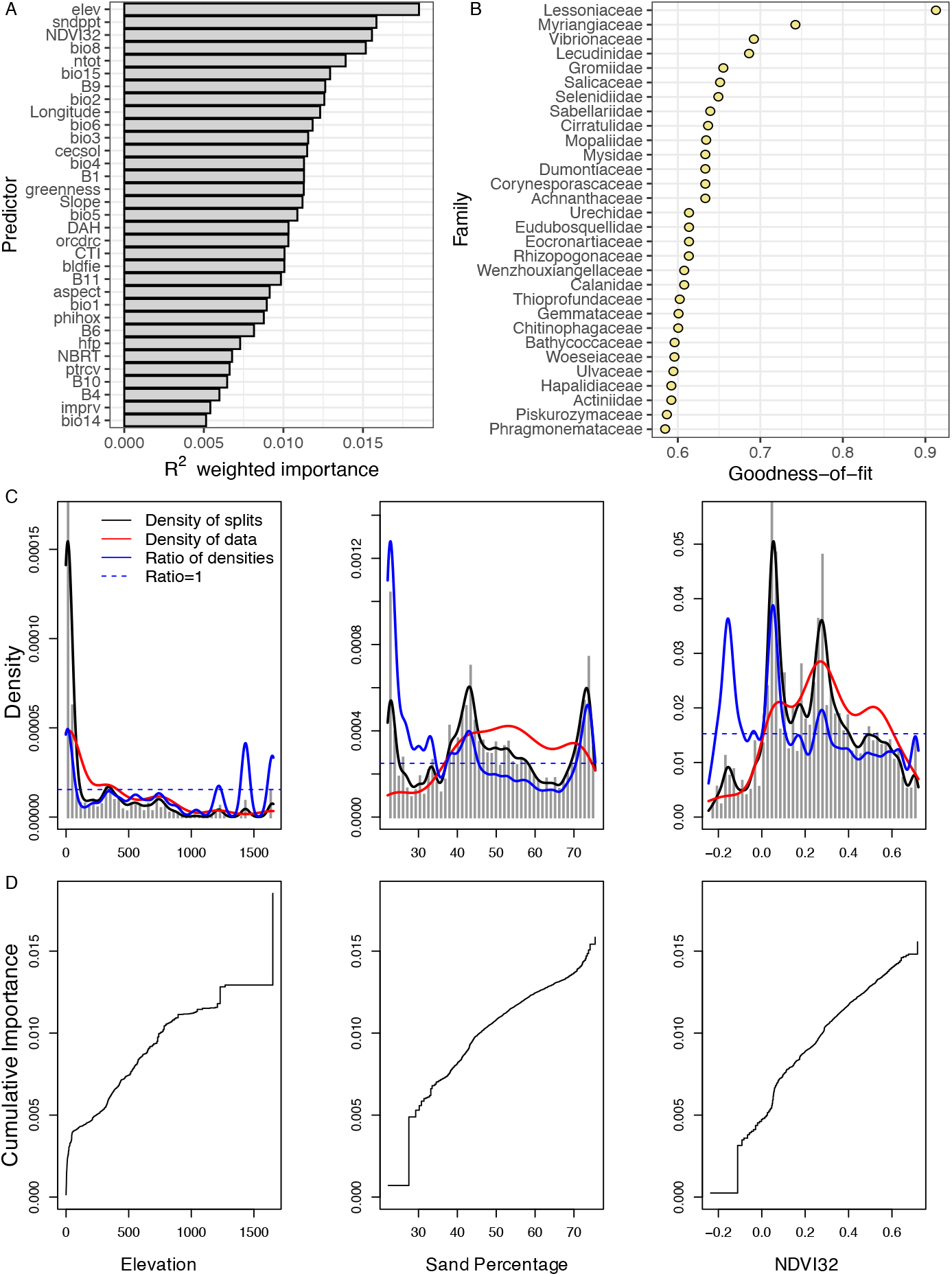
Gradient forest result for filtered CALeDNA dataset. (A) Ranked overall importance for 33 environmental predictors. (B) Ranked goodness-of-fit (1 – Relative Error rates) for the top 30 families (response variables). (C) and (D) show the community turnover along the three most important environmental gradients: elevation, sand percentage and photosynthetic activity proxy *(NDVI32).* (C) The gray histogram shows binned split importance at each gradient. Kernel density of splits (black lines), of observed predictor values (red lines) and of splits standardized by observation density (blue lines) are overlaid. The horizontal dashed line indicates where the ratio is 1. Each curve integrates to the importance of the predictor. (D) The line shows cumulative importance distributions of splits improvement scaled by R^2^ weighted importance and standardized by density of observations, averaged over all families.

Gradient forest provides information on the rate of community turnover along environmental gradients (Ellis et al. 2012). We plotted the relative density of splits and cumulative importance for environmental variables. Within the top three environmental variables, we found nonlinear community changes. For elevation, rapid community (high splits density) turnover occurred at 0 m and above 1,000 m (Figure 5C,D). For sand percentage, important splits were mainly distributed at 23%, 43% and 74% sand (local maxima with the highest density, Figure 5C,D), which have similarity to the soil texture triangle in the USDA system (Groenendyk et al. 2015). For photosynthetic activities *(NDVI32),* important splits were mainly distributed along −0.16, 0.05, and 0.28 (scale: −1 to 1; Figure 5C,D).

Our map of California biodiversity resembled EPA North America Level II and California Level III Ecoregion maps (U.S. Environmental Protection Agency 2010, 2012), which were created with different input data and methods (Figure 6C-E). In the gradient forest map (Figure 6A), the majority of central and southwestern CA community type (red) corresponded to Mediterranean California (Figure 6C. pale green, Level II 11.1.), characterized by medium photosynthetic activities *(NDVI32),* lower elevation (*elev*) and higher precipitation seasonality *(bio15).* The northwestern CA community type (purple) overlapped with Marine West Coast Forest (Figure 6C. blue, Level II 7.1.), characterized by a higher photosynthetic activity *(NDVI32)* compared to Mediterranean California. The northeastern CA type (green) corresponded with the Western Cordillera (Figure 6C. green, Level II 6.2.), with higher elevation (*elev*) and sand percentage (*sndppt*). The southeastern CA type (orange) was characterized by warm deserts (Figure 6C. yellow, Level II 10.2.), with lower elevation, almost no photosynthetic activity and higher temperature in the wettest month (*bio8*).

**Figure 6.**
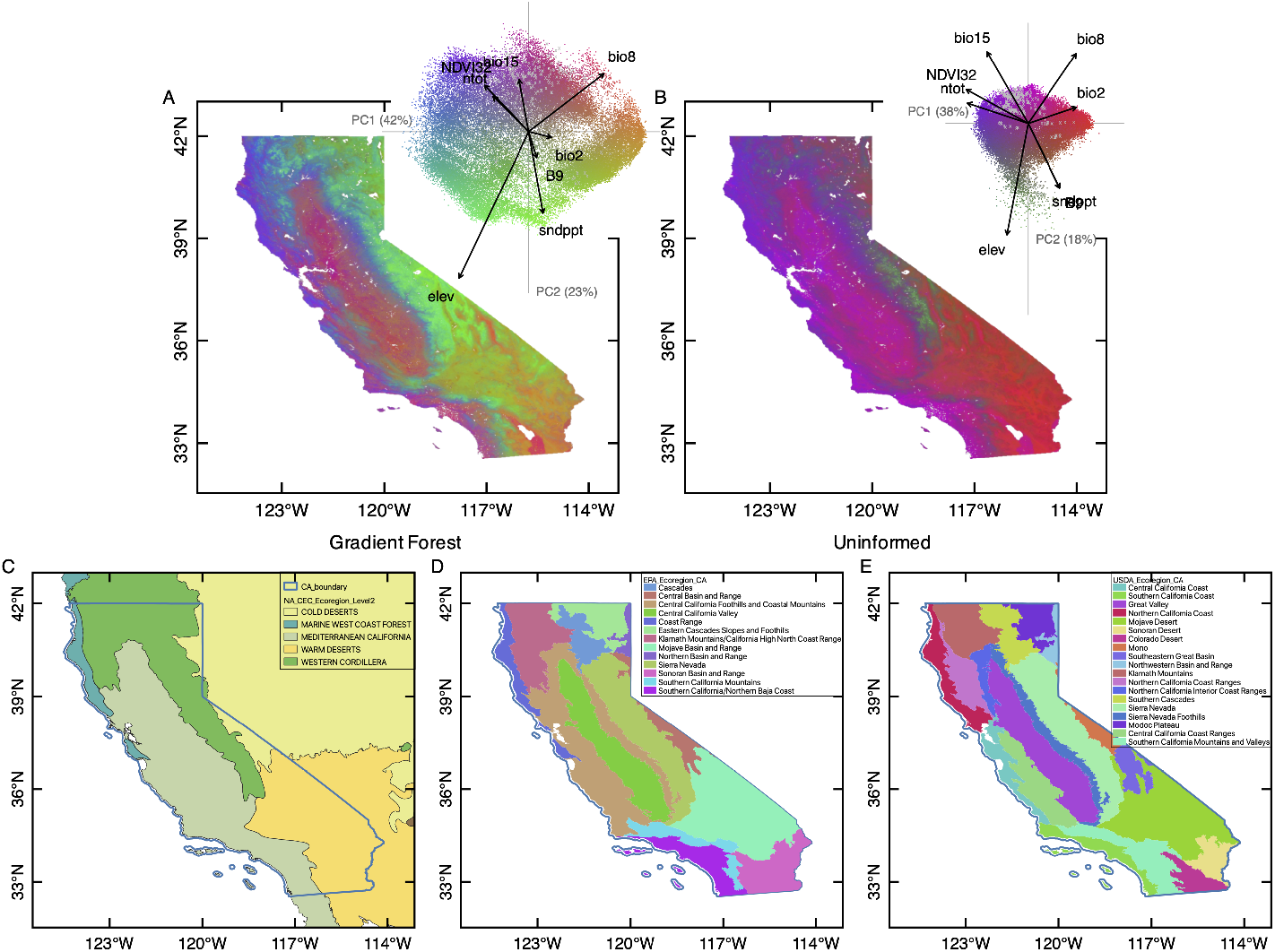
Map of transformed environmental variables following gradient forest predictions of biodiversity turnover from eDNA results (A) compared with uninformed, standardized environmental variables (B) and current major ecoregion maps (C-E) in California. The map shows the first three principal dimensions of biologically predicted (A) or uninformed (B) community compositions with an RGB color palette with 100 m resolution. The biplot of the first two PCs of the transformed environment space with (inset A) or without (inset B) biological information provides a color key for the compositional variation (n = 50 k). Similar colors approximate similar community in the transformed environmental space. The gray crosses denote the input eDNA sites (n = 272). Vectors denote the direction and magnitude of eight most important environmental correlates. (C-E) Selected major ecoregions maps are provided for comparisons with the gradient forest map (A). (C) EPA Level II Ecoregions of North America (U.S. Environmental Protection Agency 2010). (D) EPA Level III Ecoregions of California (U.S. Environmental Protection Agency 2012). (E) USDA Ecoregion Sections in California (USDA Forest Service 2007).

We checked the model prediction robustness by regenerating the California map without being informed by eDNA (Figure 6B), and these results were less visually similar compared with the eDNA-informed map (Figure 6A) and other California published maps such as the EPA North America Level II Ecoregion map (U.S. Environmental Protection Agency 2010, Omernik and Griffith 2014; Figure 6C). This purely physical approach of community turnover mapping showed adding eDNA improves gradient forest informed mapping limitedly by a 1.4% reduction in stress performance statistics; 5% increase in Mantel correlation R^2^ (Figure S25).

Because several of the most predicted families were marine, we generated a gradient forest model without coastal sites to examine which environmental variables shifted in their importance rank and to see how many families would still be modeled (meaning they transcend the coastal-fully terrestrial boundary). Without coastal sites, 802 families could still be effectively modeled and these had similar distribution of R^2^. In the model without coastal sites, we could still explain 30.4% of the variation in the biotic matrix. We observed the sharpest increases in *bio1* and human footprint (*hfp*) importance, and a sharpest decreases in *NDVI32* (where negative values correspond to water bodies; Weier and Herring 2000), *greenness*, and coastal aerosol *(B1)* importance (Figure S26) when coastal sites were removed. Other environmental variables were similar in importance and rank to the full gradient forest model.

### Ecological co-occurrence relates to gradient forest predictability

Co-occurrence patterns reflect biotic niche processes that maintain biodiversity patterns. We found a relationship between network degrees and family predictor R^2^ using an OLS linear model, which indicates environmental filtering (Horner-Devine et al. 2007). A family-level cooccurrence network produced 916 edges connecting 290 nodes (families) out of the total 304 families that met minimum frequency thresholds for analysis (Table S16; Figure 7A). The maximum edges per node was 48. The number of edges (degrees) per node showed no correlation with Neighborhood Connectivity (Pearson *r*=0.058) or Radiality (*r*=-0.113) but was highly correlated with Stress (*r*=0.900).

**Figure 7.**
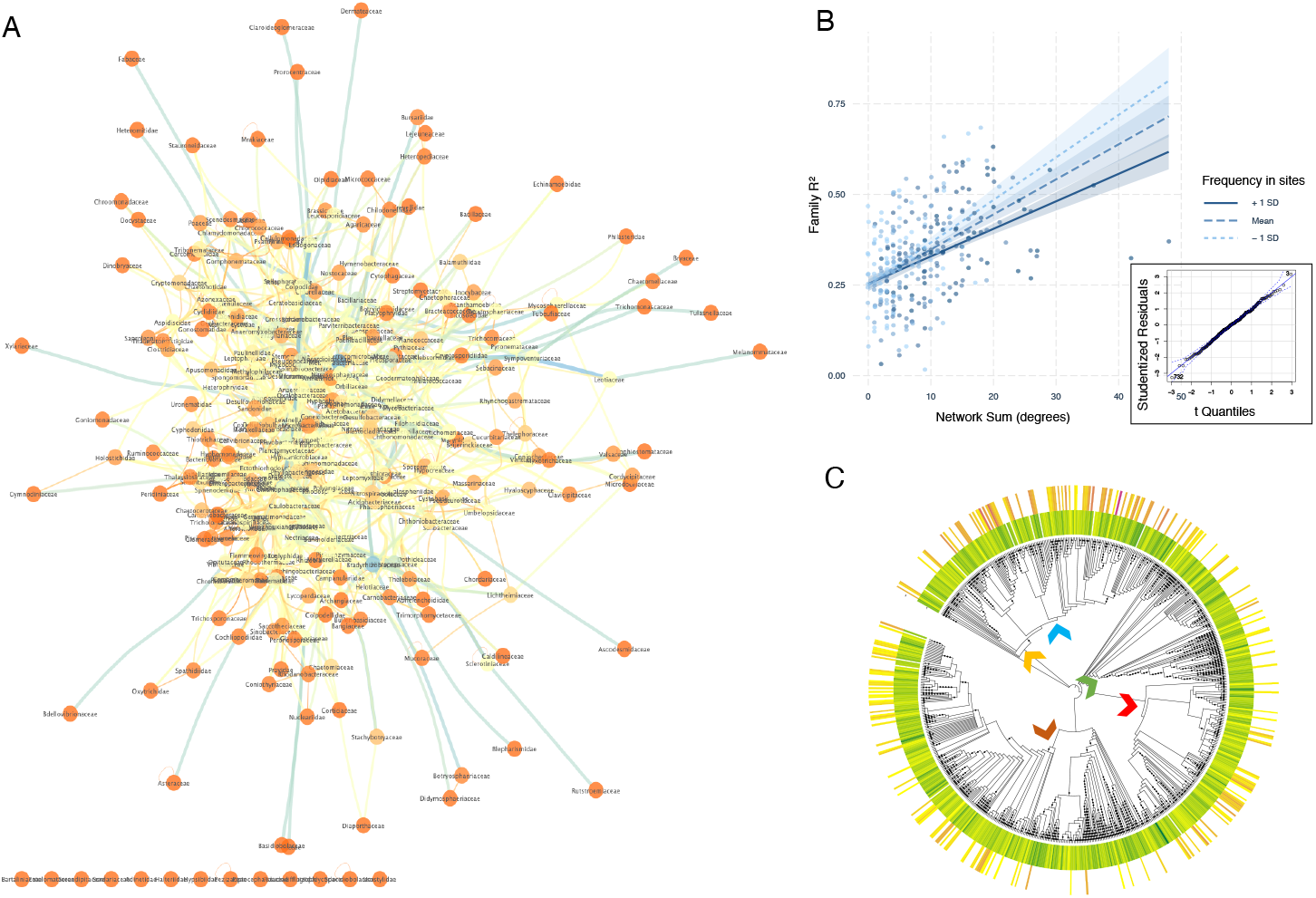
eDNA-based ecological co-occurrence network and relationship with gradient forest model goodness-of-fit R^2^. (A) 369 families (as nodes) are included in the network and 290 of those have at least 1 edge connecting them to another node. Colors are scaled by degrees for nodes and betweenness centrality for edges. Lower values have warmer colors. Node labels are families but are not intended to be legible in print. (B) OLS linear regression and quantile-quantile plot showing the interaction between network sum of degrees and frequency of taxa in sample sites with the dependent variable of gradient forest Family goodness-of-fit R^2^. There were 304 families included as joint observations in gradient forest and network results. The adjusted R^2^ = 0.22, network sum estimate = 0.01 (t-value=5.44; p=0.00), frequency in sites estimate = 0.00 (t-value = 0.18; p=0.86), and interaction between network sum and frequency in sites = 0.00 (t-value −2.38; p=0.02). (C) Phylogenetic tree made with the Open Tree of Life targeting input families as tips. Heatmap labels correspond to the range of gradient forest R^2^ (0.078-0.913) from yellow to dark green, and to the range of network degrees (0-48) from yellow to purple. Families too rare to be included in the network analysis (in fewer than 28 sites) are not colored in heatmaps. Arrows indicate the following clades: brown = fungi, mustard = Enterobacteriaceae, blue = Flavobacteriia, green = Streptophyta, red = SAR supergroup.

In the OLS linear model, interaction effects of site frequency were also considered. Model results showed a modest positive relationship (Adj R^2^= 0.22) between the number of edges and R^2^ for families, indicating the families determined by gradient forest to be under the most environmental filtering were also the families most integrated in ecological networks based on their numbers of degrees. However, the interaction between frequency in sites and network degrees was also significant (P<0.02; Figure 7B). In a phylogenetic analysis of these patterns, we observed that families with high network degrees and high gradient forest predictor values were widely distributed across clades and kingdoms, but heaviest in the clades containing the class Flavobacteriia and the SAR supergroup (Figure 7C), suggesting these communities might have lowest resilience under abiotic change and interacting partner drop out.

## Discussion

We have entered a new era of biomonitoring where species observations by the public outpace both field collections and on-the-ground observations made by scientists (Pearce-Higgins et al. 2018). With eDNA as a CCS tool (Biggs et al. 2015, Miralles et al. 2016, Sutter and Kinziger 2018), the public can be additionally armed with the capacity to inventory taxonomically broad local biodiversity from a small soil sample collection. Soils and sediments used in this study, collected by CCS volunteers, had an average of 778 taxonomic lineages identified in each sample. Several programs have demonstrated successes with eDNA as a CCS tool, and new innovations have made sampling easier for volunteers (e.g. Thomas et al. 2019) but thus far, most public efforts have targeted only a handful of species (e.g. algae and fish; JonahDNA.com). We demonstrate an approach using CCS collections obtained rapidly from a broad area within a seasonal snapshot. eDNA from these collections is co-analyzed with environmental data to explain and predict taxonomically broad biodiversity patterns, ultimately producing fine-scale beta diversity maps, and providing data that can advance ecological theory.

Our study mapped environmental data with beta diversity of varied taxonomic groups across California’s ecoregions at a higher resolution than currently available in statewide maps (Figure 6). Previous efforts have used climate and other abiotic data and models, and integrated them with traditional observational records such as herbarium specimens (Baldwin et al. 2017) to produce maps used to conserve threatened species (Jenkins et al. 2015), assess deforestation (Zarnetske et al. 2019) and evaluate species richness and endemism (Baldwin et al. 2017).

However, few mapping efforts coupled these data with remotely sensed variables such as from the Sentinel-2 instrument and local-scale observations of taxonomy biodiversity such as from eDNA. These additional measures enable community mapping at a finer grid size than 5 km (Jenkins et al. 2013, 2015, Pimm et al. 2014, Baldwin et al. 2017, Zarnetske et al. 2019). Fine-scale resolution has been shown to be better at aligning on-the-ground measures of biodiversity with spectral measures (e.g. Wang et al. 2018). As more relevant observations from space become available for analysis, community composition and turnover predictability should improve, which can help define areas that require distinct management (McKnight et al. 2007).

As a validation of our approach, our community turnover map aligned well at the regional scale with the EPA Ecoregion level III map (Figure 6D), although this conclusion is based only on a qualitative comparison. Nonetheless, there are fine-scale differences, such as the community designation near the San Francisco Bay (USDA Forest Service 2007, U.S. Environmental Protection Agency 2012). Our map is based on one season of eDNA observations from 2017 and thus has newer data than the observations that contribute to the EPA and USDA maps (Figure 6D,E). Such designation differences between maps suggest eDNA may detecting annual to decadal habitat conversion (Kadir et al. 2013). Future work should include comparisons of eDNA-enhanced maps from different seasons and years to assess community stability and the sensitivity of eDNA to temporal change.

California and other biodiversity hotspots are characterized by high floristic richness, local endemism, and high human impact (Hopper 1979, Myers et al. 2000). However, only recently with the advancement in DNA-based biomonitoring has species richness been systematically tested in microbial groups (Deiner et al. 2017). Our metabarcoding results from both macro- and microbial species confirm that richness exhibits local variation (Figure 2; Figure S7). Alpha diversity patterns observed in *18S*, *CO1* and *PITS* are consistent with past invertebrate, vertebrate and floristic community surveys that found richness does not vary according to mid-scale environmental partitions within a biodiversity hotspot (Pryke and Samways 2009, Demarais et al. 2017; Table S7; Figure S9). We assessed which metabarcode’s alpha diversity patterns could be modeled with environmental variables and found fungi (*FITS*) were predicted by indices of photosynthetic activity *greenness* and *NDVI32* (individual linear models LM; Table S8; Figure S12), soil carbon and aspect (partial least square models PLS; Table S9). These LM results suggest plant-soil reinforcement such as rhizosphere interactions may drive fungal richness (Timling et al. 2014, Tedersoo et al. 2014, Prober et al. 2015, Yang et al. 2017, Erlandson et al. 2018). We note indices of photosynthetic activity have not been included as part of most microbiome studies (Karimi et al. 2018, Bahram et al. 2018, George et al. 2019) so their importance is still being discovered. For the subset of studies we found that had included NDVI as a predictor, it was found to be important in modulating soil fungal and herbivore nematodes communities (Timling et al. 2014, Yang et al. 2017, van den Hoogen et al. 2019). We interpret the PLS results to indicate that soil carbon distribution in California is largely connected to fungal activity such as from saprobes (Erlandson et al. 2018), globally dominant fungal decomposers that assort into ecological guilds (Vĕtrovský et al. 2019).

Latitude, temperature, precipitation and pH were not significant in alpha diversity analysis (Tables S8,S9) although other studies have highlighted these factors as major correlates of alpha diversity (Fierer and Jackson 2006, Tedersoo et al. 2014, George et al. 2019, Crowther et al. 2019), explained by Rapoport’s Rule (Stevens 1989). We have considered that this discrepancy may be partially due to our omission of environmental extremes; for example, all samples had near neutral pH values (pH: Min = 5.05; Max = 8.50; Median = 6.40; Table S1.4).

However, such gradients have only been shown to affect certain taxa, and therefore the breadth of phylogenetic biodiversity encapsulated in each metabarcode may impair the detection of taxon-specific alpha diversity relationships. For example, while legumes (Garcillán et al. 2003) exhibit a latitudinal gradient with richness, invertebrates do not (Kerr 1999) and both of these groups are represented in the *18S* results.

Our beta diversity analyses show that most environmental categories can significantly partition samples according to taxonomic composition (Table S10; Figures 3 and 4; Figures S14-S17), suggesting that surface communities are largely filtered by ecology rather than neutral processes (Bahram et al. 2018). These patterns remained significant after exclusion of the more dispersed coastal sites and location effects (Table S10.3-4; Table S11; Figures S18,S19). However, we found substantial overlap in community composition ordinations, as has been shown in the global Earth Microbiome Project (Thompson et al. 2017) and regional soil biodiversity ordination plots (George et al. 2019; Table S10; Figure 3). In our ordinations, groups separated from each other when fine-scale categories are used, such as minor habitat within partitioned major habitat, suggesting a large amount of community partitioning is harbored within major habitats categories (Figure 4). We found prokaryotic diversity was particularly diagnostic of minor habitats in ordinations (Figure 4A). California is known for shifting ecotones that make management units unstable (e.g. Hennessy et al. 2018). We propose community classifications that include prokaryotes could be developed from analysis of eDNAbased composition and habitat features and these could serve as quantitative metrics to detect sites transitioning from one microhabitat type to another (Warton et al. 2015).

Examination of community compositions in site clusters using zeta diversity found evidence of high spatial decay in plants but not in other groups. Zeta (ζ) diversity is particularly useful for diagnosing local compositional variation. Compared with beta diversity that utilizes pair-wise dissimilarity, ζ4 diversity partitions regional diversity (gamma diversity) in three or more assemblages (Hui and McGeoch 2014; Figure S21A). Zeta diversity decay exhibits a power law relationship with sampling order, whereas the *PITS* dataset of Chlorophyta and Streptophyta is exponential (Table S13; Figure S21F). This observation suggests that plant-algal guilds are stochastic across space and may be especially sensitive to microclimates and stratified local nutrient availability as supported by previous studies (Sebastià 2007, Eisenlohr et al. 2013). We suggest this pattern be more extensively examined in future soil eDNA inventories of the kingdom *Plantae* to confirm if stochasticity is truly reflective of spatial arrangement or influenced by the chance of detection with DNA-based methods. The latter could be biologically dependent (e.g. presence of saprobes to process tissue and DNA), physically dependent (e.g. cell shedding), or chemically dependent (e.g. suppression of amplifiable DNA by phytochemicals). As an emerging field, we strongly encourage more attention on optimizing alignment between plant eDNA ζ diversity and plant distribution on the landscape.

Environmental variables (Tables 2 and 3) can have power to predict general biotic patterns and can illuminate possible drivers of community turnover (Figure S20) because they can readily be compared across studies (Omernik and Griffith 2014). For example, photosynthetic activities (NDVI32/greenness) had the highest correlation with the observed beta diversity structure in bacteria (*16S*), eukaryotes *(18S)* and fungi *(FITS)* in the *envfit* analyses (Table 2; Figure S20). Previous studies have supported a similar correlation of plant productivity and microbial beta diversity in global drylands (Delgado-Baquerizo et al. 2016) and in Tibetan Plateau (Yang et al. 2017). Isothermality (*bio3*) has strong positive associations with *PITS* turnover, suggesting inland arid California regions with low isothermality display nestedness in the biodiversity encompassed by these markers, as has been shown with plants in Australia (Gibson et al. 2012) and in South American seasonally dry forests (Silva and Souza 2018). Organic carbon *(orcdrc)* was strongly associated with *CO1* community turnover, which mirrors associations reported in soil macrofaunal communities, particularly nematodes (Jackson et al. 2019). Overall, zeta diversity largely support the *envfit* results, although *zeta diversity* had poorer explanatory power for *16S patterns,* which can be attributed to its greater sensitivity to common groups (Table 3; Simons et al. 2019) such as the nearly ubiquitous taxa in Proteobacteria.

Because so many variables have a relationship with observed beta diversity, as well as being correlated with each other (Figure S1), joint statistical analysis of multiple environmental variables is attractive to model biodiversity. However, examination of single variables will remain important as new regions are added to CALeDNA surveys. For example, it is unclear whether any of these patterns hold in agricultural regions of California such as the Central Valley, where we have few CCS samples. Agriculture has been proposed to ‘reset’ many of these abiotic constraints (Li et al. 2020). Single variables will also remain important to compare studies, given there is no standard set of variables. The use of Essential Biodiversity Variables could better enable comparative study of joint multivariable predictions of beta diversity (Jetz et al. 2019).

We demonstrate that using a suite of remotely sensed or modeled variables with regional or global coverage, family-level taxa presence could be strongly predicted and used to produce community turnover maps (Figures 5 and 6). Both the full gradient forest model and the model without coastal sites show that there are no predictors that are substantially stronger than others (Figure 5A; Figure S26A). Our full gradient forest model is in agreement with many proposed patterns of how the environment filters communities. Elevation (*elev*), sand percentage *(sndppt),* photosynthetic activities (*NDVI32*) and mean temperature in the wettest quarter (*bio8*) were the among the most important predictors (Figure 5A) and all of these variables had been proposed to be prominent drivers in community structures worldwide. Similarly, microbial turnover patterns had been linked to elevational gradients (Collins et al. 2018, Peters et al. 2019) and to particle size fractions (Sessitsch et al. 2001, Ehrlich et al. 2015). Our analysis complements such findings of variation in the rate of compositional turnover along environmental gradients, which have been the focus of many traditional ecology case studies. For example, fast community turnover was proposed at 0 m elevation (Figure 5C), recapitulating the observation that communities are highly structured and patchy in these coastal areas, which was also suggested by the *envfit* results to *B1* (coastal aerosol).

Finally, we suggest eDNA ecological network analyses should be leveraged so that the biotic interaction dependence can be contrasted with dependence or sensitivity to the abiotic environment. Our work shows a positive relationship between the number of degrees a family has and its propensity for environmental filtering based on gradient forest predictability, and shows there is phylogenetic conservation of this positive relationship (Figure 7). These results suggest that climate change and other disturbance can lead to community network dropout and cause a cascade of community change that may not be anticipated. Other studies focused on a single kingdom have come to similar conclusions, such as in microbial variation in an altitudinal gradient in the Atacama Desert, Chile (Mandakovic et al. 2018). However, using eDNA for these tests remains only possible at a coarse level of classification because reference sequence databases are incomplete and metabarcodes are not completely diagnostic. Ongoing efforts to sequence species and build a global taxonomic biodiversity database in the next decade (e.g. the Earth BioGenome Project, Lewin et al. 2018; the Centre for Biodiversity Genomics, Hobern 2020) are positioned to ameliorate this challenge in the coming years.

Several problems persist for eDNA-enabled environmental biology and increasing the confidence and resolution of eDNA-based detection is crucial for effective biomonitoring. First, a detection bias in favor of small body size is evident in eDNA studies (Figure 1; Tables S5,S6). Including other loci in metabarcoding (e.g. Andersen et al. 2012) or employing DNA capture approaches to target larger organisms (Seeber et al. 2019) may improve detection of large-bodied species. Novel tools such as camera traps or satellite telemetry (e.g. NASA CubeSats) to increase detectability for large organisms could be used to complement eDNA results, as could more participation by CCS to add iNaturalist data and to scrutinize eDNA results (as helped produce the TOS score; Table S6). Second, different DNA extractions from the same sample exhibit taxonomic heterogeneity (Text S3; Table S4). We are devoting efforts to examining stability and stochasticity of taxonomic profiles under varied sample processing (Castro et al. *in prep*) and DNA library preparation steps (Shirazi et al. *in prep)* in response to calls for attention to these potential biases (Prosser 2010, Goldberg et al. 2016), and were careful in this study to use approaches that are considered a standard of the field for reducing these biases.

We have shown that eDNA metabarcoding combined with remote sensing information can advance biodiversity assessment capacity. Space, flight, tower and drone-based remote sensing information is becoming increasingly available and accessible (Pettorelli et al. 2014). These data now can provide more direct, spatially continuous measures of plant functional diversity and ecosystem functioning at regional (Schneider et al. 2017, Durán et al. 2019) to global scales (Schimel et al. 2019, Bae et al. 2019). Using remotely sensed variables will not only elucidate classical community-environment interactions, but can also provide an opportunity to predict biodiversity turnover across large landscapes even when biological samples from all locations are difficult to acquire (Bush et al. 2017).

In conclusion, we demonstrate the emerging potential of coupling CCS observations and eDNA data from samples that CCS volunteers collect with remote sensing and ecological modeling to assess community-environment interactions and ultimately map community turnover. We provide one of the most comprehensive surveys of terrestrial biodiversity across three domains of life over a large, environmentally diverse state. We show the predictive and explanatory power of environmental variables on alpha, beta, and zeta diversity across highly diverse regions and at local geographic scales. The beta diversity map for California, illustrating a continuous surface of community turnover, is validated by its similarity to the standard US Ecoregion maps. Computationally intensive and artificial intelligence driven models are producing maps for mitigating the challenges of global change (Stephenson et al. 2018, Harfouche et al. 2019). Our approach contributes to the development of strategies to model living systems by integrating molecular-assisted observations with traditional and remotely sensed biodiversity measures. By encouraging more eDNA sequencing and CCS across broad regions in narrow temporal windows, and by systematically using Essential Biodiversity Variables in analyses, a new arsenal can be applied to tracking and predicting ecological change. Such data will allow the dissection of the contributions of biotic and abiotic factors to system resilience. Multidisciplinary initiatives that include specialists and CCS together can energize a necessary constituency for conservation.

## Data availability

Scripts and data associated with the analyses are archived in Zenodo archive (link pending) and github repository (https://github.com/meixilin/caledna_transect). The raw sequencing data will be deposited in the NCBI Sequence Reads Archive (accession pending).

## Author Contributions

RSM, RKW, BAS, EEC designed the study. ML, RSM, RH, ALS, MO, EEC and ZG selected analyses. RSM and EEC coordinated public sampling. ML, EF, TAS, and RSM made DNA libraries. ML, MPM, FS, AGV, DRR, RSM and EEC curated environmental metadata. ML, FS, and RH generated statewide data layers. ML led biodiversity and gradient forest analyses, and ML, ALS, and RSM generated plots. EJM generated the synthetic phylogeny. All authors performed analyses and interpretation. ML, RSM, and RKW wrote the manuscript with input from all authors.

## Acknowledgement

Funding for the CALeDNA sample processing, infrastructure, and personnel was provided by the University of California Research Initiatives (UCRI) Catalyst grant CA-16-376437 and Howard Hughes Medical Institute (HHMI) Professors Grant GT10483. Additional funding for personnel and computational infrastructure was provided by the National Science Foundation (NSF) 1759756. The research carried out at the Jet Propulsion Laboratory, California Institute of Technology, was under a contract with the National Aeronautics and Space Administration (80NM0018D0004). Government sponsorship is acknowledged. Graduate student support was additionally provided by the National Council for Scientific and Technological Development of Brazil [Grant No. 209261/2014-5] and the University of California, Los Angeles Department of Ecology and Evolutionary Biology (EEB) Summer Research Fellowship. We thank the UC Natural Reserves System managers, other natural areas managers, and the hundreds of volunteers for collections. We thank A. Mahinan, N. Stavros, W.-Y Kwan, for assisting with spatial environmental data and to A. DeVries and L. Bulbenko for preparing extractions. We thank C.M. Mueller and Z. Kurtz for help optimizing analyses.

