## Appendices for "A Biodiversity Composition Map of California Derived from Environmental DNA Metabarcoding and Earth Observation"

### Table of Contents

*Appendices* ... *1*
*Supplemental Tables* ... *4*
*Supplemental Table Captions* ... *5*
**Table S1** ... 5
**Table S2** ... 5
**Table S3** ... 5
**Table S4** ... 5
**Table S5** ... 6
**Table S6** ... 6
**Table S7** ... 6
**Table S8** ... 6
**Table S9** ... 6
**Table S10** ... 6
**Table S11** ... 7
**Table S12** ... 7
**Table S13** ... 7
**Table S14** ... 7
**Table S15** ... 7
**Table S16** ... 7
*Supplemental Methods* ... *8*
**Sampling design** ... 8
**Compilation of environmental variables** ... 9
**DNA extractions, amplification and sequencing** ... 11
**Bioinformatics and data processing** ... 12
**Comparisons with traditional surveys** ... 13
**Alpha diversity** ... 14
**Beta diversity** ... 16
**Zeta diversity** ... 17
**Gradient forest modeling** ... 18
**Network analysis** ... 21
*Supplemental Text* ... *23*
**Text S1: Habitat designation method** ... 23
Methods ... 23
**Text S2: Evaluation of rarefaction depth** ... 24
Methods ... 24
Results ... 24
**Text S3: Concordance in biological replicates** ... 26
Methods ... 26
Results ... 26
*Supplemental Figures* ... *28*
**Figure S1. Metadata correlation** ... 28
**Figure S2. Environmental variable distribution** ... 29
**Figure S3. Read depth summary** ... 31
**Figure S4. Replicate PCoA** ... 32
**Figure S5. Rarefaction summary** ... 33

**Supplemental Tables**

Are supplied as a zip archive named "Supplemental_Tables_CALeDNA_EA.zip" that contains the following files.

- S1_List_and_explanation_environmental_variables.xlsx
- S2_Metabarcoding_primer_info.xlsx
- S3_decontaminated_ASVs_five_metabarcodes.xlsx
- S4_concordance_in_replicates.xlsx
- S5_UCNRS_site_comparison.xlsx
- S6_TOS_GBIF.xlsx
- S7_kruskal_alpha_div.xlsx
- S8_individual_lm_alpha_div.xlsx
- S9_pls_alpha_div.xlsx
- S10_beta_dispersion_permanova_beta_div.xlsx
- S11_capscale_varpart_beta_div.xlsx
- S12_envfit_beta_div.xlsx
- S13_model_stats_decay_pattern_zeta_div.xlsx
- S14_gradient_forest_input.xlsx
- S15_gradient_forest_output.xlsx
- S16_network_analysis.xlsx

**<u>Supplemental Table Captions</u>**

***Table S1***

Table S1.1 Metadata table collected for the 278 sites.

Table S1.2 Full list of metadata variables and explanation.

Table S1.3 Data distribution for categorical variables.

Table S1.4 Data distribution for numerical variables.

Table S1.5 Data distribution for the raster variables collected in the range of California.

***Table S2***

Table S2.1 Metabarcodes used in this study.

Table S2.2 PCR reaction setup and cycling info.

Table S2.3 Rarefaction depth chosen for each metabarcode.

***Table S3***

Table S3.1 Decontaminated ASV table for the *16S* metabarcode.

Table S3.2 Decontaminated ASV table for the *18S* metabarcode.

Table S3.3 Decontaminated ASV table for the *CO1* metabarcode.

Table S3.4 Decontaminated ASV table for the *FITS* metabarcode.

Table S3.5 Decontaminated ASV table for the *PITS* metabarcode.

***Table S4***

Table S4.1 Concordance between biological replicates in the rarefied dataset.

Table S4.2 Concordance between biological replicates in the unrarefied dataset with minimum total reads cutoff of three.

Table S4.3 Concordance between biological replicates in the unrarefied dataset with minimum total reads cutoff of a hundred.

***Table S5***

Table S5.1 Summary of comparisons between eDNA results and the curated species inventory of the University of California Natural Reserve System (UCNRS).

Table S5.2 eDNA and UCNRS records comparisons within each reserve.

***Table S6***

Table S6 Traditional observation score of all taxonomic entries identified with eDNA.

***Table S7***

Table S7.1 Summary statistics for alpha diversity.

Table S7.2 Alpha diversity Kruskal-Wallis testing result summary.

Table S7.3 Post-hoc testing results following significant Kruskal-Wallis tests.

***Table S8***

Table S8 Result table of individual linear regression models on alpha diversity measures and environmental variables.

***Table S9***

Table S9 Result table of partial least square models on alpha diversity measures and environmental variables.

***Table S10***

Table S10.1 Result table of beta dispersion testing in beta diversity.

Table S10.2 Result table of PERMANOVA testing in beta diversity.

Table S10.3 Result table of beta dispersion testing in beta diversity excluding coastal samples.

Table S10.4 Result table of PERMANOVA testing in beta diversity excluding coastal samples.

Table S10.5 Result table of beta dispersion testing in beta diversity within major habitats.

Table S10.6 Result table of PERMANOVA testing in beta diversity within major habitats.

***Table S11***

Table S11.1 Result table of the constrained ordination test in beta diversity analyses.

Table S11.2 Result table of the variance partitioning test in beta diversity analyses.

***Table S12***

Table S12 Summary of envfit result in PCoA beta diversity analysis.

***Table S13***

Table S13 Zeta diversity model specification and decay pattern.

***Table S14***

Table S14.1 Input environmental matrix for gradient forest models.

Table S14.2 Input biological matrix for gradient forest models.

***Table S15***

Table S15.1 Variable importance from 20 replicated gradient forest runs.

Table S15.2 Gradient forest model's goodness-of-fit for each family.

***Table S16***

Table S16.1 Co-occurrence network result co-analyzing all metabarcodes.

Table S16.2 Input measurements for studying the relationship between gradient forest R^2 and network statistics.

#### **Supplemental Methods**

Below are the full methods for Lin et al. Supplemental information has been merged with the main text methods for continuity and clarity for the reader.

##### ***Sampling design***

We aimed to sample biodiversity from a wide variety of habitats across the state of California using directed volunteers and eDNA metabarcoding. CALeDNA scientists recruited volunteers to perform sampling via social media and direct communication (see ucedna.com). Volunteers were asked to select sample locations based on the variety in habitat profiles they could observe during their hikes and to collect both soil (terrestrial samples with any ratio of sand to clay and decomposed organic matter; leaf and other litter were instructed to be avoided) and sediment (submerged deposits in aquatic environments). Volunteers collecting coastal samples systematically sampled swash, dune, and estuarine areas along the coast. Some sampling trips were organized as 'bioblitzes' and facilitated by CALeDNA staff or collaborating scientists, where additional environmental metadata was obtained. As a University of California (UC) research project, sampling permits were not required for UC Natural Reserve System (UCNRS) lands. For non-UCNRS areas, we obtained permission from California Fish and Wildlife to include eDNA on existing permits. Verbal permission was granted from the Santa Monica Mountains Conservancy and city parks.

Sampling steps were guided by a smartphone webform (Meyer et al. 2019) made in Kobo Toolbox (kobotoolbox.org). Surface samples were collected by filling 2 mL tubes with substrate from $< 2$ cm depth in three biological replicates, each 30 cm apart. Volunteers changed gloves between sampling sites, and avoided touching samples by using a rock or stick from the area to remove large debris when necessary. Samples were frozen at -80 °C immediately upon their return to CALeDNA headquarters at UC Los Angeles.

To minimize the potential effect of seasonal variations in eDNA profiles, we selected samples from March 2017 to July 2017. We classified the predominant biome type of each sample by visualizing the collection geolocation in Google Maps ortho-imagery with input from location descriptions in volunteers' reports, National Land Cover Database (Homer et al. 2015) and community observation records from iDigBio (https://www.idigbio.org/) and iNaturalist (https://www.inaturalist.org/). This information was summarized as the *transect* variable in the sample metadata (Table 1; Table S1). We selected 100 samples from each of three *transect* types (coast/coastal, shrub/ShrubScrub, and forest) that covered the broadest latitudinal range possible. Samples with ambiguous metadata were removed, resulting in a total of 278 samples (98 coast, 89 shrub and 91 forest) used in subsequent analyses.

***Compilation of environmental variables***

We assembled environmental variables across six main categories: location, habitat, bioclimate, topography, human impact and vegetation (Table S1). Sampling site latitude, longitude, GPS resolution, site photographs, and collection notes from the Kobo webform were used to assign environmental metadata.

For habitat assignment, we extracted values at each sampling point from shapefiles in EPA Level III Ecoregions of California (U.S. Environmental Protection Agency 2012). Major and minor habitat types for each site were assigned by evaluating photographs together with location and following the California Wildlife Habitat Relationships System (CWHR), with additional categories from the *Revision to Marine and Estuarine Habitats of the CWHR System* (Mayer and Laudenslayer 1988, Shaffer 2002, California Department of Fish and Game 2005;

Text S1). For bioclimatic variables, GeoTiff files for 19 standard BioClim variables at 30 s resolution were downloaded from http://worldclim.org/version2 (Fick and Hijmans 2017). For soil properties, GeoTiff files at 0 m soil depth and 250 m resolution were downloaded from https://soilgrids.org/ (Hengl et al. 2017). Topography features (such as elevation and slope) were extracted from the Elevation Derivatives for National Applications database (U.S. Geological Survey 2005). For human impact features, the human footprint (*hfp*) map was downloaded from https://doi.org/10.7927/H4M61H5F (WCS and CIESIN 2005). For surface reflectance and vegetation indices, GeoTiff files for 13 Sentinel bands (from 2017-01-01 to 2018-07-01; after standard cloud filtering), LandSat8 derived indices (from 2017-04-01 to 2017-06-15) at each site were downloaded at 100 m resolution on Google Earth Engine (Gorelick et al. 2017). For detailed methods and description, please refer to Table S1.2 and Text S1.

All raster layers were aligned and projected to a unified 100 x 100 m grid from Google Earth Engine (Coordinate Reference System for this project: ESPG 4326, WGS84). Layers with a higher original resolution were down-sampled using a mean aggregation method. Layers with a lower or same original resolution were projected using bilinear method for continuous values and nearest neighbor method for categorical values. Layers were then stacked and clipped to California's extent. Raster values for each sampling site were extracted by geographical coordinates. For coastal sites outside of the raster's geographical coverage, values were extracted by the closest point available in 0.5 km radius. If the closest point fell outside of the radius, that site was assigned "NA" value. Unless otherwise specified, all computation and analyses were performed in R version 3.5.3 (R Core Team 2019). Raster operations were performed using R package *raster* (Hijmans 2017).

Considering that many environmental variables are correlated, we evaluated the Pearson's correlation coefficient of the 56 numerical environmental variables and hierarchically clustered the variables according to the coefficients into variable groups using R functions *cor*, *hclust* and *cutree*. To reduce collinearity and improve interpretability in community modeling, we created a reduced set of 33 numerical environmental variables that had an $R^2 < 0.8$ (Table 1) for downstream analysis.

***DNA extractions, amplification and sequencing***

DNA extraction, amplification and sequencing followed Curd et al. 2019. Briefly, three biological replicate soil samples from each site were fully homogenized and pooled. 250 mg of pooled sample was extracted using QIAGEN DNeasy PowerSoil Kit (Qiagen, Valencia, CA, USA) according to the manufacturer's instructions. Negative controls were included in every batch of 12-18 extractions. DNA was amplified by polymerase chain reaction (PCR), using primers for five barcode regions: *16S* (515F and 806R; Caporaso et al. 2012), *18S* (Euk_1391f and EukBr; Amaral-Zettler et al. 2009), *CO1* (mlCOIintF and Fol-degen-rev; Yu et al. 2012, Leray et al. 2013), fungal *ITS1* ("*FITS*"; ITS5 and 5.8S; White et al. 1990, Epp et al. 2012), and plant *ITS2* ("*PITS*" ITS-S2F and ITS-S3R; Gu et al. 2013). For samples belonging to the coast transect, they were additionally amplified using *12S* barcode targeting fish (MiFish; Miya et al. 2015) for another study. Although the result is not described here, the raw sequence file contains those sequences.

Primers were modified to have Nextera Transposase Adapters at the 5' end of original sequence (primer sequence and thermocycling profiles in Table S2.1-2). All PCR amplifications were performed in triplicate and with additional PCR negative controls. Positive amplifications were confirmed by gel electrophoresis. Triplicates were then pooled by sample and barcode.

Pooled amplicons were quantified using a Qubit dsDNA BR Assay Kit, and equal molecules of each amplicon type were pooled by sample. Sample DNA libraries were uniquely double-indexed using Nextera Index A and D Kit, and sequenced with the Illumina MiSeq v6 platform for 2x300 bp reads (QB3-Berkeley FGL; University of California, Berkeley, CA, USA) with a target sequencing depth of 50,000 reads/sample/metabarcode. Five of the 278 sites were processed as biological replicates by different technicians to inspect taxonomic variation in independent DNA extraction and technical processing.

***Bioinformatics and data processing***

We used default settings in the *Anacapa* Toolkit (Curd et al. 2019) for multi-locus sequence data processing and taxonomy assignment. In brief, quality control of raw sequences was performed using *Cutadapt* (Martin 2011) and *FastX-Toolkit* (Gordon et al. 2010), and inference of Amplicon Sequence Variants (ASVs) was made with *DADA2* (Callahan et al. 2016). Taxonomy assignment was made on each ASV using *Bowtie2* (Langmead and Salzberg 2012) and the Bayesian Lowest Common Ancestor algorithm (*BLCA*; Gao et al. 2017) on custom metabarcode-specific reference databases, created using *Creating Reference libraries Using eXisting tools* (*CRUX*; Curd et al. 2019). *Bowtie2* first aligns ASVs against the corresponding metabarcode-specific reference database and returns up to 100 alignments for each ASV. *BLCA* then determines the lowest common ancestor (LCA) from *Bowtie2* hits for each ASV and assigns a bootstrap confidence for each level of the taxonomic path. Taxonomy assignments with a bootstrap confidence cutoff score over 0.6 were kept for each ASV. ASVs with the exact same inferred LCA passing confidence filter were summed into one taxonomic entry as the species/phylotype/MOTU equivalent in this study and will be referred as “taxonomic entry” in the following text.

To informatically control for contamination, we further removed all singleton or doubleton ASVs, and ASVs that occurred more than or equal to three times in all blank samples from subsequent analyses (Table S3). Resulting ASVs from each primer were converted to *phyloseq* objects using the R package *ranacapa* (Kandlikar et al. 2018).

Within each metabarcode, sequences for different sites were rarefied to the same read depth to account for inequities in sequencing depth variations. This was performed with the *custom_rarefaction* function in the *R* package *ranacapa* with 10 replications (Table S2.3). Reads with no assignment were not removed before rarefaction. The rarefaction depth was selected to include as many samples as possible while exceeding the exponential and linear stages of the taxon accumulation curve (Text S2). Sites that did not reach the required depth were dropped from the rarefied dataset (Table S2.3). The rarefied dataset was used in subsequent alpha and beta diversity analyses only, and other analyses used the unrarefied datasets.

To evaluate how material aliquoted for DNA extraction and independent lab processing influenced taxon profiles, we estimated concordance in the decontaminated and further rarefied datasets between biological replicates (Text S3).

***Comparisons with traditional surveys***

To compare the eDNA taxonomic results to traditional surveys, we compared eDNA results to the curated species inventory of the University of California Natural Reserve System (UCNRS), which records Chordata, Arthropoda, and Streptophyta. The UCNRS boundary database from https://ucnrs.org/gis-database/ was used to classify the transect sample sites as "within UCNRS" if they were recorded as one of the UCNRS location and within 1 km range of the corresponding UCNRS boundary. Chordata, Arthropoda and Streptophyta phyla were selected for comparison since only these phyla were recorded in UCNRS inventory. We counted how many taxon records were shared or unique to eDNA results or traditional records (UCNRS) at classification levels of order, family and genus combining all reserves and within each reserve.

We then developed a metric of traditional observation score (TOS) in eDNA taxonomic assignment. TOS uses all species observation and collection records in the Global Biodiversity Information Facility (GBIF) database from a broad region centered on California to score whether the taxon assignment of an eDNA ASV has been observed. A TOS > 0 suggests there is support for the assignment of an ASV based on its presence in the TOS region. The TOS region we selected is a polygon spanning the entire Western United States, Southwest Canada, and Eastern Pacific Ocean (until Hawaii), with coordinates (-155.16652° 20.94569°, -102.11362° 20.93745°, -101.74438° 51.23835°, -155.16935° 51.33748°, -155.16652° 20.94569°). GBIF records were downloaded on January 1, 2020 and the dataset is permanently available through https://doi.org/10.15468/dl.3j4tgy (Gbif.Org 2020). We calculated the TOS for family, genus, and species listed in a concatenated list from all the metabarcode results. Score assignment was weighted by classification level, with Family=1, Genus=2, and Species=4. If a taxon was only resolved to order or higher classification, it was not scored. TOS were 'adjusted' by dividing the TOS by the classification level they were resolved to in eDNA results, using the same weights. This adjusted TOS ranged from 0 to 1. Species entries that were not Latin binomials e.g. "Aphyllophorales sp. EXP0530F" were counted at the genus or family levels when these were specified or were not given a TOS assignment.

***Alpha diversity***

Alpha diversity was calculated using Observed and Shannon's Diversity Index in R package *vegan* (Oksanen et al. 2019). Alpha diversity was binned by nine categorical environmental variables (location, ecoregion, major habitat, minor habitat, initial transect assignment, substrate type classification, sample clusters within 1 km distance, USDA soil major class, national land coverage classification; Table 1). Alpha diversity was only plotted for categorical values with at least five sampling sites. The significance level was set at 0.05. The difference between each category was tested using the Kruskal-Wallis Test with Bonferroni correction for multiple testing. *Post hoc* analysis was performed using the Dunn Test if the Kruskal-Wallis Test was significant (function *dunnTest* in R package *FSA*; Ogle et al. 2019).

We evaluated the relationships of alpha diversity measures and the reduced set of 33 continuous environmental variables (Table 1) as well. First, we performed individual linear regressions (`alpha_diversity ~ variable`) for all combinations of metabarcode, alpha diversity measures (Observed/Shannon's Index) and environmental variables using function *lm* in R. To provide a more complete evaluation and account for the collinearity in the 33 variables, we also used partial least square (PLS) models with methods adapted from Lallias et al. (Lallias et al. 2015, George et al. 2019). PLS models are designed to handle regressions that have many, possibly correlated, explanatory variables and relatively few observations. A PLS model will first project the predictor variables onto a number of orthogonal principal components (or latent variables) and then perform the linear regression on these components to reduce the effect of multicollinearity (Lallias et al. 2015, Mevik 2019). We fit PLS models for alpha diversity in each metabarcode using function *plsr* in R package *pls* (Mevik et al. 2019) with three components included in the model (ncomp = 3). The $R^2$ for PLS models was calculated by $R^2 = 1 - \frac{SSE}{SST}$, where SST is the (corrected) total sum of squares of the response, and SSE is the residual sum of squares (Mevik et al. 2019). The variable importance in projection measure was calculated using function *VIP.R* in R package *pls* extension (https://mevik.net/work/software/VIP.R).

***Beta diversity***

Unless otherwise specified, all functions mentioned in this section are from R package *vegan*. Community composition was visualized by plotting sample relative abundance of the top ten phyla for metabarcodes *16S*, *18S*, and *CO1*, and top ten classes for *PITS* and *FITS*, partitioned by the major habitat. This was implemented by function plot_taxa in R package *microbiomeSeq* (Ssekagiri et al. 2017).

Composition was analyzed using unconstrained ordination. We calculated the binary Jaccard dissimilarity distance from the rarefied dataset for each metabarcode dataset and performed principal coordinate analysis (PCoA), plotting the first two principal coordinates. Permutational multivariate ANOVA (PERMANOVA) analysis was used to determine if the centroids of the calculated binary Jaccard dissimilarity distance differ within each of the nine categorical environmental variables. PERMANOVA was implemented by function *adonis* with 2999 permutations. We excluded categories in the variable if there were less than 5 sampling sites in that category. We also tested for the assumption of homogeneity of dispersion using the *betadisp* function. *Permutest* function was used to test for the significance, with 2999 permutations. P values were adjusted using the Bonferroni method for both PERMANOVA and *betadisp* tests. The significance level was set at 0.05.

The effect of spatial correlation and the large variation in aquatic/coastal sites could confound with PERMANOVA analysis. To control for the effect of spatial correlation, partial constrained analysis of proximities (CAP) was done for the habitat and soil property variables while removing the effect of geographic coordinates (an example formula: `dissimilarity ~ majorhab + Condition(Longitude + Latitude)`. The partition of variance was evaluated using the *varpart* function. The significance of the model was tested using 999 permutations implemented by function *anova.cca*. The significance level was set at 0.05. To control for the large variation in aquatic/coastal sites, PERMANOVA was repeated by excluding the samples from the coastal sites. We also partitioned the data by the four categories in the *majorhab* variable (aquatic; herbaceous; shrub and tree dominated habitats) and performed PCoA analysis within each major habitat. We evaluated the source of variation related to minor habitat using PERMANOVA as described above with the exception that minor habitat categories with less than 5 sampling sites were not removed.

*Post hoc* explanation of the ordination axes was performed by fitting the reduced set of numerical variables (Table 1) onto the PCoA result using the *envfit* function. The significance of each variable was tested based on random permutations (n = 1999) of the data. We additionally fitted the surface of each variable onto the ordination using the *ordisurf* function.

***Zeta diversity***

In addition to the classic alpha and beta diversity analyses, we investigated relationships between environmental variables and eDNA biodiversity turnover utilizing zeta diversity. Zeta diversity is a generalized extension of alpha and beta diversity (Hui et al. 2014) which allows comparisons to be made between the compositions of any number of sampled communities. Using this framework has allowed for novel investigations of the ecology of introduced species (Leihy et al. 2018), community assembly processes (Latombe et al. 2017) and environmental monitoring (Simons et al. 2019).

To measure the fraction of unique categories of organisms held in common between any four sampled nearby communities, we calculated zeta four diversity ($\zeta_4$). We chose four as our sampling order because most reserves were sampled fewer than six times by a single CCS team. Where more than four sites were available, clusters of four sites were made by randomly selecting from within a geographical area in which samples were no more than ~10 km apart (the typical hiking range for a CCS team). The value of $\zeta_4$ was scaled for each geographic cluster as a fraction of its average taxonomic richness per sample ($\zeta_1$) using the function *Zeta.decline.ex* in the R package *ZETADIV* (Latombe et al. 2018). We tested the likelihood of two models of community assembly, using the Akaike Information Criterion (AIC) score within *ZETADIV*, through calculations of how zeta diversity decays with sampling order. Based on prior analyses (Hui et al. 2014) decays which follow a power-law of the form $\zeta_N = \zeta_1 N^{-b}$, or an exponential of the form $\zeta_N = \zeta_1 e^{b.(N-1)}$, were associated with a niche differentiation or stochastic process of community assembly, respectively. Scaled $\zeta_4$ diversity values were then plotted on a map of California using the R package *Leaflet* (Cheng et al. 2019).

Environmental factor groups were made by binning environmental variables according to their categories (Table 1). To determine the variation in $\zeta_4$ diversity attributed to either geographic distance or an environmental factor group, we used the function *Zeta.varpart* to analyze generalized linear models (GLM) generated using the function *Zeta.msgdm* within the R package *ZETADIV*. For each GLM we only used samples where both the geographic cluster ID and particular environmental factors were known.

***Gradient forest modeling***

We used the gradient forest classification model in R package *gradientForest* (Ellis et al. 2012) to test which environmental variables best explained eDNA-detected taxon presence patterns across California using all 272 sites collected from three transects. Due to large variation in the coastal sites, we also performed additional analyses excluding all coastal sites using the same methods described below.

We first prepared the input biological matrix. To include results from all five metabarcodes, we merged the ASV tables from five metabarcodes using the *merge_phyloseq* function in R package *phyloseq* and this table will be referred as "merged decontaminated taxonomy table". From the merged decontaminated taxonomy table, taxonomic entries were further agglomerated to family level (using custom R function *glom_tax*) to minimize stochastic errors in the metabarcoding data generation and to increase interpretability in downstream analysis. Taxonomic entries not resolved to family level were removed. Read counts were converted to presence/absence records when there were at least three reads at each site. We excluded any families that occurred in less than 2.5% or more than 97.5% sites from downstream analysis because these extremes would inflate false positives. The stability of the response variables was visually inspected by Jaccard dissimilarity PCoA using biological replicates (Text S3).

The gradient forest model was built with the reduced model containing 33 numerical environmental variables (Table 1; Figures S1,S2) that had a correlation coefficient $< 0.8$ with all other variables. We fit a classification-tree based gradient forest model using default settings to the biological matrix derived, but increased the number of trees to 2000 per family to increase the stability of the model (Breiman 2001).

The gradient forest analysis provides a ranked list of all environmental variables based on their relative predictive power of community turnover. It also offers the analysis of split density and the cumulative importance for each environmental variable. The model also ranks the families by their correlation strength with environment and provides a summary of where families occurrence turnovers are distributed along the environmental gradient. We inspected these model products by manually comparing "highly-predictable" families to the original metabarcoding results and by evaluating which families had a low percent identity or low number of matched reads to *CRUX* reference database sequences. We also manually inspected the ASVs within each family and confirmed that multiple reference sequences were used in *BLCA* inference that determined the level of classification we report in results.

To assess model robustness and stability, we repeated the gradient forest model 20 times with the same settings and recorded the number of families with positive $R^2$ values, the overall corrected $R^2$, the importance of each explanatory variable, and the $R^2$ of each family. To assess model power and reliability, we used a permutation approach akin to Bay et al. 2018: we randomized the predictor matrix 100 times and ran the model, recording the number of families with positive $R^2$ values as well as the overall corrected $R^2$ for these 100 runs. We compared these results to the corresponding results from the observed data via histogram.

To visualize the community turnover gradient forest model over space, we used the input of all 33 environmental variables from 100 m x 100 m grids in the State of California (CA). At each grid, the environmental variables were transformed and standardized based on their relative importance in the model without extrapolation according to the gradient forest manual (Pitcher et al. 2011). We note that transforming variables without extrapolation ensures that transformed prediction would not exceed the maximum importance of model prediction, but often this imposes a non-linear transformation for extreme values (Pitcher et al. 2011). Then, we used the top three principal components from the transformed environmental variables and visualized them by red, green and blue (RGB) bands (Ellis et al. 2012). More similar colors on the spectrum represented closer community composition. Each environmental variable loading was visualized by function *biplot* in R.

To differentiate model performance from the innate high-dimensional nature of the environmental variable matrix, we scaled the environmental variables and performed the same PCA and visualization procedure without using the model ("uninformed map"). Both the model-based map and the uninformed map were qualitatively compared with regional maps to evaluate their ability to recapitulate known major biodiversity patterns. The improvement in predictability was further quantified by comparing dissimilarity matrices with methods adapted from Pitcher et al. 2012. Briefly, we performed a mantel test and a monotonic regression between the biological matrix and either the *uninformed* map or *gradient forest informed* map. The correlation coefficient from the Mantel test and the stress diagnostics from the monotonic regression were used as a metric to weigh model performance.

***Network analysis***

Results for each metabarcode were summarized by family and used in ecological co-occurrence network analysis. Prior to calculating networks, results were filtered to only retain families with 10 or more reads in at least 10% of the samples (28 sites). The SPIEC-EASI method was employed using R package *SpiecEasi* (Kurtz et al. 2015) for cross domain analysis that incorporates all five metabarcodes into one complex network (see Tipton et al. 2018) with method 'mb' for Meinshausen-Buehlmann Neighborhood selection, lambda minimum ratio of 1e-2, nlambda = 40, and pulsar parameter threshold of 0.05. Resulting undirected network matrices were imported into *Cytoscape* (v. 3.6.1; Shannon et al. 2003) and *NetworkAnalyzer* was used to calculate topological parameters.

To observe the relationship between network degrees and the prediction $R^2$ of each family from gradient forest, an ordinary least squares (OLS) linear regression model was made:

`Family R2 ~ Network_Sum * Frequency_in_sites` using the *lm* function in R. Interaction plotting was made using the R package *Interactions* (v. 1.1.1; Long 2019).

To evaluate the co-occurrence and gradient forest predictor patterns in a phylogenetic framework, the 915 families used in the gradient forest modeling were mapped onto the Open Tree of Life (tree.opentreeoflife.org) and a synthetic tree was generated using synthesis release v12.3. Datasets were mapped next to the phylogeny tips using the Interactive Tree of Life (https://itol.embl.de/). Scripts used for converting taxonomy to phylogeny are deposited in https://github.com/McTavishLab/OT2020/blob/master/Cal%20eDNA%20families.ipynb.

#### Supplemental Text

##### *Text S1: Habitat designation method*

###### Methods

We classified sample site major and minor habitat according to the California Wildlife Habitat Relationships System (CWHR) with additional minor habitat categories from the revision of marine and estuarine habitats of CWHR relationship system (Shaffer 2002, California Department of Fish and Game 2005). The previous classifications CWHR only included marine, estuarine, lacustrine and riverine categories (Mayer and Laudenslayer 1988). The revision now includes 22 marine and 19 estuarine habitats (Shaffer 2002). Using the key to habitats from Shaffer 2002 we classified the habitats hierarchically as follows: Marine or Estuarine, then Nearshore, and then as Embayment or Coast. Embayment or Coast separation summarizes protection from wave action as protected or exposed, respectively. For this, we plotted the coordinates for each site in Google Earth to determine exposure to wave action in each location. We also used the photos taken from the habitat to classify the habitat as intertidal or shoreline and beach, for example Habitat 3 “Marine Nearshore Exposed Coast Shoreline and Beach”. We used the same approach for the estuaries. We plotted the coordinates in Google Earth to evaluate habitat designation: Bay, River Mouth, Estuary or Tidal Flat. We also used the photos to confirm habitat condition based on sediment (Estuary or Tidal Flat), and the presence of vascular plants (Tidal Marsh).

***Text S2: Evaluation of rarefaction depth***

#### Methods

To account for the sequencing depth variation and discard the samples with low sequencing depth, while retaining as many samples as possible, we performed evaluations of rarefaction depths at 1, 2, 40, 100, 1000, 2000, 4000, 6000, 10000, 15000, 20000 reads/sample for *16S, 18S, CO1, FITS* and *PITS* metabarcodes. We plotted alpha diversity measures against these different rarefaction depths using both Shannon Index and the observed species count (taxon accumulation curve). We also plotted the samples retained against different depths. To provide an estimation of sample coverage loss based on the rarefaction depth we chose, we evaluated the estimated sample coverage before and after rarefaction using R package *iNEXT* (Chao et al. 2014).

#### Results

We found that the rarefaction depths we chose were sufficient to reach the asymptotic phase of Shannon Index accumulation curve (Figure S5). The observed species count displayed a reduced accumulation rate but did not reach asymptotic phase, but the rarefaction depths we chose were sufficient to exceed the exponential growth phase of the species accumulation curve while retaining the most samples (Figure S6; Text S2. Table). The *iNEXT* analysis showed that the estimated sample coverage was not significantly reduced at the chosen levels.

Text S2. Table: Sample coverage before and after rarefaction reported by the *iNEXT* function.

| **Metabarcode** | **Sample Coverage (Before rarefaction)** | **Sample Coverage (After rarefaction)** |
|---|---|---|
| *16S* | 0.998 | 0.979 |
| *18S* | 0.971 | 0.948 |
| *CO1* | 0.955 | 0.916 |
| *FITS* | 0.962 | 0.931 |
| *PITS* | 0.951 | 0.872 |

***Text S3: Concordance in biological replicates***

#### Methods

Metabarcoding results are not expected to be stable in independent extractions due to stochastic processes in different amplification reactions and due to small-scale heterogeneity in environmental substrates (Ficetola et al. 2015, Yamamoto et al. 2017, Alberdi et al. 2018). To examine the extent of this variation in our samples, we analyzed five pairs of replicated samples by repeating DNA extraction and lab processing and then assessed the extent of concordance between the replicates. Methods were adapted from Alberdi et al. 2018. We evaluated the concordance of taxonomic entries occurrences at family, genus, and original lowest common ancestor (LCA) level. The proportion of families that had consistent presence/absence patterns between each replicate pair was reported for the rarefied dataset and un-rarefied dataset at two minimum thresholds (100 reads/sample or 3 reads/sample). We also performed a Jaccard dissimilarity PCoA ordination to evaluate compositional similarity in the context of the remaining 273 samples used in this study. For this analysis, we used a cutoff of at least 3 reads per sample per family, which mirrors the filtering methods we used in gradient forest analyses.

#### Results

Results indicated differences between replicates were persistent at different minimum read cut offs and after rarefaction (Table S4.1-3). We found that the unrarefied results from biological replicates with a three-minimum-read cutoff had an average of 44% of families in common, and 30% of original taxonomic entries are shared between biological replicates. However, when comparing biological replicates while including both presence and absence records from the total taxa in this particular sample set, an average of 93% of families and 97%

of taxonomic entries overlapped (Table S4.2). Despite this variation, we found replicates clustered near each other in the Jaccard PCoA ordination (Figure S4), suggesting compositional similarity remained sufficient to group sites based on compositional community features.

#### Supplemental Figures

***Figure S1. Metadata correlation***

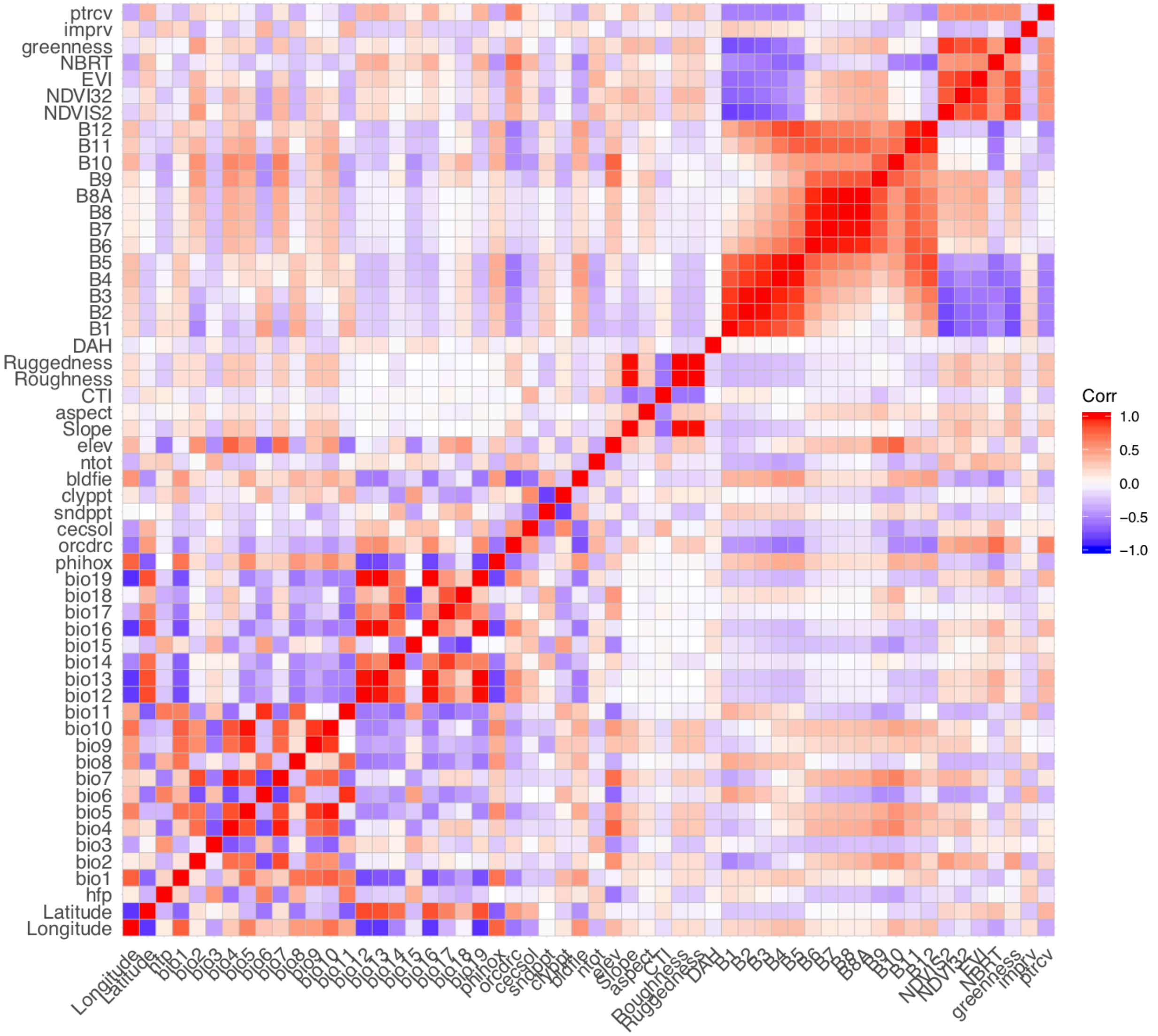

Figure S1 Metadata correlation plot. The heatmap is a plot of Pearson's correlation coefficients for 56 numerical environmental variables. For explanation of variable names, please refer to Table S1.

***Figure S2. Environmental variable distribution***

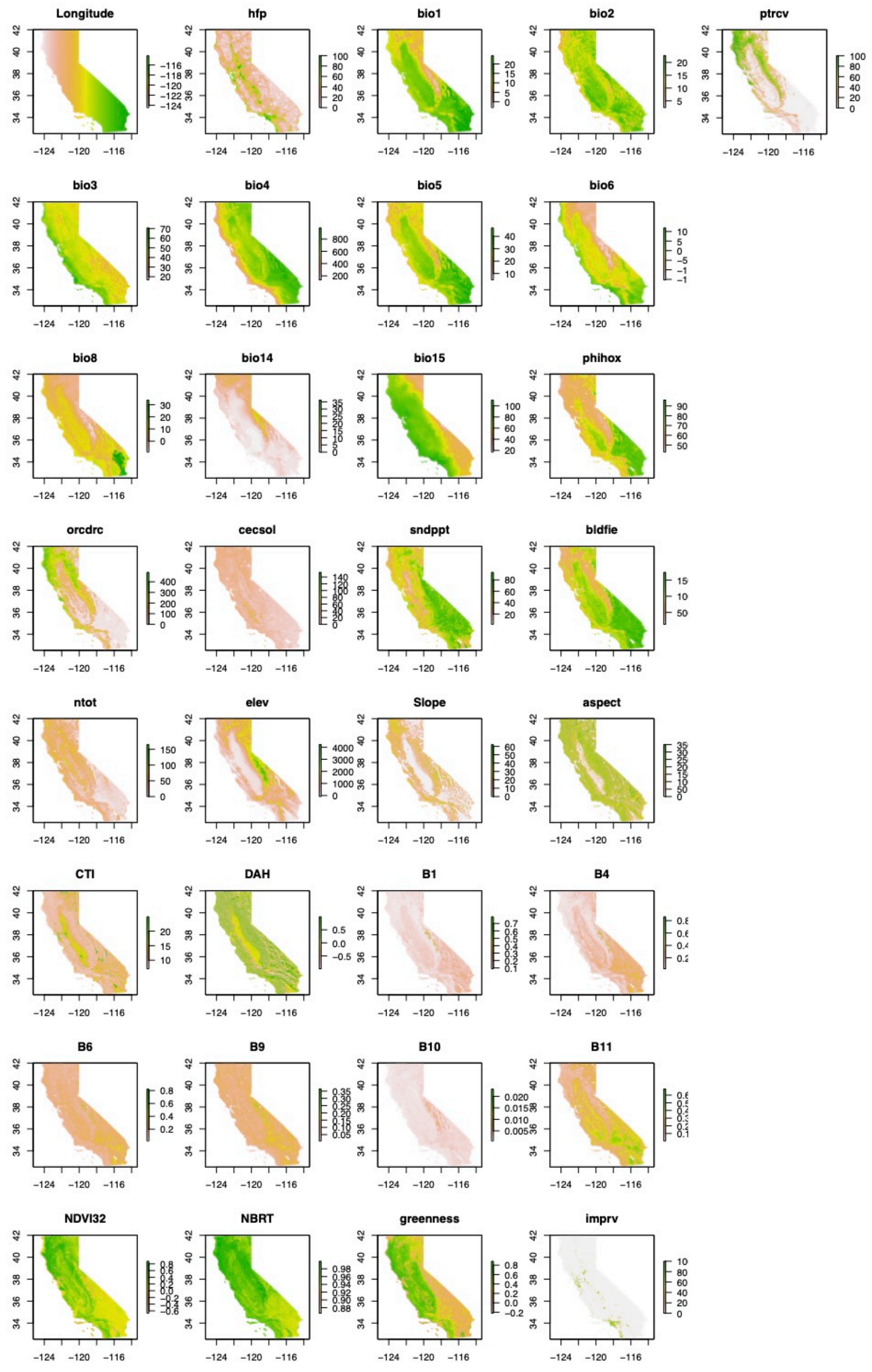

Figure S2 Raster values of the 33 reduced environmental variables (Table 1). Each variable's value distribution is plotted across the state of California at 100 m resolution with a terrain color scheme. White color represents lower values, green medium values and pink higher values.

***Figure S3. Read depth summary***

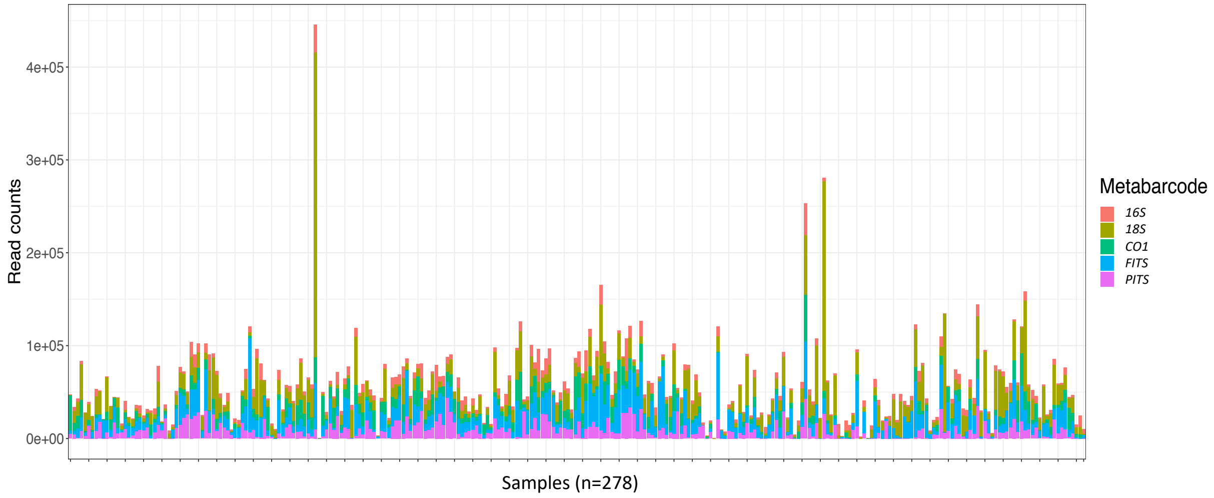

Figure S3 Sequence read depth for individual samples. Read counts are stacked for each metabarcode in the decontaminated taxonomy table (Table S3). Each bar represents one sample site.

***Figure S4. Replicate PCoA***

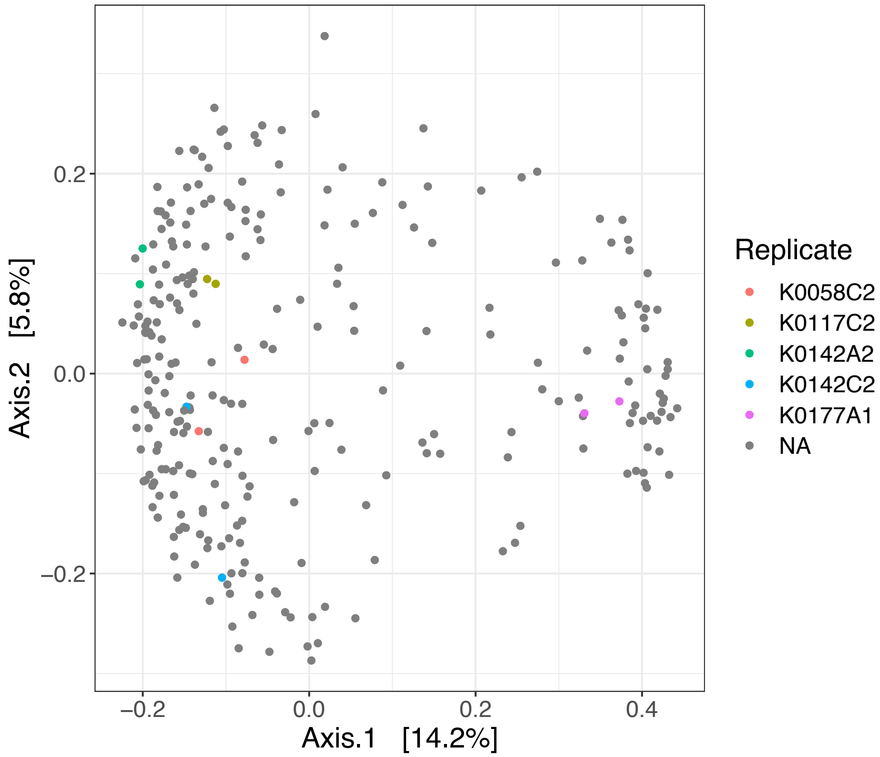

Figure S4 PCoA shows community composition is similar in replicated sample DNA extraction and processing. The sites without replicates were marked in gray as a background distribution. The replicated sites were colored by their sample name.

***Figure S5. Rarefaction summary***

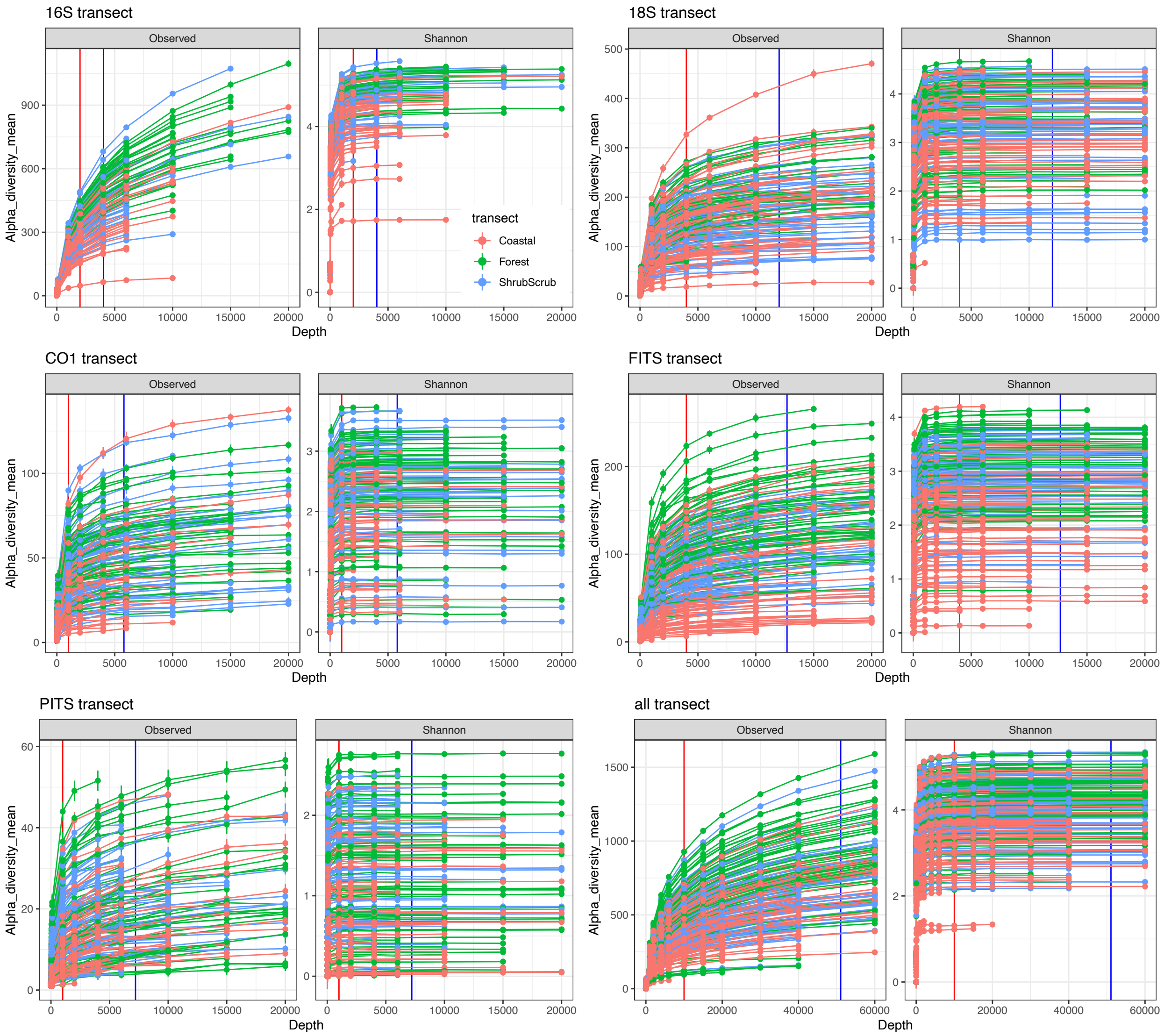

Figure S5 Species accumulation curve generated from the rarefaction in each metabarcode. The red vertical line shows the chosen rarefaction depth and the blue vertical line shows the median read depth. Each accumulation curve stands for one sample, colored by the transect it belongs to.

***Figure S6. Rarefaction and samples kept***

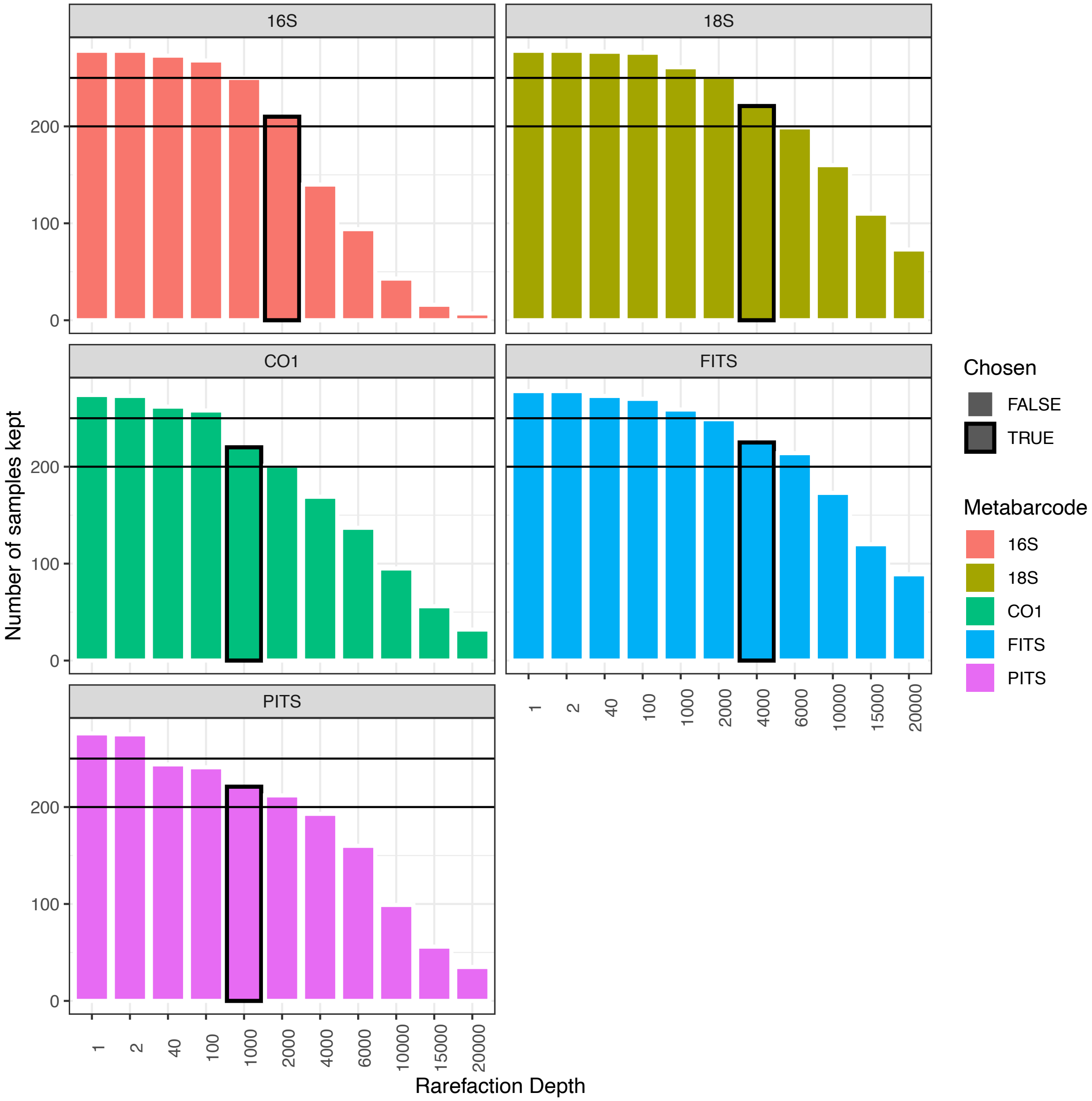

Figure S6 Plot of various rarefaction depths and the number of samples retained. The number of samples that sufficed the rarefaction depth conditions were plotted as a bar plot. The chosen rarefaction depth for this analysis was marked with a black box for each metabarcode. The horizontal lines stand for sample sizes n = 250 and n = 200.

587 *Figure S7. Alpha diversity boxplot: loc*

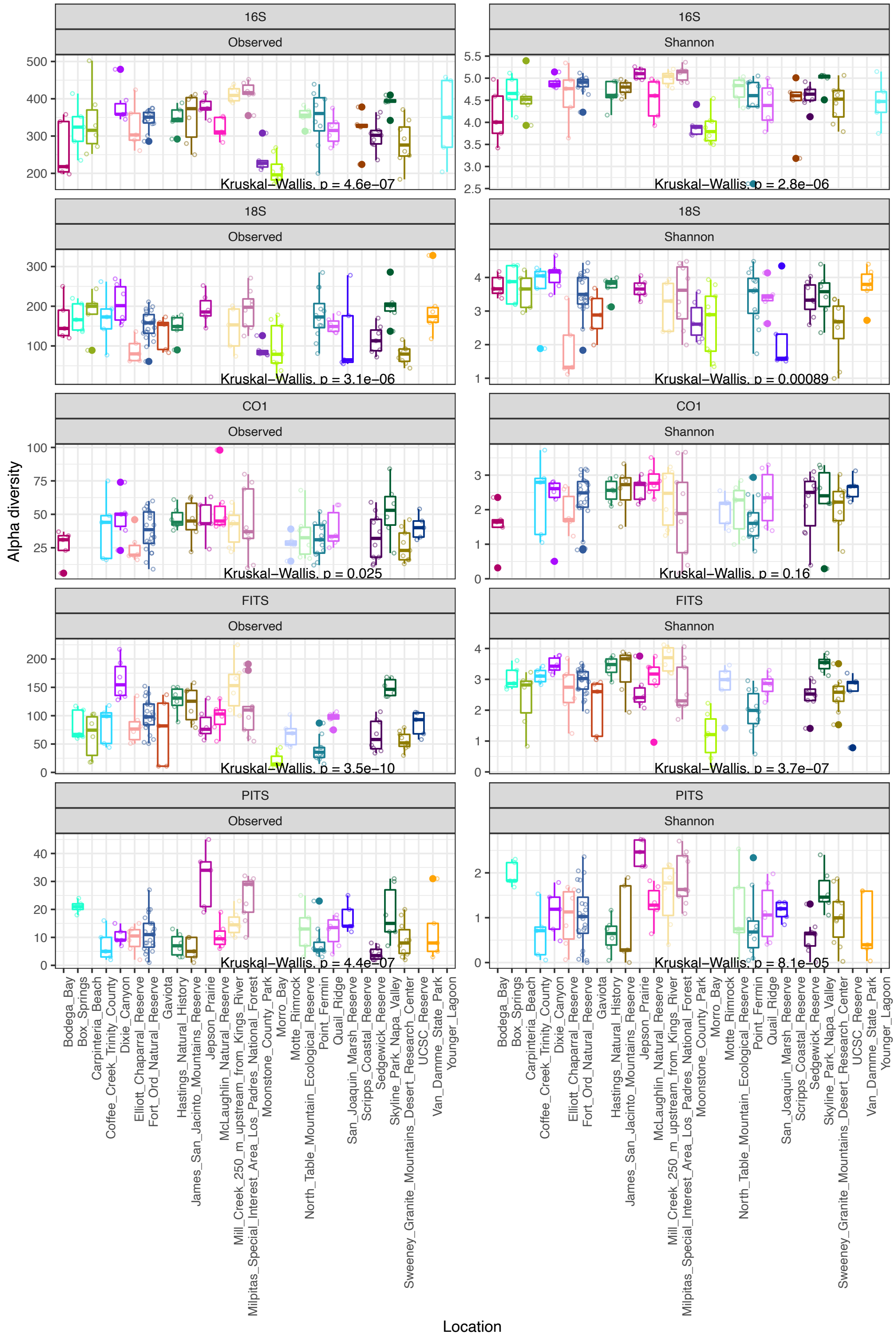

588

Figure S7 Boxplot of alpha diversity measures for five metabarcodes binned by location. Observed or Shannon diversity for rarified *16S, 18S, CO1, FITS*, and *PITS* metabarcodes dataset are grouped by location (*loc*). The box notch stands for the medians. The lower and upper ranges of each box represents the 25th and 75th percentiles. The whiskers extend to data points no more than 1.5 * IQR (inter-quartile range) from the hinges. All data points are plotted additionally. P-values from Kruskal-Wallis tests on mean alpha diversity metrics across categories are denoted at the bottom of each panel. Locations containing less than 5 samples were discarded from the analysis. Locations are sorted in alphabetical order.

***Figure S8. Alpha diversity boxplot: transect***

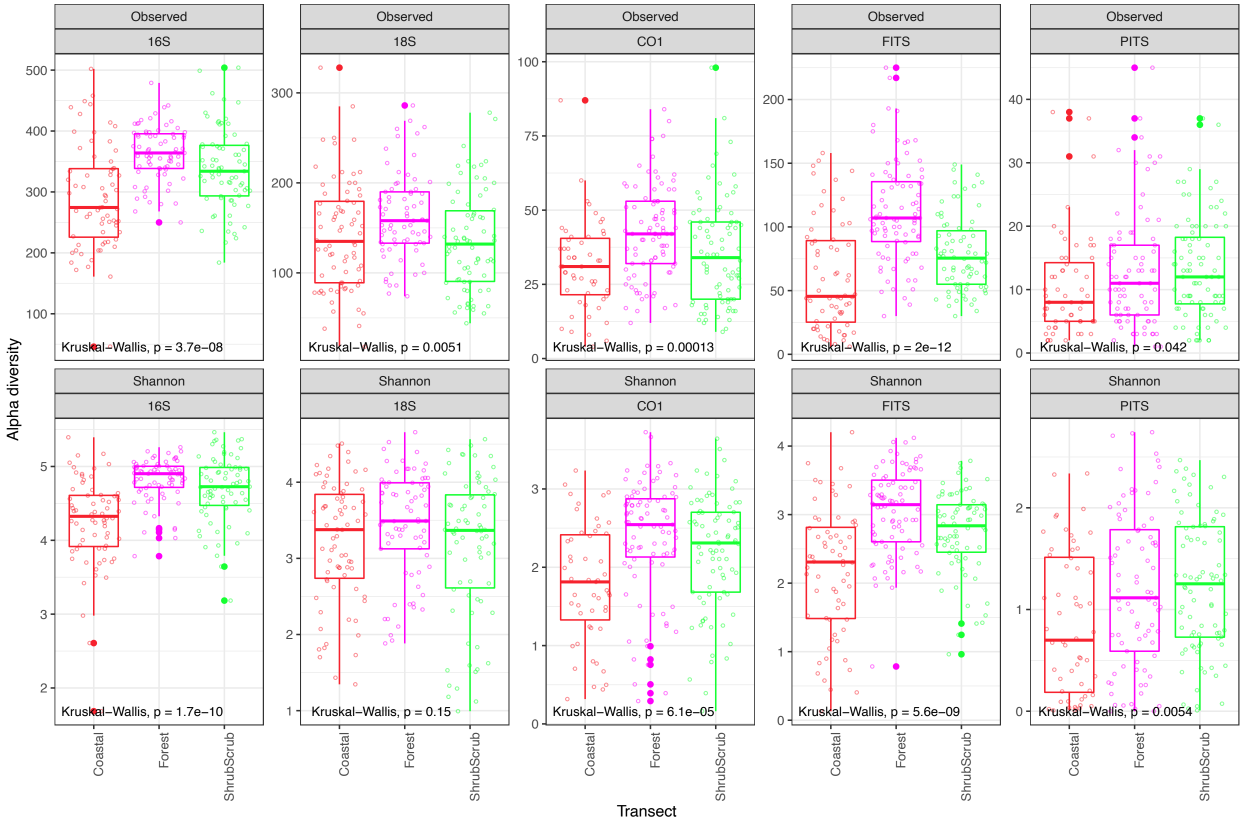

Figure S8 Boxplot of alpha diversity measures for five metabarcodes binned by transect. Observed or Shannon diversity for rarified *16S, 18S, CO1, FITS*, and *PITS*, metabarcodes dataset are grouped by transect. The box notch stands for the medians. The lower and upper ranges of each box represent the 25th and 75th percentiles. The whiskers extend to data points no more than 1.5 * IQR from the hinges. All data points are plotted additionally. P-values from Kruskal-Wallis tests on mean alpha diversity metrics across categories are denoted at the bottom of each panel.

***Figure S9. Alpha diversity boxplot: majorhab***

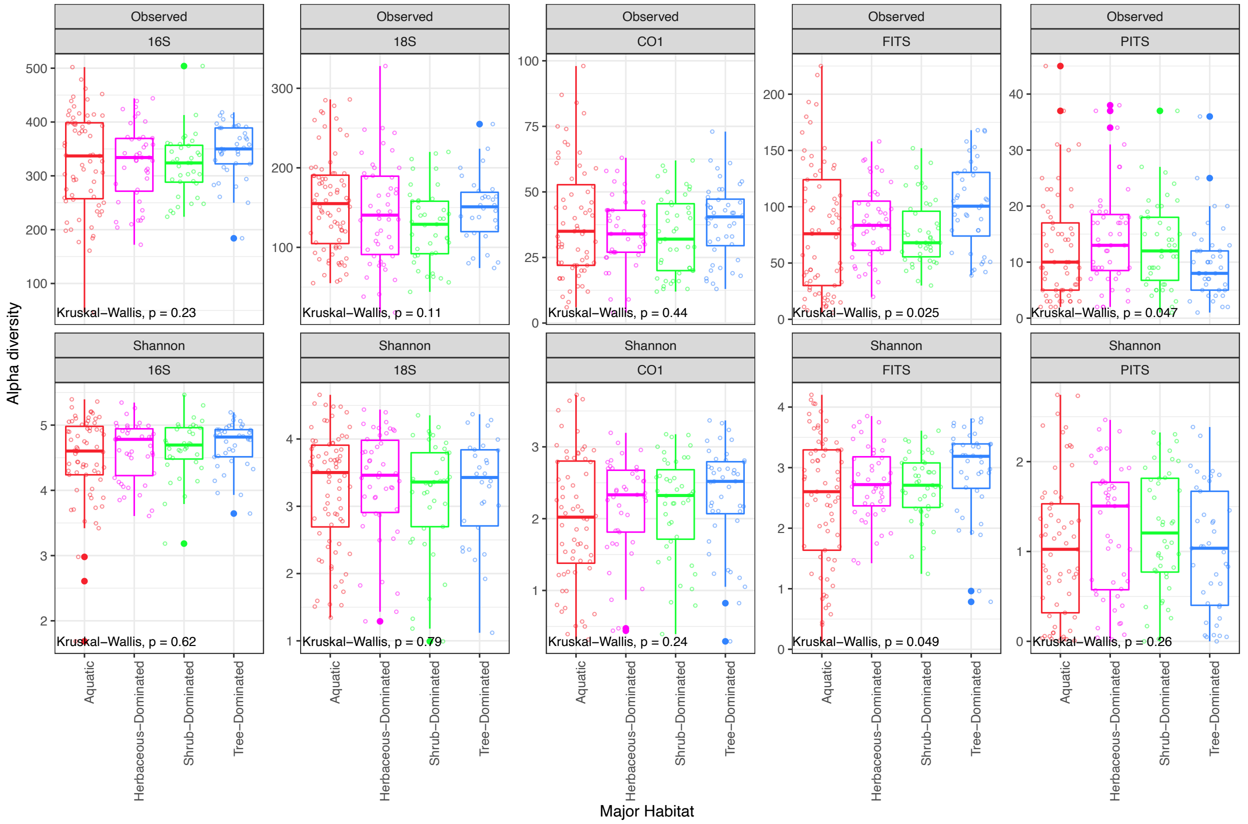

Figure S9 Boxplot of alpha diversity measures for five metabarcodes binned by major habitat. Observed or Shannon diversity for rarified *16S, 18S, CO1, FITS*, and *PITS*, metabarcodes dataset are grouped by major habitat (*majorhab*). The box notch stands for the medians. The lower and upper ranges of each box represent the 25th and 75th percentiles. The whiskers extend to data points no more than 1.5 * IQR from the hinges. All data points are plotted additionally. P-values from Kruskal-Wallis tests on mean alpha diversity metrics across categories are denoted at the bottom of each panel. Major habitat categories containing less than 5 samples and samples missing major habitat classifications were discarded from the analysis.

***Figure S10. Alpha diversity boxplot: NLCD***

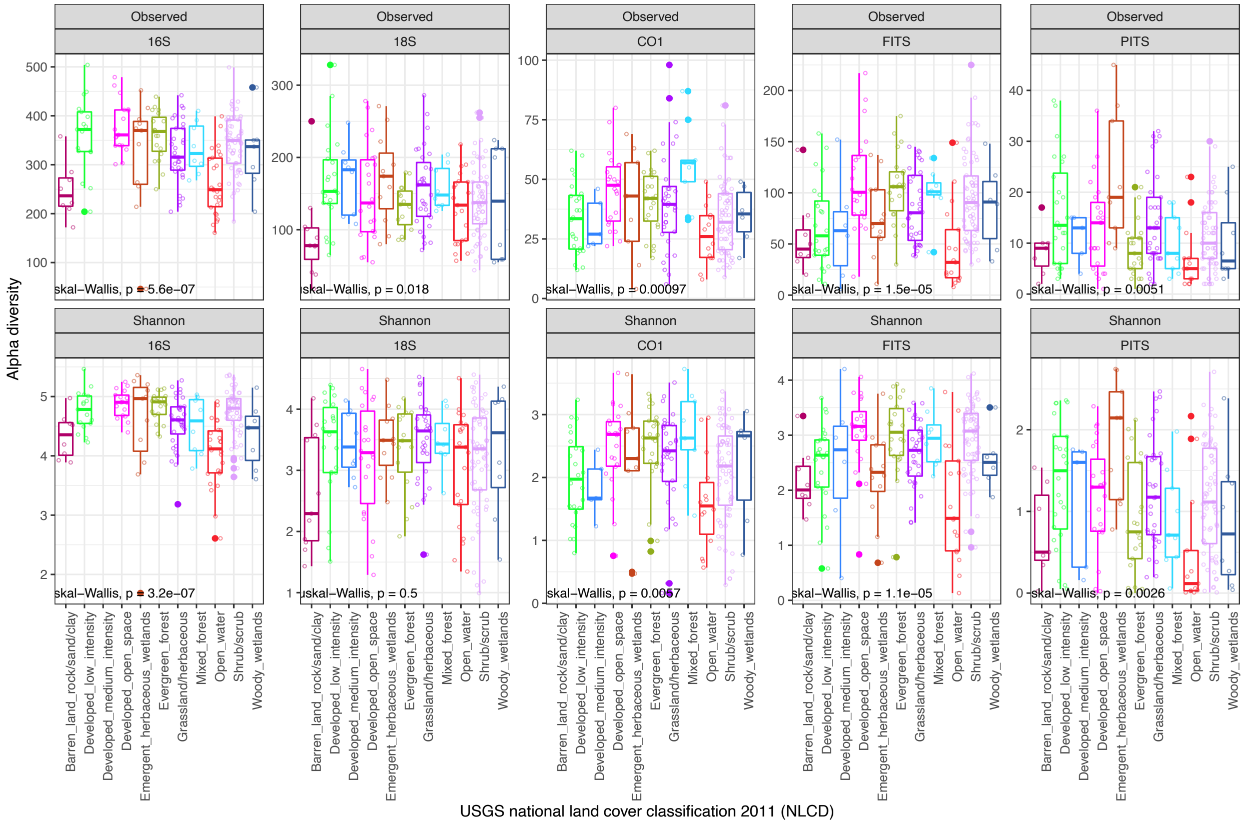

Figure S10 Boxplot of alpha diversity measures for five metabarcodes binned by USGS national land cover classifications (*NLCD*). Observed or Shannon diversity for rarified *16S, 18S, CO1, FITS*, and *PITS* metabarcodes dataset are grouped by *NLCD*. The box notch stands for the medians. The lower and upper ranges of each box represent the 25th and 75th percentiles. The whiskers extend to data points no more than 1.5 * IQR from the hinges. All data points are plotted additionally. P-values from Kruskal-Wallis tests on mean alpha diversity metrics across categories are denoted at the bottom of each panel. *NLCD* categories containing less than 5 samples were discarded from the analysis.

***Figure S11. Alpha diversity boxplot: substrate type***

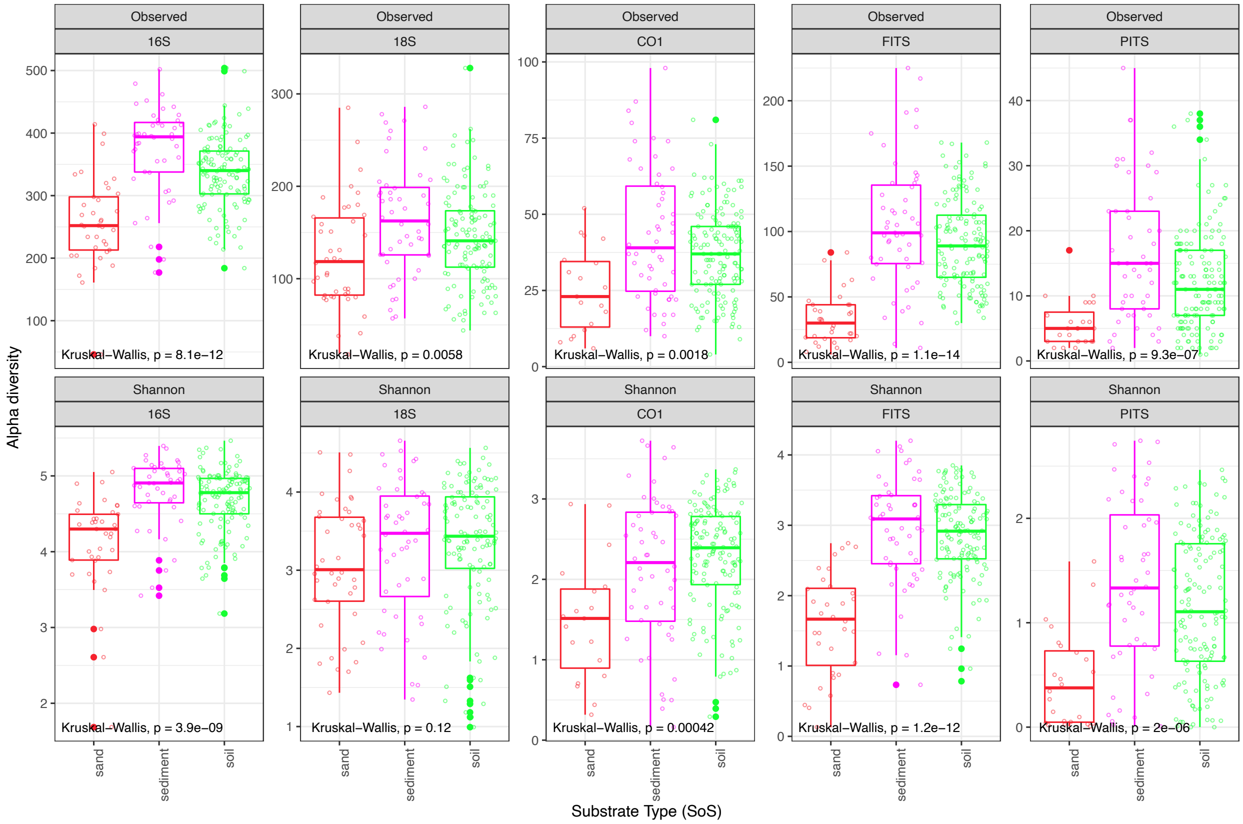

Figure S11 Boxplot of alpha diversity measures for five metabarcodes binned by substrate type (*SoS*). Observed or Shannon diversity for rarified *16S, 18S, CO1, FITS*, and *PITS* metabarcodes dataset are grouped by *SoS*. The box notch stands for the medians. The lower and upper ranges of each box represent the 25th and 75th percentiles. The whiskers extend to data points no more than 1.5 * IQR from the hinges. All data points are plotted additionally. P-values from Kruskal-Wallis tests on mean alpha diversity metrics across categories are denoted at the bottom of each panel. Samples missing substrate type classifications were discarded from the analysis.

***Figure S12. Alpha diversity individual linear model***

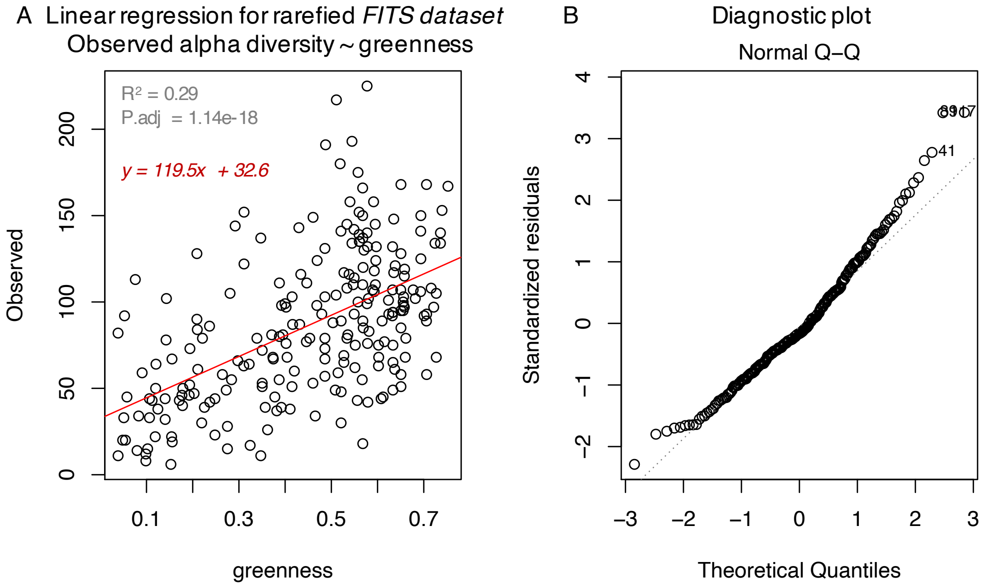

Figure S12 Individual linear regression result and the diagnostic plot (Normal Q-Q plot) on alpha diversity and environmental variables with the highest $R^2$ value. The response variable was the observed alpha diversity in the rarefied *FITS* dataset and the explanatory variable was the *greenness* measurement. The fitted model was plotted as a red line in (A). We also present (B) the Normal Q-Q plot, generated with the *plot.lm()* function in R.

***Figure S13 Taxonomic composition for each metabarcode***

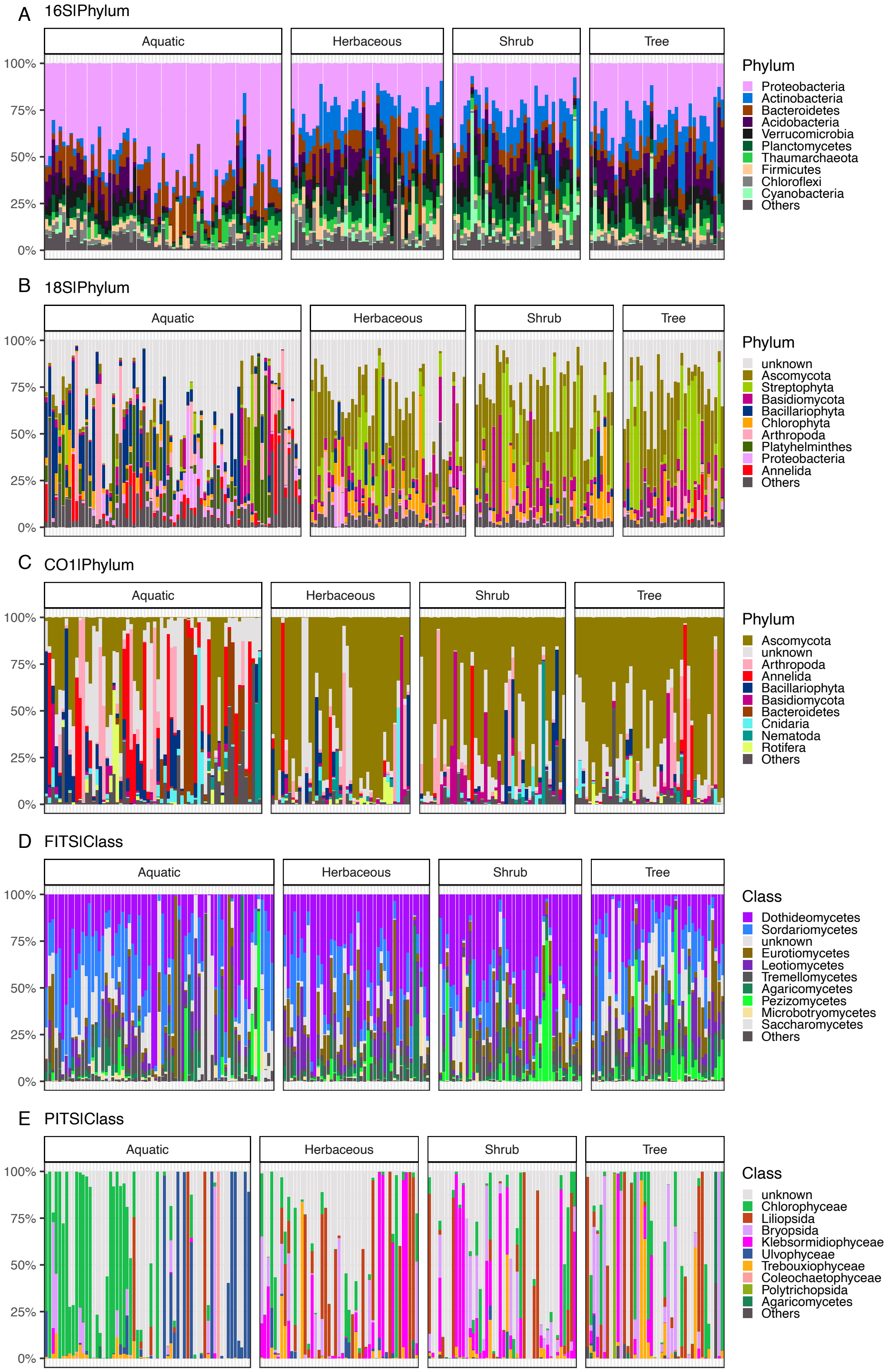

Figure S13 **Taxonomic composition for each metabarcode: (A) *16S*, (B)*18S*, (C) *CO1*, (D) *FITS*, (E) *PITS*.** Decontaminated taxonomic entries were rarefied to the same sequencing depth for each sample within each metabarcode (Table S2.3). The 10 most abundant taxonomic groups are plotted as relative abundance in each sample within metabarcodes: (A - C) are reported at the phylum level and (D - E) are reported at the class level. Each taxonomic group is assigned a unique color. Each bar represents one sample's taxonomic profile. Samples are partitioned by their major habitat classification as Aquatic, Herbaceous, Shrub or Tree-Dominated.

***Figure S14. Beta diversity PCoA: transect***

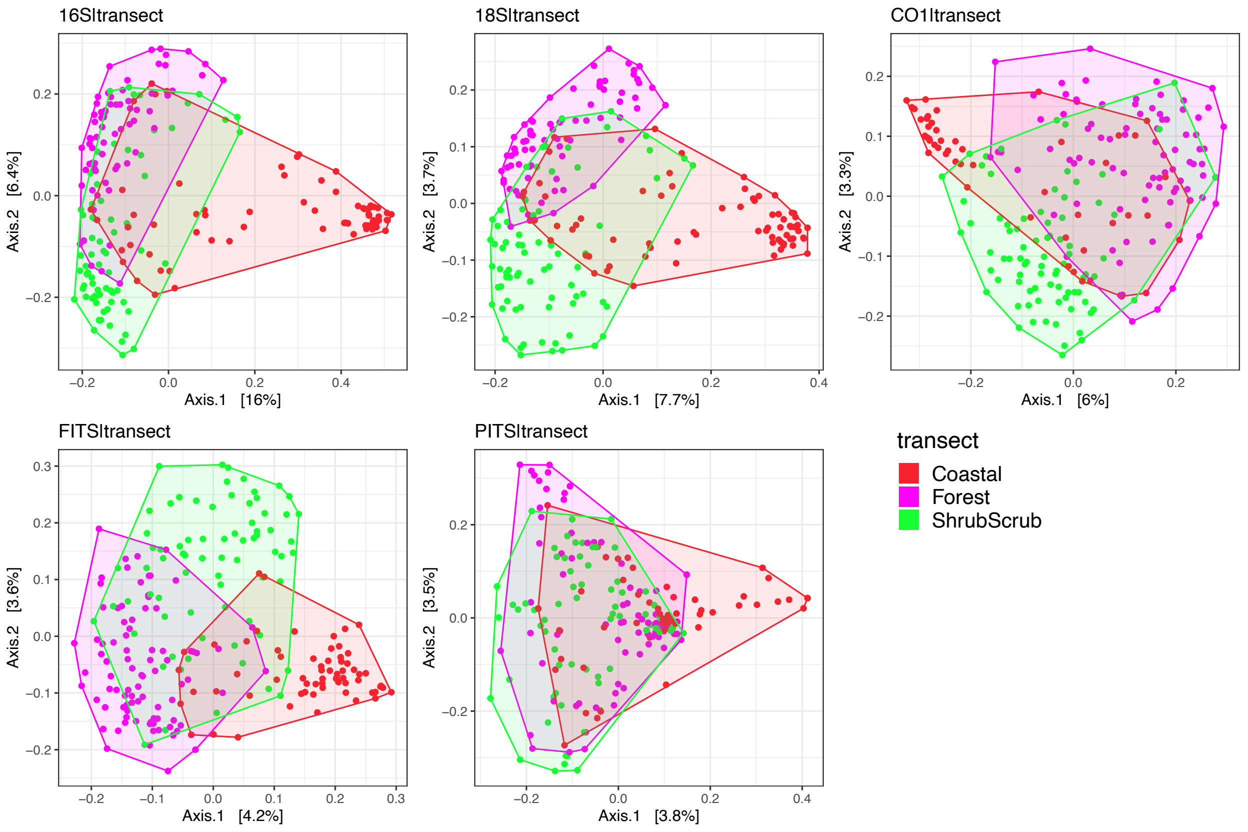

Figure S14 Principal Coordinate Analysis (PCoA) on beta diversity grouped by transect classification. PCoA was performed based on Jaccard dissimilarity calculated from the rarefied dataset. The first two principal coordinates are plotted with percentage of variance explained included in axis label. PCoA derived from *16S, 18S, CO1, FITS,* and *PITS* metabarcodes are plotted individually. Each point represents a sample site and is colored by the *transect* classification it belongs to. Results of *PERMANOVA* and *beta dispersion* can be found in Table S10.1-2.

***Figure S15. Beta diversity PCoA: majorhab***

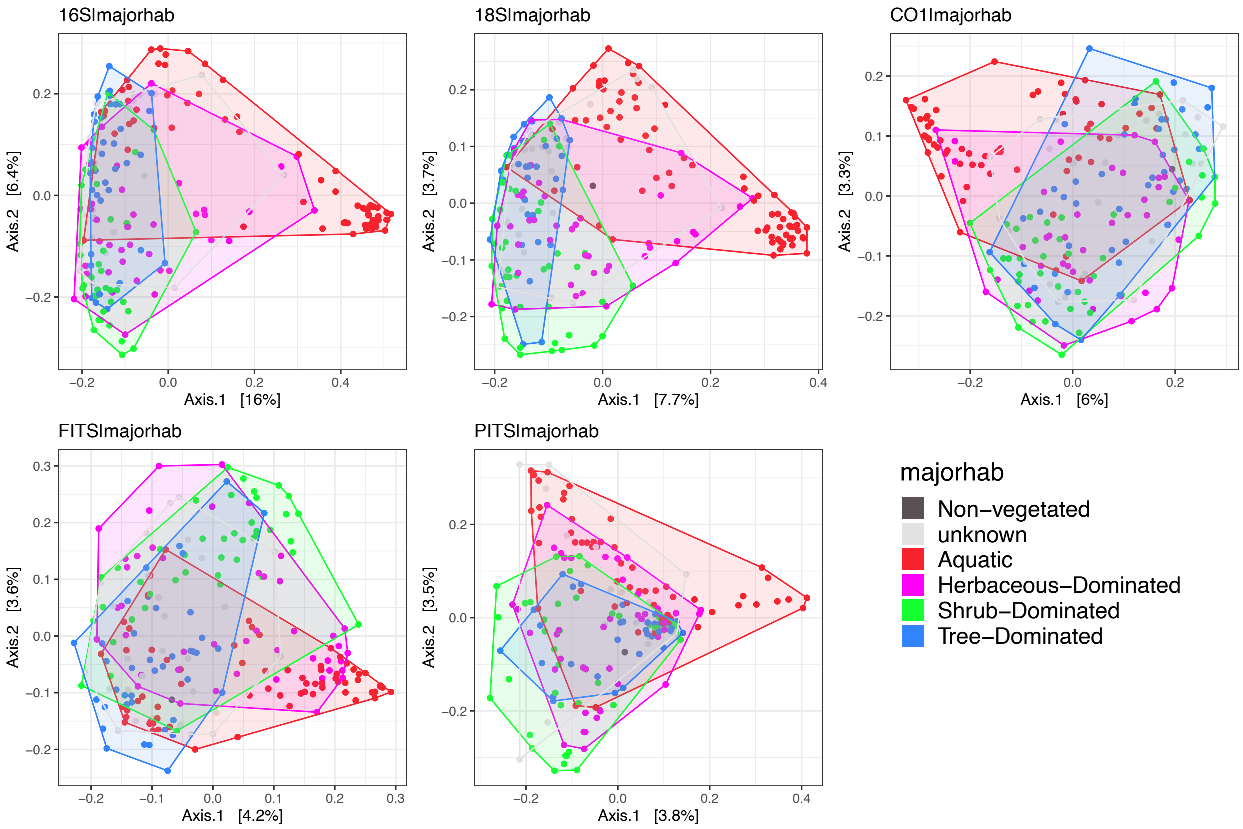

Figure S15 Principal Coordinate Analysis (PCoA) on beta diversity grouped by major habitat classification. PCoA was performed based on Jaccard dissimilarity calculated from the rarefied dataset. The first two principal coordinates are plotted with percentage of variance explained included in axis label. PCoA derived from *16S, 18S, CO1, FITS,* and *PITS* metabarcodes are plotted individually. Each point represents a sample site and is colored by the major habitat classification (*majorhab*) it belongs to. Only one sample was classified as "Non-vegetated" (Table S1.3). Results of *PERMANOVA* and *beta dispersion* can be found in Table S10.1-2.

***Figure S16. Beta diversity PCoA: NLCD***

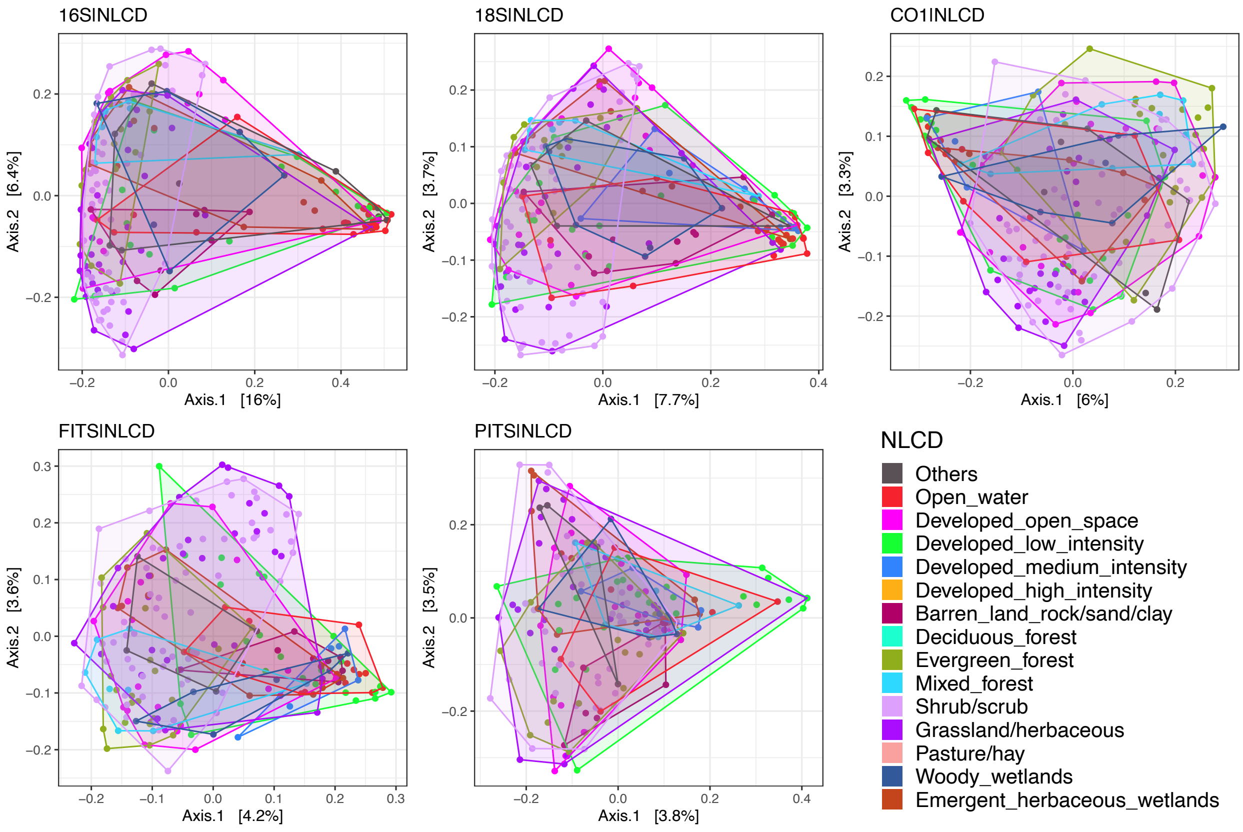

Figure S16 Principal Coordinate Analysis (PCoA) on beta diversity grouped by *NLCD* classification. PCoA was performed based on Jaccard dissimilarity calculated from the rarefied dataset. The first two principal coordinates are plotted with percentage of variance explained included in axis label. PCoA derived from *16S, 18S, CO1, FITS,* and *PITS* metabarcodes are plotted individually. Each point represents a sample site and is colored by the *NLCD* classification it belongs to. Results of *PERMANOVA* and *beta dispersion* can be found in Table S10.1-2. *NLCD* categories containing less than 5 samples in each metabarcode dataset were discarded from the statistical analyses but still plotted on the ordinations, colored as dark gray ("Others").

***Figure S17. Beta diversity PCoA: substrate type***

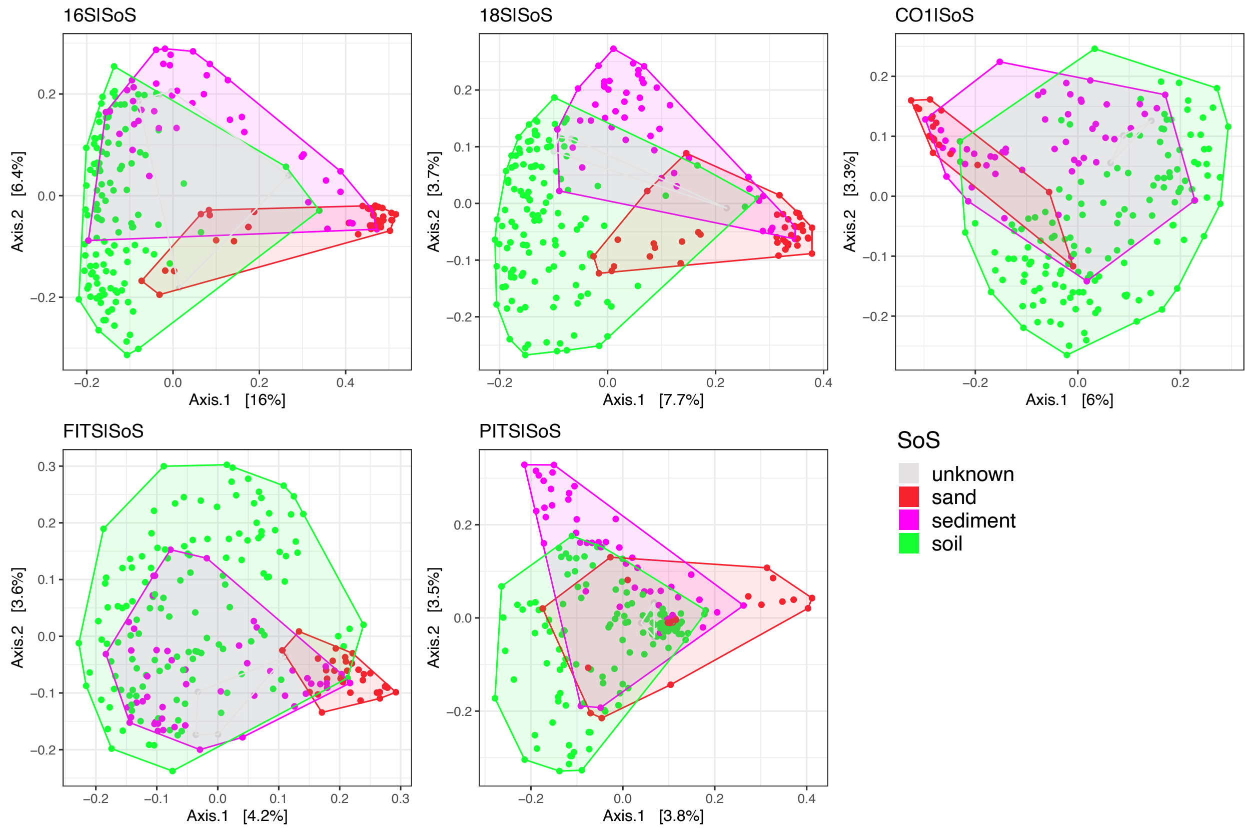

Figure S17 Principal Coordinate Analysis (PCoA) on beta diversity grouped by CALeDNA substrate type (*SoS)* classification. PCoA was performed based on Jaccard dissimilarity calculated from the rarefied dataset. The first two principal coordinates are plotted with percentage of variance explained included in axis label. PCoA derived from *16S, 18S, CO1, FITS,* and *PITS* metabarcodes are plotted individually. Each point represents a sample site and is colored by the substrate type it belongs to. Results of *PERMANOVA* and *beta dispersion* can be found in Table S10.1-2.

700 ***Figure S18. Beta diversity PCoA: no coastal sites***

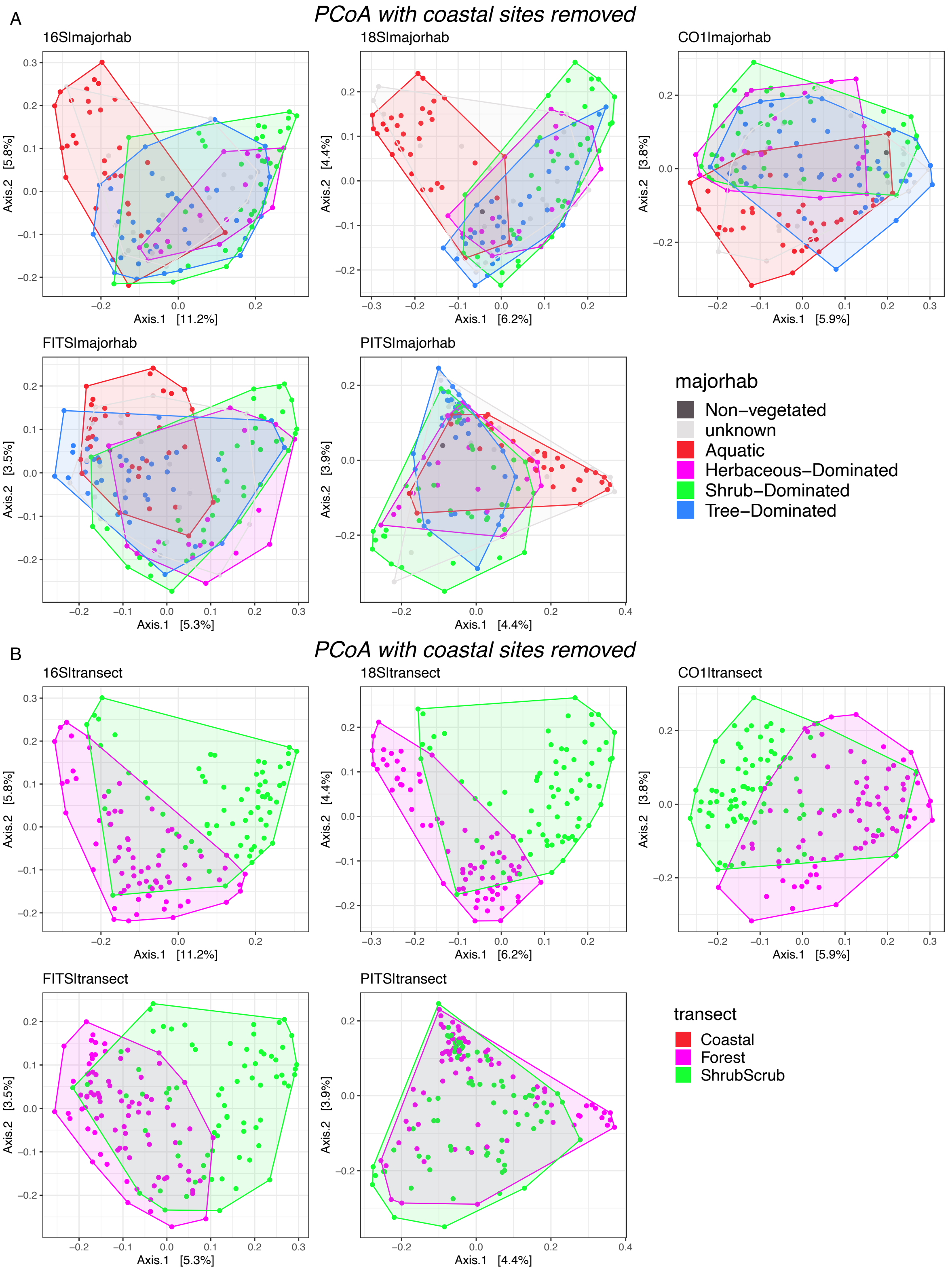

701

Figure S18 Principal Coordinate Analysis (PCoA) on beta diversity excluding coastal sites (n = 98). We recalculated the binary Jaccard dissimilarity and performed PCoA. The first two principal coordinates are plotted with percentage of variance explained included in axis label. PCoA derived from *16S, 18S, CO1, FITS*, and *PITS* metabarcodes are plotted individually. Each point represents a sample site and is colored by the (A) major habitat (B) transect it belongs to. Results of *PERMANOVA* and *beta dispersion* can be found in Table S10.3-4.

***Figure S19. Beta diversity capscale majorhab***

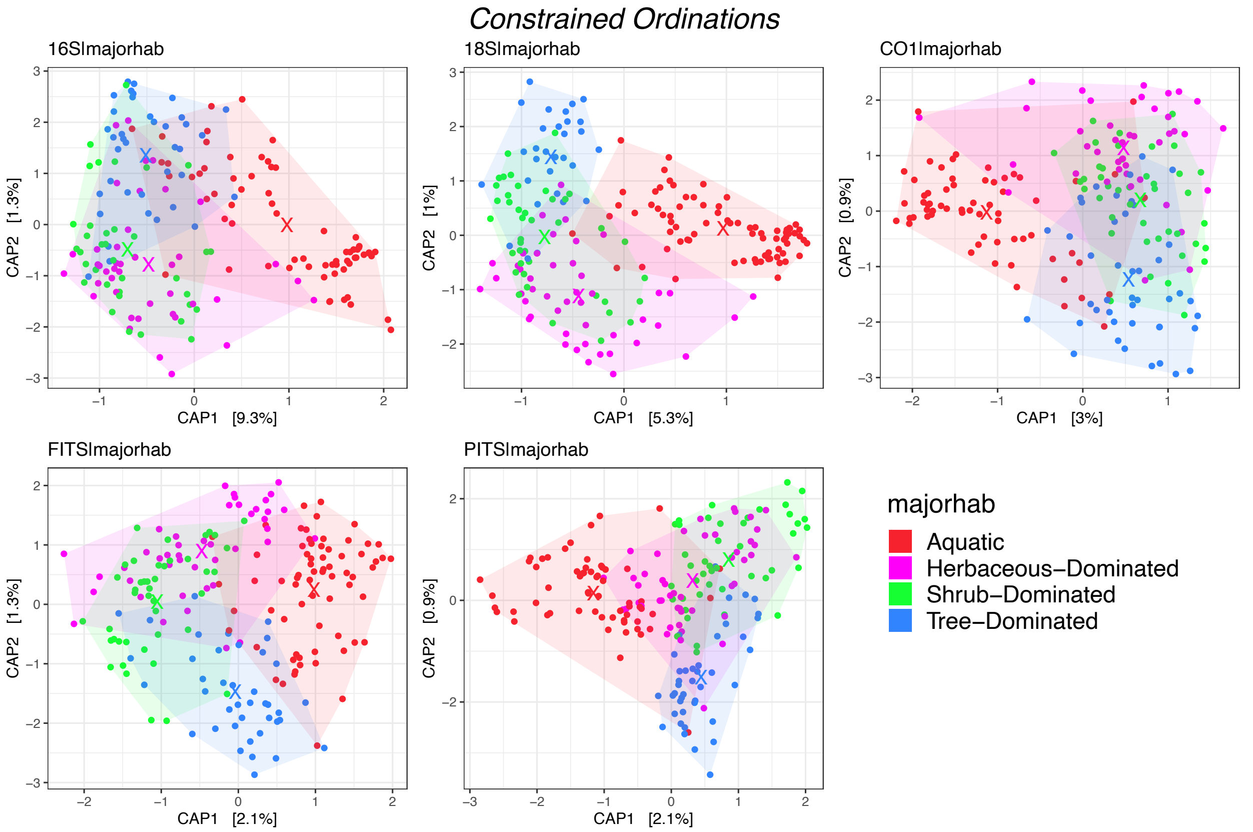

Figure S19 Partial CAP analysis of community composition by major habitat while excluding the location effect. The first two axes were plotted for each metabarcode. Each point stands for a sampling site. The “X” marks the position of each group centroids. Major habitat categories containing less than 5 samples and samples missing major habitat classifications were discarded from the analysis.

#### *Figure S20. Beta diversity: envfit*

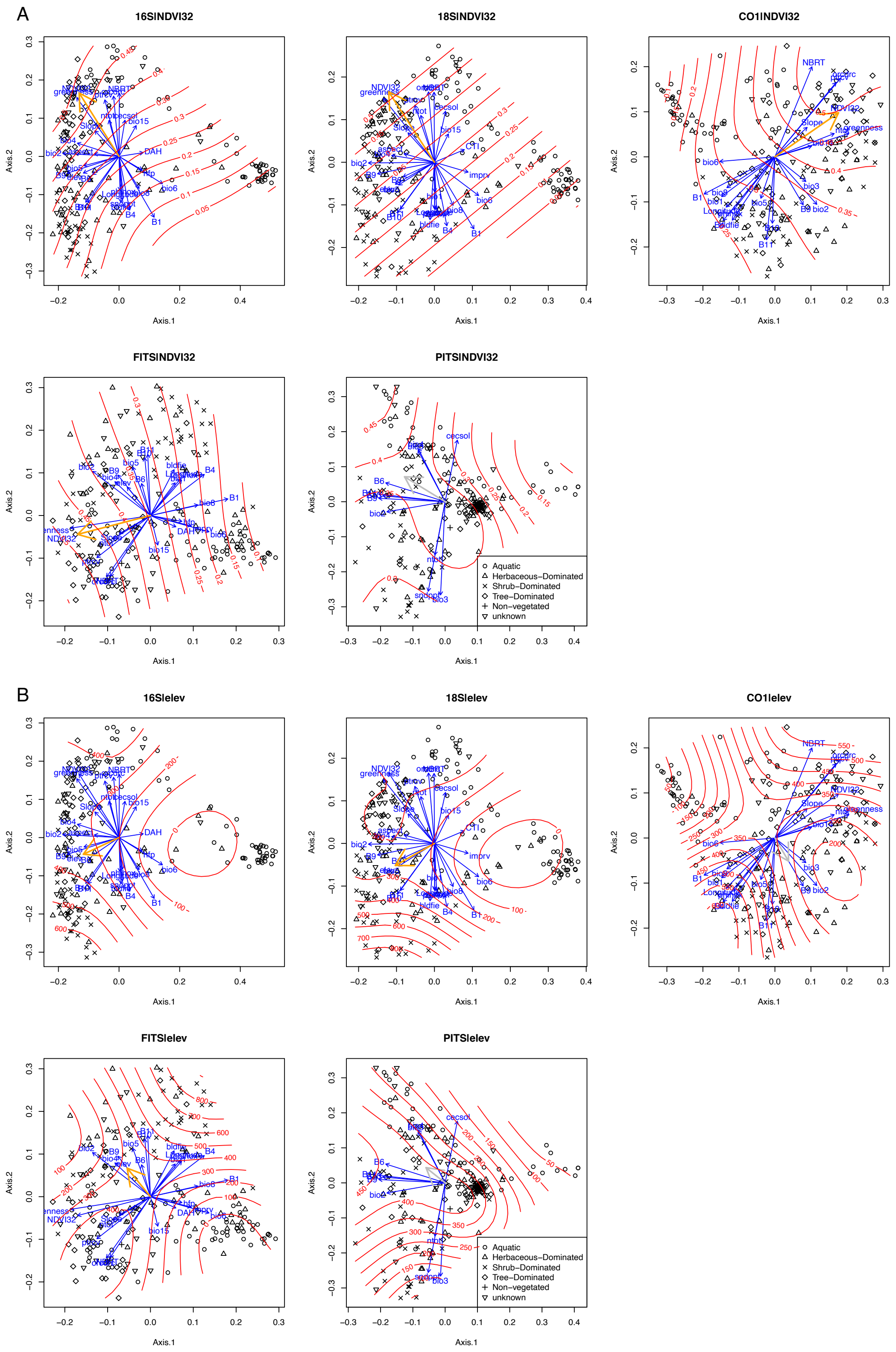

Figure S20 Gradients of environmental variables overlaid on PCoA results. Samples were plotted on the first two Principal Coordinates with different shapes of points denoting different major habitat classification. Significant variables from *envfit* output ($P < 0.001$) were overlaid as blue arrows. The direction represents the direction of the most changes and length of arrows is proportional to the strength of correlation on ordinations. *Ordisurf* output for (A) *NDVI32* and (B) *elevation* were overlaid as red contour lines with variable values denoted at each contour lines. The *envfit* arrow for the corresponding variable is bolded, with significant correlations ($P < 0.001$) colored in orange and non-significant correlations ($P < 0.001$) colored in gray.

***Figure S21. Zeta diversity***

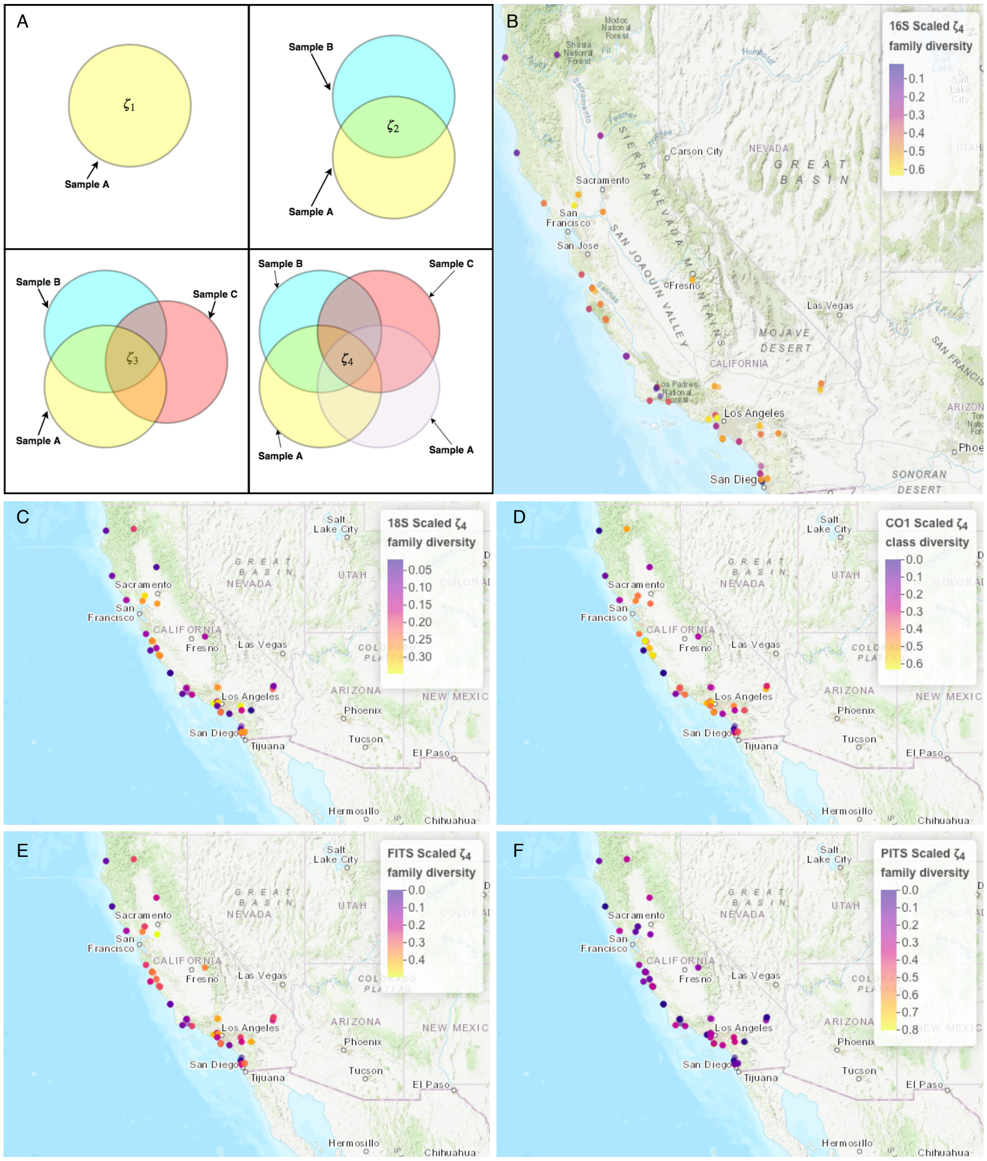

Figure S21 **Illustration and distribution of zeta diversity.** (A) An illustration of the first four orders of ζ diversity. The mean number of unique categories of organisms per sample, a measure of α diversity, is represented by the value $\zeta_1$. In comparing two or more samples the average number of unique categories held in common between any two samples is represented by the value $\zeta_2$. The mean value then of β diversity, as described by Jaccard distance, for sets of two or more samples is $\zeta_2/(2\zeta_1 - \zeta_2)$ (Simons et al. 2019). For sets of three or more samples the value of $\zeta_3$ represents all of the unique categories held in common between three samples. This process can be extended to N samples, allowing for a determination of the values $\zeta_1$ through $\zeta_N$. In this study, we set *N* as 4. (B - F) Example Scaled $\zeta_4$ diversity for the 210 communities detected with the (B) *16S*, (C)*18S*, (D) *CO1*, (E) *FITS* and (F) *PITS* metabarcode. Taxa are defined at the family level.

***Figure S22. Gradient forest permutation***

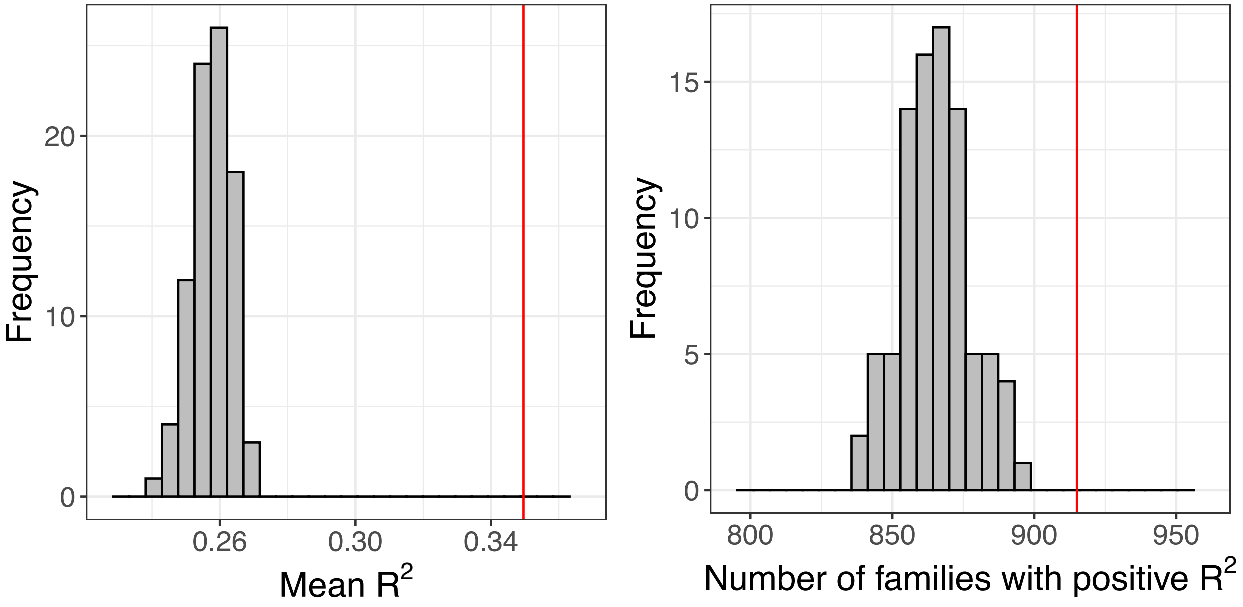

Figure S22 The distribution of $R^2$ and number of families responded with positive $R^2$ in permuted gradient forest models (N = 100). The red line shows the corresponding value in the unpermuted real model.

#### *Figure S23. Gradient forest most predictable families*

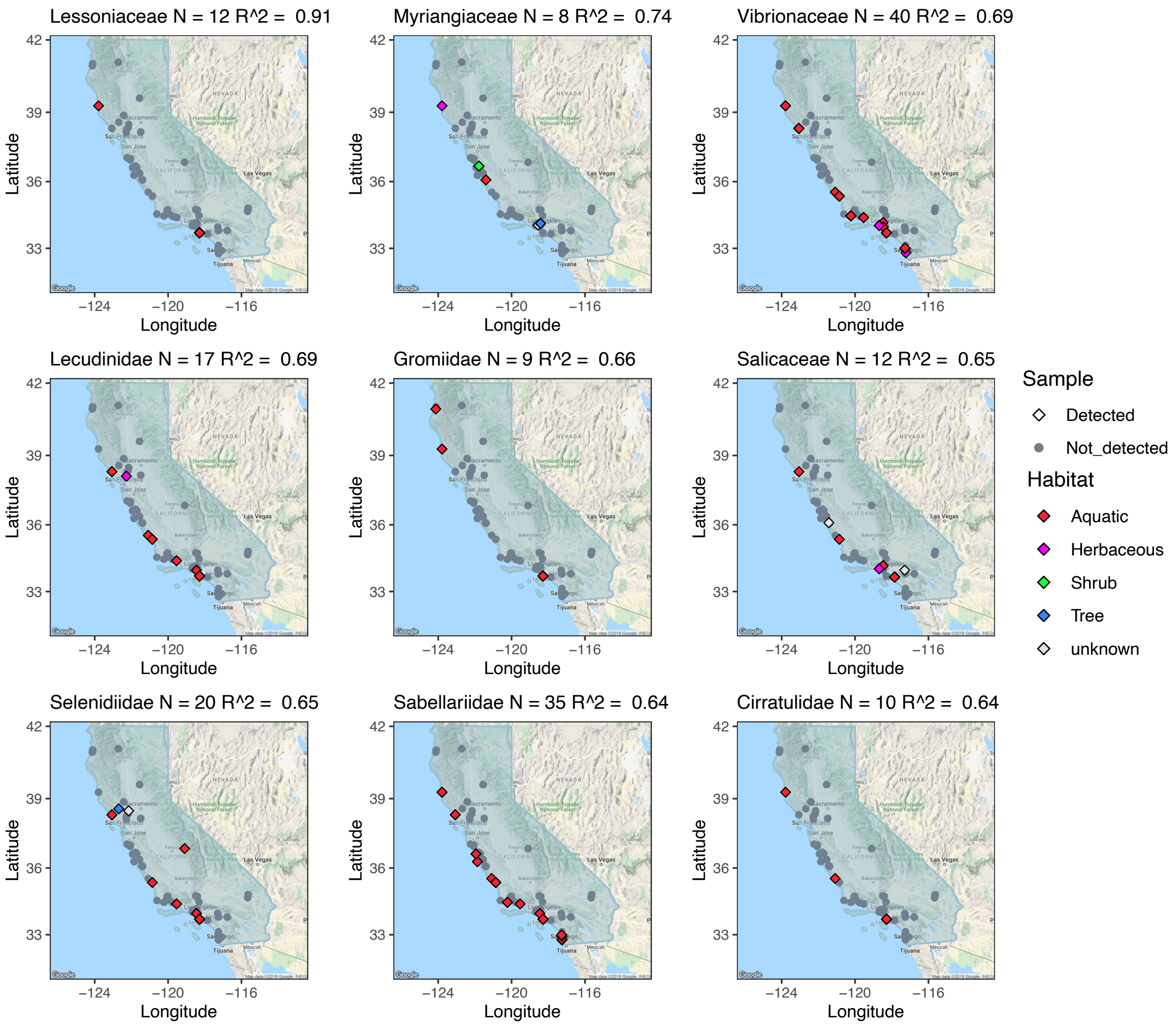

Figure S23 Map of samples containing the top 9 families most effectively modeled. Their individual $R^2$ and total number of occurrences are noted above each panel. Sites without the specified families are marked in gray points as background. Sites with occurrences are marked in diamonds and colored by the major habitat it belonged to. The blue outline shows the range of California.

***Figure S24. Gradient forest families' predictability and positive occurrences***

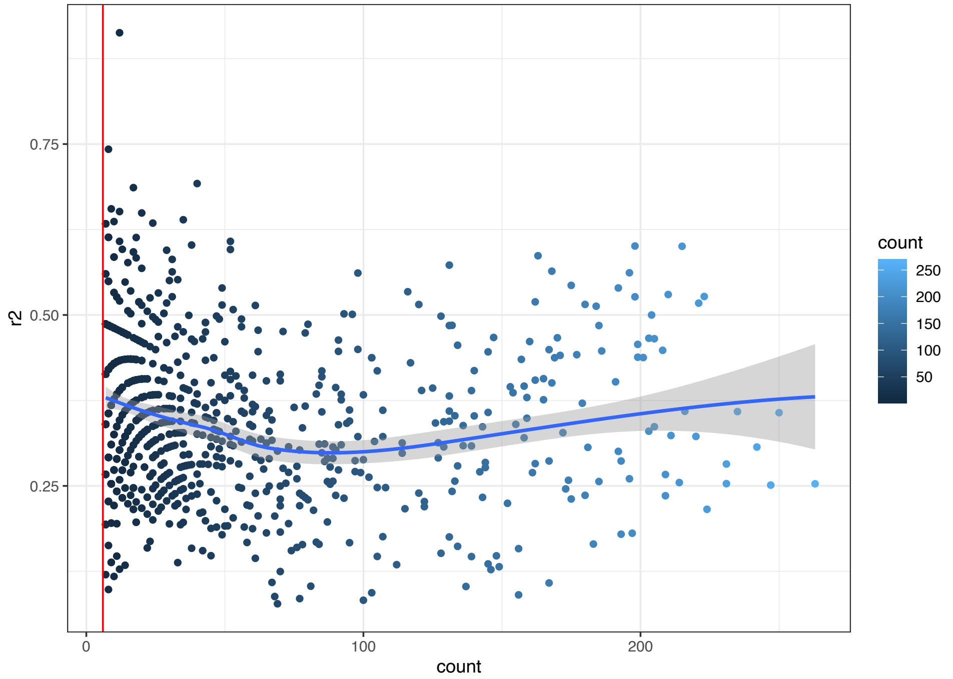

Figure S24 Individual families $R^2$ plotted against the number of positive observations in the gradient forest. The site frequency counts for each of the 915 families with positive $R^2$ were plotted. The red vertical line stands for the two-tailed 95% percentile cutoff, with a lowest observation frequency of 6. The blue line represents the best-fitted trends with gray shades denotes the 95% confidence interval (the *geom_smooth* function in R package *ggplot2*, Wickham 2016).

***Figure S25. Improvement over uninformed maps for gradient forest maps***

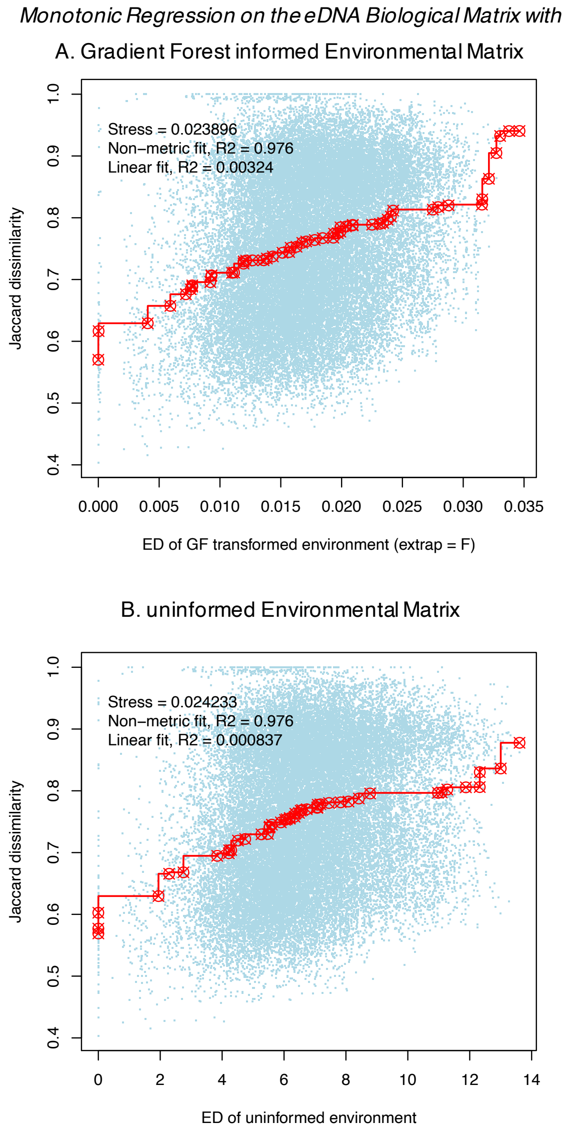

Figure S25 Stress plot derived from Monotonic regression on the Jaccard dissimilarity in biological community over (A) Euclidean distance (ED) matrix in the *gradient forest* transformed environmental space or (B) ED matrix in the uninformed environmental space. Axis labels: *ED of GF transformed environment* = Euclidean distance matrix for sites in the *gradient forest* transformed (not using extrapolation method for extreme values, *extrap = F*) environmental space; *ED of uninformed environment* = Euclidean distance matrix for sites in the uninformed environmental space; *Jaccard dissimilarity* = community ecology Jaccard dissimilarity matrix for sites in biological space calculated from the *gradient forest* input biological matrix.

***Figure S26. Gradient forest without coastal sites***

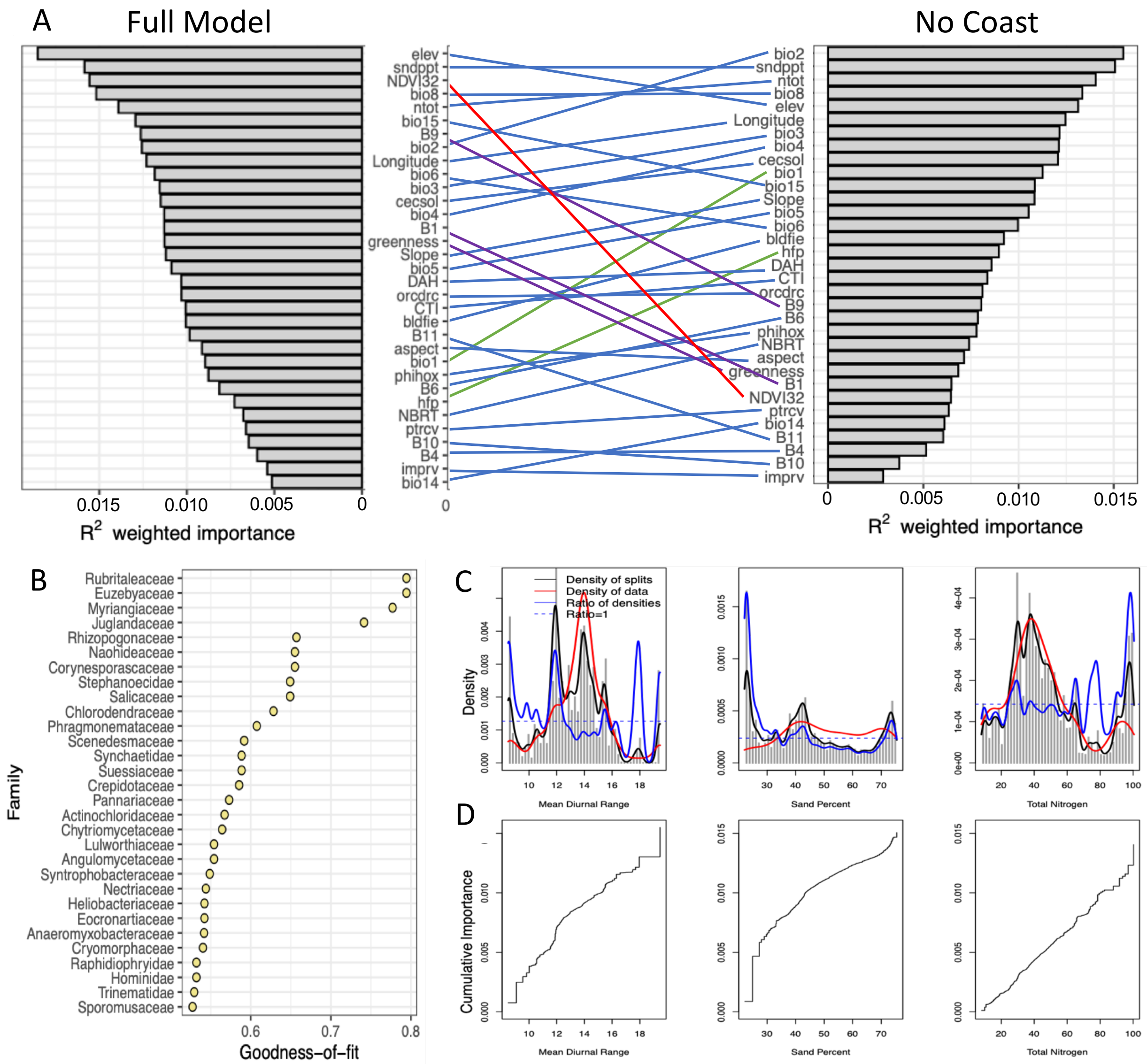

Figure S26 (A) Ranked overall importance for 33 environmental predictors in the gradient forest result with coastal samples removed versus the full model. Lines connect the same predictors across tests, with line color indicating changes in rank in the no coast model: green indicates increased importance, purple indicates decreased importance, and red indicates strong decreased importance. (B) Ranked goodness-of-fit (1 - relative error rate) for the top 30 families (response variables). (C-D) Plotted is the community turnover along the three most important environmental gradients: mean diurnal range (*bio2*), sand percentage (*sndppt*) and total nitrogen (*ntot*). (C) The gray histogram shows binned split importance at each gradient. Kernel density of splits (black lines), of observed predictor values (red lines) and of splits standardized by observation density (blue lines) are overlaid. The horizontal dashed line indicates where the ratio is 1. Each curve integrates to the importance of the predictor (A). (D) The line shows cumulative importance distributions of splits improvement scaled by $R^2$ weighted importance and standardized by density of observations, averaged over all families.
