## Supplemental Tables for "A Biodiversity Composition Map of California Derived from Environmental DNA Metabarcoding and Earth Observation": README.txt

This zip archive "Supplemental_Tables_CALeDNA_biorxiv" is part of the Supplemental Information for: 
Lin et al. “A Biodiversity Composition Map of California Derived from Environmental DNA Metabarcoding and Earth Observation” 
Uploaded to the preprint archive https://www.biorxiv.org/ on June 19, 2020

Created by: Meixi Lin

Supplemental Tables: 
•	S1_List_and_explanation_environmental_variables.xlsx
•	S2_Metabarcoding_primer_info.xlsx
•	S3_decontaminated_ASVs_five_metabarcodes.xlsx
•	S4_concordance_in_replicates.xlsx
•	S5_UCNRS_site_comparison.xlsx
•	S6_TOS_GBIF.xlsx
•	S7_kruskal_alpha_div.xlsx
•	S8_individual_lm_alpha_div.xlsx
•	S9_pls_alpha_div.xlsx
•	S10_beta_dispersion_permanova_beta_div.xlsx
•	S11_capscale_varpart_beta_div.xlsx
•	S12_envfit_beta_div.xlsx
•	S13_model_stats_decay_pattern_zeta_div.xlsx
•	S14_gradient_forest_input.xlsx
•	S15_gradient_forest_output.xlsx
•	S16_network_analysis.xlsx
